# DNA methylation modulates nucleosome retention in sperm and H3K4 methylation deposition in early mouse embryos

**DOI:** 10.1101/2024.02.06.579069

**Authors:** Grigorios Fanourgakis, Laura Gaspa-Toneu, Pavel A. Komarov, Evgeniy A. Ozonov, Sebastien A. Smallwood, Antoine H.F.M. Peters

## Abstract

DNA methylation (DNAme) serves a stable gene regulatory function in somatic cells (1). In the germ line and during early embryogenesis, however, DNAme undergoes global erasure and re-establishment to support germ cell and embryonic development (2). While *de novo* DNAme acquisition during male germ cell development is essential for setting genomic DNA methylation imprints, other intergenerational roles for paternal DNAme in defining embryonic chromatin after fertilization are unknown. To approach this question, we reduced levels of DNAme in developing male germ cells through conditional gene deletion of the *de novo* DNA methyltransferases DNMT3A and DNMT3B in undifferentiated spermatogonia. We observed that DNMT3A serves a DNAme maintenance function in undifferentiated spermatogonia while DNMT3B catalyzes *de novo* DNAme during spermatogonial differentiation. Mutant male germ cells nevertheless completed their differentiation to sperm. Failing *de novo* DNAme in *Dnmt3a*/*Dnmt3b* double deficient spermatogonia is associated with increased nucleosome occupancy in mature sperm, preferentially at sites with higher CpG content, supporting the model that DNAme modulates nucleosome retention in sperm (3). To assess the impact of altered sperm chromatin in the formation of embryonic chromatin, we measured H3K4me3 occupancy at paternal and maternal alleles in 2-cell embryos using a newly developed transposon-based tagging assay for modified chromatin. Our data show that reduced DNAme in sperm renders paternal alleles permissive for H3K4me3 establishment in early embryos, independently of possible paternal inheritance of sperm born H3K4me3. Together, this study provides first evidence that paternally inherited DNAme directs chromatin formation during early embryonic development.

## INTRODUCTION

Inheritance of genetic information from parents to offspring is fundamental to sexual reproduction. Recent studies have suggested alternative modes of inheritance that extend beyond the DNA sequence itself. Such epigenetic modes of inheritance encompass various means of transmission, such as DNA methylation (DNAme) - the presence of a methyl group at the carbon-5 position of cytosine in the CpG dinucleotide context-, post translational modifications (PTMs) on histone proteins, and noncoding RNAs, which have been implicated in the transmission of parental traits and the regulation of gene expression in subsequent generations (4,5).

Global patterns of DNAme undergo multiple waves of acquisition and resetting during the mammalian life cycle. After germ cell specification during embryogenesis, genome-wide DNAme levels are reduced in proliferating primordial germ cells (PGCs), reaching residual levels of ∼3-5% upon their arrival in embryonic gonads. DNAme is subsequently restored in a sexually dimorphic manner, with male germ cells (referred to as prospermatogonia) carrying high levels (∼80%) by the time of birth (6–9). Most global DNAme is reestablished in male embryonic germ cells by three *de novo* cytosine methyltransferases, DNMT3A, DNMT3B and DNMT3C along with their cofactor DNMT3L (10–13). Residual DNAme acquisition occurs after birth leading to highly methylated genomes in mature spermatozoa (14–16). After birth, <u>u</u>ndifferentiated <u>s</u>permato<u>g</u>onia (SgU) divide mitotically, undergo differentiation forming <u>d</u>ifferentiating <u>s</u>permato<u>g</u>onia (SgD), and subsequently undergo meiosis and spermiogenesis to form highly differentiated haploid spermatozoa. While spermatozoa contain high levels of DNAme throughout their genomes, most regions enriched in CpG dinucleotides (CpG islands, CGIs), generally serving gene regulatory functions, remain unmethylated (16).

During the late stages of spermatid development, chromatin undergoes a major reprogramming event with most nucleosomes being replaced by small basic proteins called protamines, causing extensive nuclear compaction (17). A small fraction of nucleosomes, however, is retained in sperm, up to approximately 2% in mouse and 15% in human(18–20). Several studies reported nucleosome distributions throughout the sperm genome (21,22) while we and others reported enrichment at sequences enriched in CpG dinucleotides, largely coinciding with promoters and exons of coding and non-coding genes (3,19,20,22). We further observed an inverse correlation between the degree of nucleosomal retention and the level of DNAme at such sequences (3,19,20). It is currently unknown whether DNAme serves a regulatory role in nucleosomal remodeling during spermiogenesis. Recently, previously reported nucleosome enrichments profiles of sperm chromatin were challenged to originate largely from cell free chromatin derived from somatic cellular contaminants (23). Methodological difficulties in preparing open sperm chromatin, however, preclude a decisive conclusion (23,24).

Upon fertilization, a process of extensive epigenetic reprogramming takes place, enabling the establishment of totipotency and the subsequent differentiation of distinct cell lineages. Following the nucleosomal repackaging of the paternal genome with maternally provided histones, global DNAme is promptly and actively removed (2,25,26). DNAme patterns at specific genomic regions called imprinting control regions (ICRs) are, however, maintained, driving allele specific gene regulation at later stages of development in a parent-of-origin specific manner. Such imprinted form of gene regulation is the most well-defined example of DNAme based intergenerational epigenetic inheritance (27). While maternal transmission of histone PTMs such as H3K27me3 is required for repression of maternal loci in embryos’ placentae, representing a non-canonical form of genomic imprinting (28), an instructive role of histones and associated PTMs (29) in paternal epigenetic inheritance remains to be identified. To date, reports analyzing histone perturbations in sperm and associating them with altered phenotypes or expression in embryos are primarily correlative rather than causative in nature (17,29–34). For example, spermatozoa of rats that had been exposed to toxicants during *in utero* development were reported to have altered histone retention sites that were, in part, also present in sperm of subsequent unexposed generations (30). Spermatogonial specific overexpression of the histone demethylase KDM1A (LSD1) resulted in severely impaired development of offspring. Transmission of phenotypic defects, even via non-transgenic descendant fathers, correlated with increased H3K4me3 but not H3K4me2 occupancy in sperm (29,33). Restricting folate in diet of male mice, known to modulate DNAme homeostasis (35,36), altered H3K4me3 profiles in sperm, which were correlated with altered gene expression in embryonic progeny (32). The molecular mechanism and penetrance of such possible modes of epigenetic inheritance remain, however, to be demonstrated.

Given the known antagonistic interplay between the H3K4 and DNA methylation pathways (37–40), perturbations in one chromatin pathway may impinge on other chromatin characteristics. Understanding the extent by which chromatin states in sperm influence epigenetic reprogramming in early embryos is critical to unraveling the mechanisms underlying modes of intergenerational epigenetic inheritance and its implications for gene expression regulation.

Here we investigated DNAme as a possible determinant of chromatin configuration in sperm and as driver of paternal epigenetic inheritance. By studying germ cells single and double conditionally deficient for *Dnmt3a* and/or *Dnmt3b* we identified developmental and genomic context specific contributions of each enzyme to *de novo* and maintenance DNAme during spermatogenesis. By exploiting chromatin perturbations specific to germ cells, we identified two novel roles for DNAme in sperm in (a) modulating nucleosome occupancy and H3K4me3 enrichment at CpG-rich regions in developing sperm and (b) preventing premature acquisition of H3K4me3 specifically on paternally inherited alleles in the early embryo.

## RESULTS

### Germ line *Dnmt3a/Dnmt3b* deficiency induces DNA hypomethylation in sperm

Previous studies explored the role of *de novo* DNA methyltransferases in the reestablishment of DNAme during fetal life and the generation of spermatogonial stem cells (10–13,41). Their role during adult spermatogenesis is, however, poorly understood (42,43). By immunofluorescent staining of seminiferous tubules, we detected prominent signals of DNMT3A and DNMT3B proteins in differentiating spermatogonia marked by cKIT labeling (Figure 1A), as shown previously (44). To assess the role of these proteins during adult spermatogenesis, we generated conditional deletion models of *Dnmt3a* (*3aKO*), *Dnmt3b* (*3bKO*) and double *Dnmt3a*/*Dnmt3b* (*DKO*). Excision of floxed alleles of *Dnmt3a* and/or *Dnmt3b* was driven by the improved *iCre* recombinase transgene under the control of the *Stra8* promoter, which is active in postnatal undifferentiated and differentiating (45). DNMT3A and DNMT3B proteins were undetectable in the nuclei of spermatogonia from *3aKO* and *3bKO* animals respectively (Figure 1A), and deletion of one paralogue did not affect protein expression of the other (Supplementary Figure 1A). By RNA sequencing analysis in spermatogonia, the floxed exons for both *Dnmt3a* and *Dnmt3b* genes were undetectable in the *DKO* samples (Supplementary Figures 1B, 1C). Further, by whole genome sequencing, we did not detect any reads mapping to the floxed regions of both *Dnmt3a* and *Dnmt3b* genetic loci in *DKO* sperm (Supplementary Figures 1D). Altogether, our models show efficient conditional deficiencies of DNMT3A and/or DNMT3B allowing us to investigate their impact on DNAme and the progression of adult spermatogenesis.

**Figure 1.**
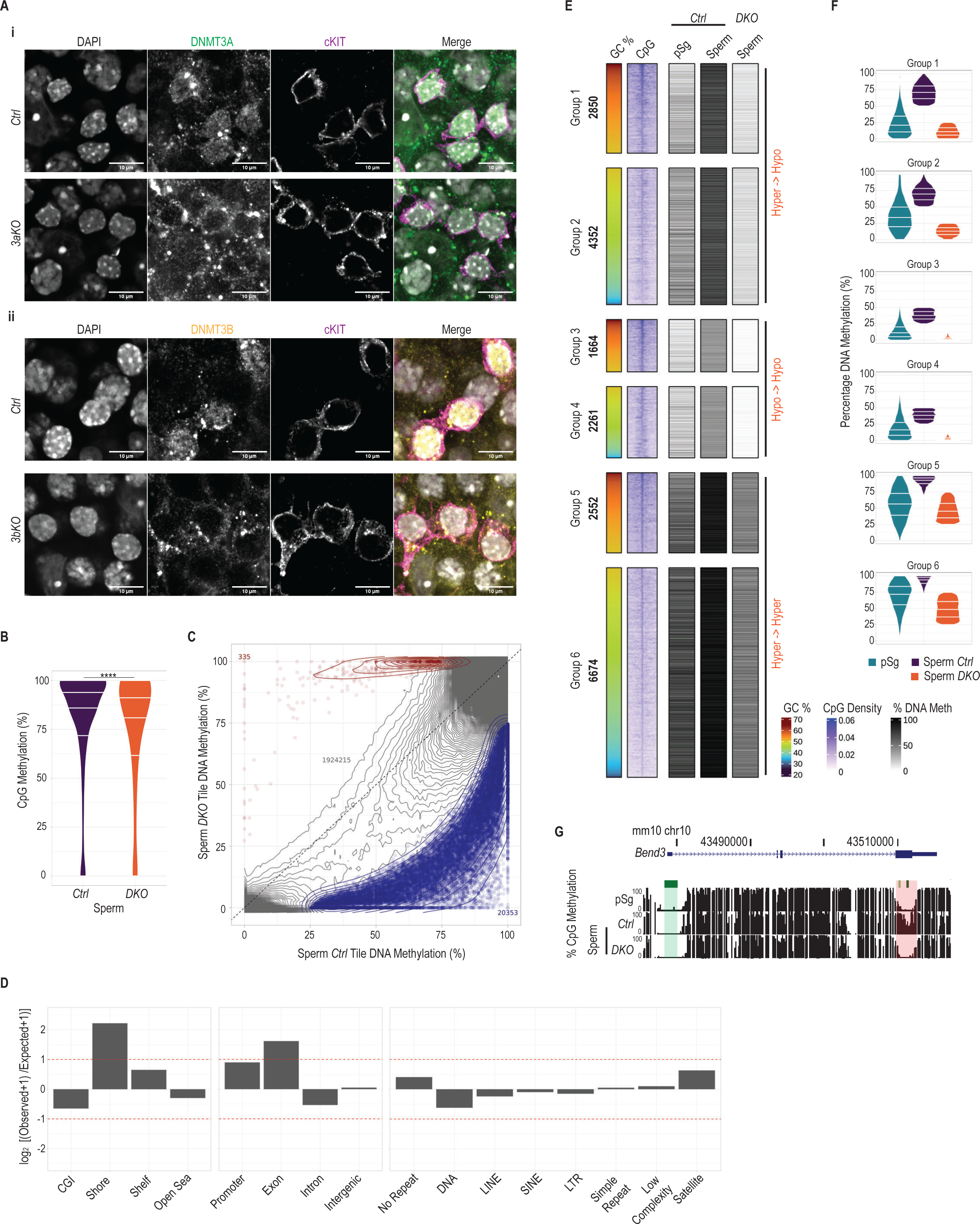
Identification of differentially hypomethylated regions in *DKO* sperm. A) Whole mount immunofluorescence staining of (i) DNMT3A in seminiferous tubules of *Ctrl* and *3aKO* testes, and (ii) DNMT3B in seminiferous tubules of *Ctrl* and *3bKO* testes. All samples were co-stained for cKIT and DNA was visualized by DAPI. Maximum projections of multiple confocal z-stacks are shown. Scale bars = 10 μm. B) Violin plot showing the percentage of DNAme of genomic CpGs assessed by EMseq in *Ctrl* (n=2) and *DKO* sperm (n=2) samples. Center lines in violin plots represent median values while upper and lower lines indicate the interquartile range (IQR; from the 25th to 75th percentile). C) Scatterplot showing percentage DNAme of 500bp genomic tiles assessed by EMseq in *Ctrl* (x-axis) and *DKO* sperm (y-axis). Tiles with significantly increased methylation in *DKO* sperm are highlighted in red (HyperDMRs) and those with reduced levels in blue (HypoDMRs). Contour lines denote the density of NonDMR tiles. D) Bar plots showing log2 enrichments of features related to CpG Islands, genes and repetitive elements among hypoDMRs tiles compared to the whole genome. P-values were calculated by Fisher’s exact test. E) Heatmap showing classification of hypoDMRs based on DNAme percentages assessed by EMseq in *Ctrl* (n=2) and *DKO* (n=2) sperm samples, and the GC percentage within hypoDMRs. pSg (n=2) DNAme percentage assessed by PBATseq (Shirane et al., 2020) is shown as an external early germ cell developmental stage reference. The density of CpGs at hypoDMRs (±5 kb from their center) is indicated. Within each group, hypoDMRs are ordered according to decreasing GC percentages. F) Violin plots showing percentages of DNAme assessed by EMseq of hypoDMRs in pSg, *Ctrl* and *DKO* sperm for each group. Center lines in violin plots represent median values while upper and lower lines indicate the interquartile range (IQR; from the 25th to 75th percentile). G) UCSC genome browser snapshot of the Bend3 locus, showing tracks for DNAme levels in pSg, *Ctrl* and *DKO* sperm. The promoter CGI and the hypoDMR are highlighted in green and red respectively.

Histological analysis of testicular sections showed an overall normal tissue morphology and presence of male germ cells at all developmental stages upon conditional ablation of the DNMT3A and/or DNMT3B proteins (Supplementary Figure 1E). We observed ∼15%-25% reduction in testicular weight upon removal of either DNMT3B or DNMT3A enzyme respectively which was further aggravated to ∼30% upon depletion of both enzymes (Supplementary Figure 1F). Nevertheless, *DKO* males sired similar numbers of live offspring (Supplementary Figure 1G) without evident phenotypic abnormalities or health issues until adulthood as compared to *Ctrl* males. These data suggest that DNMT3A/DNMT3B enzymes are largely dispensable in postnatal germ cells for the completion of spermatogenesis and production of competent gametes.

To investigate the impact of *Dnmt3a*/*Dnmt3b* deficiency on DNAme status of gametes, we performed whole genome Enzymatic Methyl-seq (EM-seq) (46) using FACS sorted sperm from 2 *Ctrl* and 2 *DKO* animals. We assayed the DNAme status of 98.6% and 99.0% of genomic CpGs to a mean combined coverage of 6.0X and 6.5X for the *Ctrl* and *DKO* samples respectively, with replicates exhibiting high correlations to each other (Supplementary Figure 2A). Compared to *Ctrl* sperm, the mean CpG methylation level in *DKO* sperm was reduced from 81.7% to 76.5% (Figure 1B). Reduced methylation was measured at CpGs residing in multiple genomic contexts (Supplementary Figures 2B, 2C). DNAme at paternally methylated imprinting control regions (ICRs) was, however, unaffected (Supplementary Figure 2D). These results contrast with those of previous studies showing extremely low global DNAme levels, including paternally methylated ICRs, upon deleting *de novo* DNA methyltransferases in fetal germ cells (10,13) or DNAme maintenance components in postnatal germ cells (42).

We calculated DNAme levels in 1’944’903 500bp non-overlapping genomic tiles which contained at least 5 CpGs and were covered by minimally 25 sequencing reads. When performing differential methylation analysis with stringent thresholds (methylation difference > 25% and FDR < 0.05), we identified 1% of all examined regions (n=20’688/N=1’944’903) as significantly differentially methylated regions (DMRs) in *DKO* versus *Ctrl* sperm, with the vast majority (20’353; 98.3%) being hypomethylated (hypoDMRs) (Figure 1C). Tiles in *DKO* sperm showed a wide range in the reduction of DNAme (Figure 1C). We found that CGI shores and exonic regions were overrepresented among hypoDMRs, while actual CGIs, intronic and repetitive sequences were represented proportionally to all genomic windows examined in this analysis (Figure 1D). In summary, these data indicate that DNMT3A/DNMT3B activities regulate DNAme dynamics at specific genomic elements during adult spermatogenesis.

To enable further mechanistic dissection of the variable hypomethylation observed in *DKO* sperm, we first classified the hypoDMRs into 6 groups based on their sequence composition (GC percentage) and their DNAme status in *Ctrl* and *DKO* samples (Figure 1E). In addition, we compared the sperm methylomes to those of *wild-type* prospermatogonia (pSg) at postnatal day 1 (PBAT-seq data from (47)), representing the starting stage of postnatal germ cell development. Group 1 & 2 hypoDMRs displayed low-to-mid levels (∼10%-50%) of DNAme in pSg which increased to high levels (>50%) in *Ctrl* sperm. In *DKO* sperm, DNAme levels were reduced below levels present in pSg (<20%), as exemplified for the *Bend3* locus, suggesting that these are bona fide *de novo* methylated regions during adult spermatogenesis (Figures 1E, 1F, 1G). HypoDMRs in groups 3 & 4 displayed only low DNAme (<20%) in pSg, mid-levels of DNAme (25%-50%) in *Ctrl* sperm and extremely low DNAme (<5%) levels in *DKO* sperm, suggesting that DNMT3A/DNMT3B enzymes are also required to sustain DNAme and/or counteract demethylation activities at these regions (Figures 1E, 1F). Finally, groups 5 & 6 display mid-to-high DNAme in pSg (∼30%-80%) and *Ctrl* sperm (75%-100%). However, despite a significant decrease in DNAme in *DKO* sperm, these regions maintained mid-levels of DNAme (25%-75%), suggesting that DNMT3A/DNMT3B confer *de novo* activity, while other DNA methyltransferases (such as *Dnmt1* which is highly expressed in spermatogonia) may act on these regions as well (Figures 1E, 1F, Supplementary Figure 3D). In general, over ∼65% of GC-rich regions (groups 1, 3 & 5) are located within a genic feature and over 40% of them reside in a vicinity of a CpG island (Supplementary Figures 2E, 2F). In summary, these results identify a novel function for the DNMT3A and/or DNMT3B enzymes in *de novo* depositing and preserving DNAme during spermatogenesis.

### *Dnmt3a* and *Dnmt3b* control *de novo* DNAme in spermatogonial stem cells and differentiated cells respectively

We then set out to identify the enzymes responsible for the developmental dynamics of *de novo* DNAme acquisition during spermatogenesis. We FACS isolated spermatogonia and sperm from *Ctrl*, *3aKO*, *3bKO* and *DKO* animals based on DNA content and expression of cell surface markers E-cadherin (CDH1/ CD324), alpha6-integrin (ITGA6/ CD49f) and stem cell growth factor receptor (cKIT/CD117) (48–50) (Supplementary Figure 3A). Flow cytometry analysis showed that the relative abundances of <u>u</u>ndifferentiated <u>s</u>permato<u>g</u>onia (SgU), expressing CDH1^high^, ITGA6^high^, cKIT^low^, and of <u>d</u>ifferentiated <u>s</u>permato<u>g</u>onia (SgD), expressing CDH1^low^, ITGA6^low^, cKIT^high^, were similar between *Ctrl* and *DKO* animals (Supplementary Figures 3B). Immunofluorescence staining of sorted cells recapitulated the histological findings, with higher expression of DNMT3A and DNMT3B detected in SgD compared to SgU (Supplementary Figure 3C). RNA-seq analysis confirmed the purity of samples based on expression of general and differentiation specific germ cell markers and the absence of somatic cell markers from the FACS sorted germ cells (Supplementary Figures 3D, 3E, 3F). Interestingly, RNA-seq analysis identified hardly any significantly differentially expressed genes between *DKO* and *Ctrl* spermatogonia (Supplementary Figures 3G, 3H), corroborating that loss of DNMT3A/DNMT3B expression does not majorly affect spermatogonial development.

We next performed Reduced Representation Bisulfite Sequencing (RRBS) using DNA extracted from the FACS isolated spermatogonial populations (SgU and SgD) and mature sperm from *Ctrl*, *3aKO*, *3bKO* and *DKO* animals. We assayed DNAme from two biological replicates for each sample for 12.4% to 17.3% of genomic CpGs to a mean combined coverage ranging from 23.4X to 32.1X, which showed high correlation to each other (Supplementary Figure 4A). Then we calculated DNAme levels in 214’798 500bp non-overlapping genomic tiles which contained at least 5 CpGs and were covered by minimally 20 sequencing reads and performed differential methylation analysis with stringent thresholds (methylation difference > 20% and FDR < 0.05). Due to the lower genomic coverage of the RRBS experiment we detected 1’987 of 20’353 hypoDMRs (9.7%) previously identified by EM-seq (Supplementary Figure 4B). In *Ctrl* samples, hypoDMRs gained more DNAme when SgU differentiated to SgD, than during subsequent development from SgD to sperm (Figure 2A, 2E). This was particularly significant when analyzing changes at individual hypoDMRs (Supplementary Figures 4Bi). For all six groups, these DNAme dynamics during spermatogenesis were well captured within the first two dimensions of a principal component analysis (PCA) (Supplementary Figure 5A). We conclude that postnatally, most *de novo* DNAme occurs during differentiation of spermatogonia, prior to their entry into meiosis and haploid differentiation (7). Consistently, deletion of both *Dnmt3a* and *Dnmt3b (DKO)* resulted in considerable reduction of DNAme in all cell types examined (Supplementary Figure 4Bii, Supplementary Figures 5A, 5B), abolishing any gains in DNAme during spermatogenesis (Figure 2B, 2E). Accordingly, the PCA showed a high similarity between the methylomes of all *DKO* germ cell types while they differed majorly from those of *Ctrl* samples (Supplementary Figure 5A).

**Figure 2.**
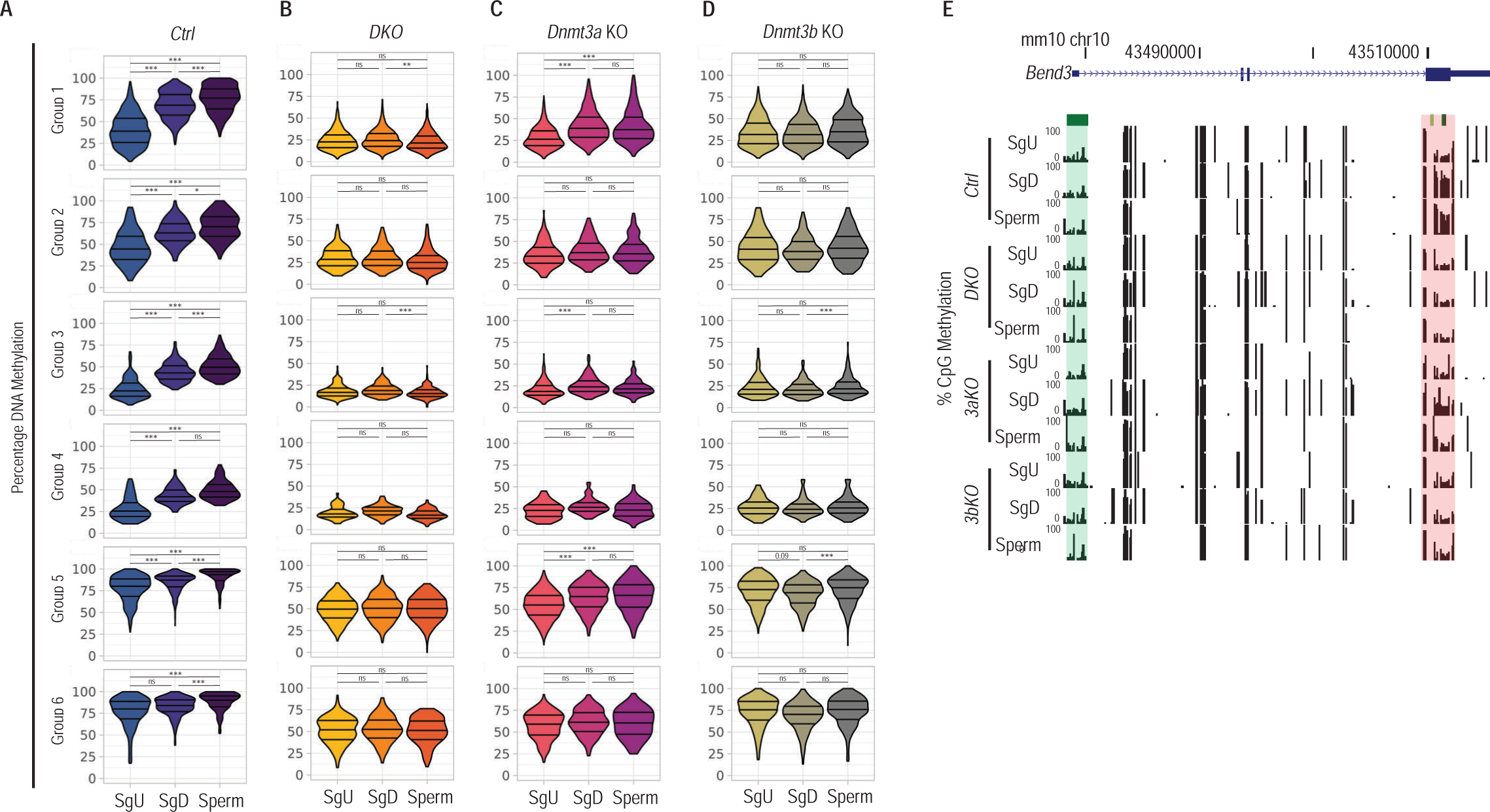
DNAme dynamics of hypoDMRs during spermatogonial differentiation upon Dnmt3a and/or Dnmt3b deletion. A) Violin plots showing percentages DNAme of hypoDMRs assessed by RRBS in *Ctrl* undifferentiated spermatogonia (SgU) (n=2), *Ctrl* differentiated spermatogonia (SgD) (n=2) and *Ctrl* sperm (n=2) samples. Two-sample Wilcoxon tests were performed between indicated groups * P < 0.05, ** P ≤ 0.01, *** P ≤ 0.001. B) Violin plots showing percentages DNAme of hypoDMRs assessed by RRBS in *DKO* undifferentiated spermatogonia (SgU) (n=2), *DKO* differentiated spermatogonia (SgD) (n=2) and *DKO* sperm (n=2) samples. Two-sample Wilcoxon tests were performed between indicated groups * P < 0.05, ** P ≤ 0.01, *** P ≤ 0.001. C) Violin plots showing percentages DNAme of hypoDMRs assessed by RRBS in *3aKO* undifferentiated spermatogonia (SgU) (n=2), *3aKO* differentiated spermatogonia (SgD) (n=2) and *3aKO* sperm (n=2) samples. Two-sample Wilcoxon tests were performed between indicated groups * P < 0.05, ** P ≤ 0.01, *** P ≤ 0.001. D) Violin plots showing percentages DNAme of hypoDMRs assessed by RRBS in *3bKO* undifferentiated spermatogonia (SgU) (n=2), *3bKO* differentiated spermatogonia (SgD) (n=2) and *3bKO* sperm (n=2) samples. Two-sample Wilcoxon tests were performed between indicated groups * P < 0.05, ** P ≤ 0.01, *** P ≤ 0.001. E) UCSC genome browser snapshot of the Bend3 locus, showing tracks for DNAme levels in SgU, SgD and sperm in *Ctrl*, *3aKO*, *3bKO* and *DKO* samples. The promoter CGI and the hypoDMR are highlighted in green and red respectively.

To identify enzyme specific roles of DNMT3A and DNMT3B during spermatogenesis, we studied methylome changes in single mutants. Intriguingly, we measured gains of DNAme in *3aKO* SgD and sperm relative to *3aKO* SgU particularly obvious in GC-rich groups 1, 3 & 5 (Figures 2C, 2E), while *Dnmt3b* depleted SgU, SgD and sperm show comparable levels of DNAme (Figure 2D, 2E). Indeed, PC analysis showed that the DNAme states in all three *3bKO* cell types were similar to each other, while *3aKO* SgU methylome differed from *3aKO* SgD and sperm methylomes (Supplementary Figure 5A). These data argue that *de novo* DNAme acquisition during spermatogonial differentiation is primarily catalyzed by DNMT3B rather than DNMT3A.

In addition, we noticed that *Dnmt3a* depletion led to decreased DNAme at many hypoDMRs in SgU, particularly obvious in groups 1, 2, 5 & 6, which display intermediate to high DNAme in *Ctrl* cells (Supplementary Figures 4Biii and 5Bi). In contrast, *Dnmt3b* depletion in SgU lead to a marginal decrease of DNAme in groups 1 and 5 (Supplementary Figure 5Bi), with only a minor fraction of hypoDMRs displaying statistically significant reduction of DNAme (Supplementary Figure 4Biv). Accordingly, the DNAme status of *3aKO* SgU was largely similar to *DKO* SgU in the PCA, while the methylome of *3bKO* SgU was mostly similar to that of *Ctrl* SgU (Supplementary Figure 5A). Hence, these data show that DNMT3A serves a more prominent role than DNMT3B in depositing DNAme in SgU cells, thereby maintaining DNAme levels set prior to SgU formation.

Nonetheless, a comparison of methylomes between single and double mutant samples supports the notion that the two enzymes serve some overlapping functions during spermatogenesis (Supplementary Figures 4Biii, 4Biv, 5Bii, 5Biii). In summary, during postnatal spermatogenesis we identified a major but not exclusive function of DNMT3A to establish and/or maintain basal DNAme levels at hypoDMRs in SgU, while DNMT3B mostly catalyzes *de novo* DNAme at hypoDMRs in response to differentiation cues in SgD.

### Stable H3K36me3 marking around hypoDMRs in *Ctrl* and *DKO* spermatogonia

Next, we investigated possible mechanisms of DNMT3A/DNMT3B recruitment to chromatin. Previous studies showed that both enzymes are recruited to chromatin by their PWWP domains interacting with nucleosomes that have been di-or trimethylated on H3K36 by the histone methyltransferases NSD1 and NSD2 or SETD2, respectively (47,51–54). H3K36me3 is co-transcriptionally deposited within transcribed genes by SETD2 interacting with RNA polymerase II (55,56). Approximately 8% and 40% of hypoDMRs reside close to promoters or within genes respectively, that are abundantly expressed in SgU and SgD cells (Supplementary Figure 6A). These genes serve basic cellular or developmental processes rather than spermatogenesis-specific functions (Supplementary Figure 6B). To investigate a possible role of H3K36 methylation in DNMT3A/DNMT3B recruitment, we generated genome wide profiles of H3K36me3 by ultra-low input native ChIP-seq (ULI-NChIP-seq) (57) for 2 biological replicates of SgU and SgD cells from *Ctrl* and *DKO* animals (Supplementary Figure 7A). In *Ctrl* samples, H3K36me3 occupancy levels increased or got reduced during spermatogonial differentiation at a comparable number of genomic tiles. HypoDMRs behaved similarly as other genomic tiles (Supplementary Figures 7Bi, 7Ci). In SgU and SgD *DKO* samples, 2-fold more regions gained H3K36me3 signal compared to *Ctrl* samples (Supplementary Figures 7Bii, 7Biii). HypoDMRs behaved, however, similarly as nonDMR regions (Supplementary Figures 7Cii, 7Ciii).

Irrespective of the developmental stage or *Dnmt3a*/*Dnmt3b* pro- or deficiency, hypoDMRs were generally embedded in regions with moderate H3K36me3 occupancy, as compared to intragenic regions of genes characterized by high to low H3K36me3 levels, matching their high to low expression states (Supplementary Figure 8A, 8B, 8C, 8D). Contrasting typical CpG-rich promoters with extensive H3K36me3 depletion, we measured only a minor depletion of H3K36me3 at the centers of hypoDMR regions in GC-rich groups 1 and 3 which exhibit low DNAme in *Ctrl* SgU. GC-rich group 5 hypoDMRs characterized by higher DNAme in *Ctrl* SgU showed an even more uniform H3K36me3 signal (Supplementary Figures 8E, 8F). We conclude that H3K36me3 at and around hypoDMRs may suffice for DNMT3A/DNMT3B recruitment in SgU and SgD cells. It is, however, unlikely to specify DNAme gain during spermatogonial differentiation.

### HypoDMRs convert from H3K4me3 to DNAme positive states during spermatogonial differentiation

Contrary to H3K36 methylation, H3K4me3 is known to inhibit DNMT3 dependent catalysis DNAme by blocking the interaction between DNMT3A’s ADD domain with unmodified H3 N-termini (37–40). To identify possible roles of H3K4me3 in hypoDMRs, we generated genome wide profiles of H3K4me3 by ULI-NChIP-seq (57) for 2 biological replicates of SgU and SgD cells from *Ctrl* and *DKO* animals (Supplementary Figure 9A). In *Ctrl* SgU, hypoDMRs of mainly groups 1-4 displayed moderate H3K4me3 levels, comparable to levels of non-CGI promoters (Figures 3A, 3B, 3F, Supplementary Figures 10A, 10B, 10C). These hypoDMRs had low to intermediate levels of DNAme in *Ctrl* samples (Figure 2A). In contrast, hypoDMRs with high DNAme levels, belonging to groups 5 & 6, were devoid of H3K4me3 in *Ctrl* SgU (Figures 3A, 3B). Differential occupancy analysis showed a global reduction of H3K4me3 in *Ctrl* SgD relative to *Ctrl* SgU (Supplementary Figure 9Bi). Statistical testing demonstrated that GC-rich hypoDMRs showed the highest enrichment among those that had lost H3K4me3 in *Ctrl* SgD versus *Ctrl* SgU (Figure 3C). More specifically, during the transition from SgU to SgD, H3K4me3 levels became reduced at hypoDMRs gaining DNAme (Figures 3A, 3B, 3F, Supplementary Figures 10D), suggesting that H3K4me3 is removed during spermatogonial differentiation prior to DNAme acquisition.

**Figure 3.**
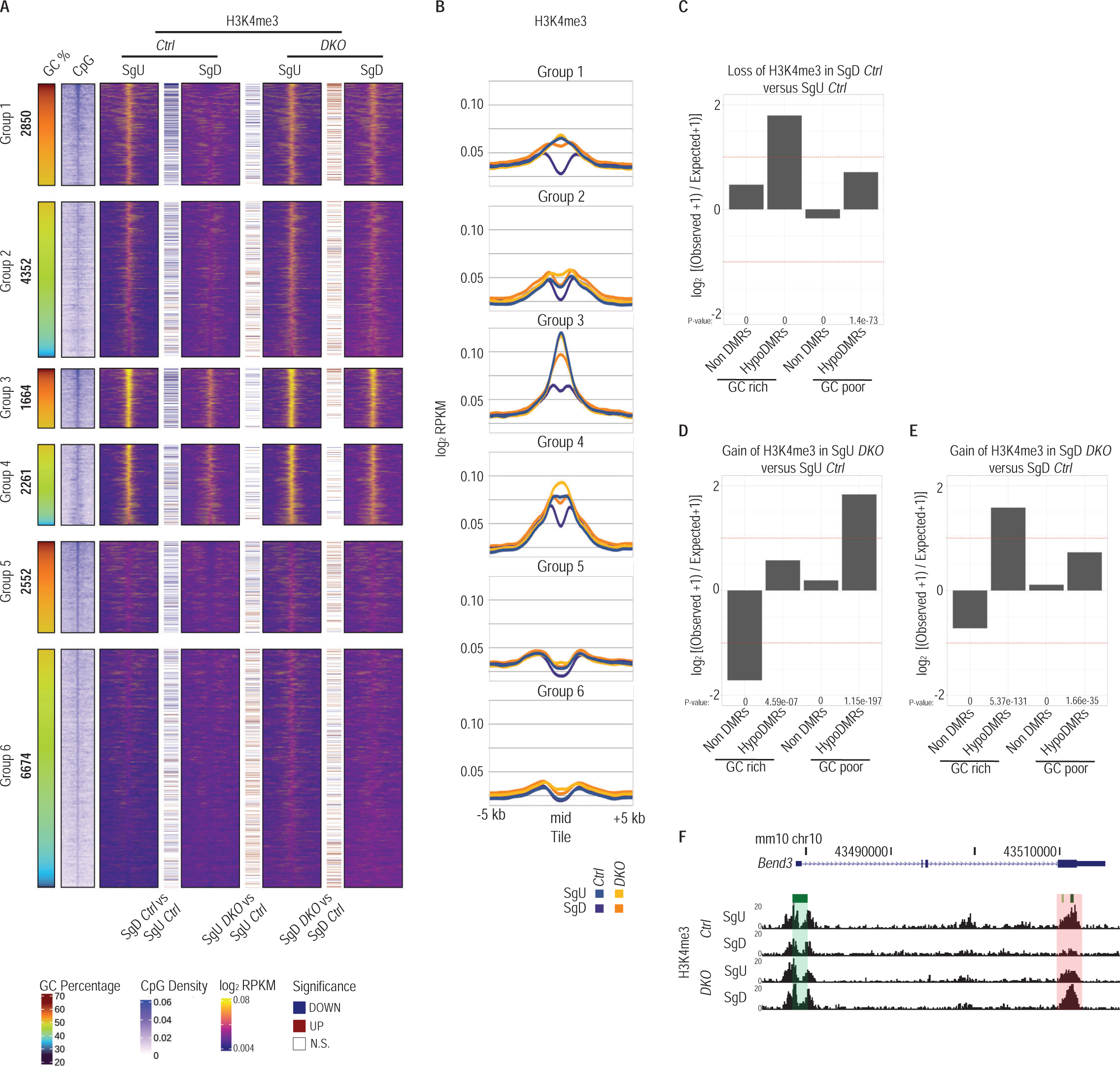
hypoDMRs in *DKO* spermatogonia retain H3K4me3. A) Heatmaps showing H3K4me3 signal at the center of hypoDMRs (±5kb) assessed by ULI-NChIP-seq in *Ctrl* SgU (n=2), *Ctrl* SgD (n=2), *DKO* SgU (n=2) and *DKO* SgD (n=2). Colored lines between heatmaps denote hypoDMRs with significantly differential H3K4me3 occupancy in the indicated contrasts. The GC percentage and CpG density at hypoDMRs (±5 kb) are indicated. B) Line plots showing average H3K4me3 signal for each group of hypoDMRs (±5kb) assessed by ULI-NChIP-seq in *Ctrl* SgU (n=2), *Ctrl* SgD (n=2), *DKO* SgU (n=2) and *DKO* SgD (n=2). C) Bar plots showing log2 enrichments of GC-rich or GC-poor nonDMR and hypoDMR tiles among tiles with significant H3K4me3 loss in *Ctrl* SgD versus *Ctrl* SgU. P-values were calculated by Fisher’s exact test. D) Bar plots showing log2 enrichments of GC-rich or GC-poor nonDMR and hypoDMR tiles among tiles with significant H3K4me3 gain in *DKO* SgU versus *Ctrl* SgU. P-values were calculated by Fisher’s exact test. E) Bar plots showing log2 enrichments of GC-rich or GC-poor nonDMR and hypoDMR tiles among tiles with significant H3K4me3 gain in *DKO* SgD versus *Ctrl*. P-values were calculated by Fisher’s exact test. F) UCSC genome browser snapshot of the Bend3 locus, showing tracks for H3K4me3 occupancy in *Ctrl* and *DKO* SgU and SgD. The promoter CGI and the hypoDMR are highlighted in green and red respectively.

When comparing H3K4me3 occupancy in *DKO* versus *Ctrl* samples in SgU, we observed that GC-poor hypoDMRs were overrepresented among tiles with increased H3K4me3 in *DKO* (Figures 3D, Supplementary Figures 9Bii, 9Cii). In contrast, we observed significant enrichment of GC-rich hypoDMRs among tiles with increased H3K4me3 in *DKO* SgD (Figures 3E, Supplementary Figures 9Biii, 9Ciii). The lack of *de novo* DNAme in *DKO* SgD (relative to *Ctrl* SgD) was evidently associated with increased H3K4me3 levels especially in GC-rich groups 1 & 3 (Figure 3A, 3B, 3F, Supplementary Figure 10E). Notably, such changes in H3K4me3 occupancy during development and in response to *Dnmt3*a/*Dnmt3b* deficiency occur in absence of transcriptional misregulation (Supplementary Figure 3G, 3H). In summary, the data indicate that H3K4me3 methyltransferases (possibly in conjunction with demethylases) versus DNMT3 enzymes mutually antagonize each other in differentiating spermatogonia e.g., by inhibiting DNMT3 catalytic activity versus preventing recruitment of KMT2A, KMT2B and SETD1A/CXXC1 H3K4 methyltransferases to methylated CpG-rich sequences.

### DNAme reduces nucleosome retention at GC-rich regions during spermiogenesis

During spermatid differentiation and nuclear compaction, a small fraction of nucleosomes evades removal and replacement by protamines (17). We and others reported previously higher nucleosomal occupancy in sperm chromatin at CpG-rich sequences lacking DNAme (3,19,20). To assess whether DNAme may restrain nucleosome retention during spermiogenesis, we performed native ChIP-seq (24) with an antibody specific to nucleosomes (58) using 2 biological replicates of *Ctrl* and *DKO* sperm (Supplementary Figure 11A). In *Ctrl* sperm, low DNAme hypoDMRs of groups 3 & 4 displayed nucleosomal occupancy, with more signal detected in GC-rich group 3. In contrast, the nucleosome signal was low or absent at hypoDMRs belonging to high DNAme groups 1,2,5,6 (Figures 4A, 4B). Genome wide differential enrichment analysis showed comparable numbers of regions with increased or decreased nucleosome occupancy (Supplementary Figure 11B). However, GC-rich hypoDMRs but not nonDMRs were significantly overrepresented among tiles with increased nucleosomal occupancy in *DKO* versus *Ctrl* sperm samples (Figure 4C, Supplementary Figure 11C). Comparing *Ctrl* and *DKO* sperm, more GC-rich group 1 hypoDMRs displayed elevated nucleosome occupancy than GC-poor group 2 hypoDMRs, although both hypoDMR groups exhibiting comparable relative reductions in DNAme (Figures 4A, 4B, 4D, Supplementary Figure 11D). Smaller increases in nucleosomal occupancy were detected for GC-rich hypoDMRs in groups 3 & 5, possible due to the presence of nucleosomes at group 3 hypoDMRs already in *Ctrl* samples and due to the remaining of intermediate levels of DNAme at group 5 hypoDMRs (Figures 4A, 4B, Supplementary Figure 11D). GC-poor group 4 & 6 hypoDMRs showed similar nucleosome occupancies between genotypes. Together, these data support a model in which a high proportion of unmethylated CpG dinucleotides within more GC-rich sequences enables nucleosome retention during spermiogenesis.

**Figure 4.**
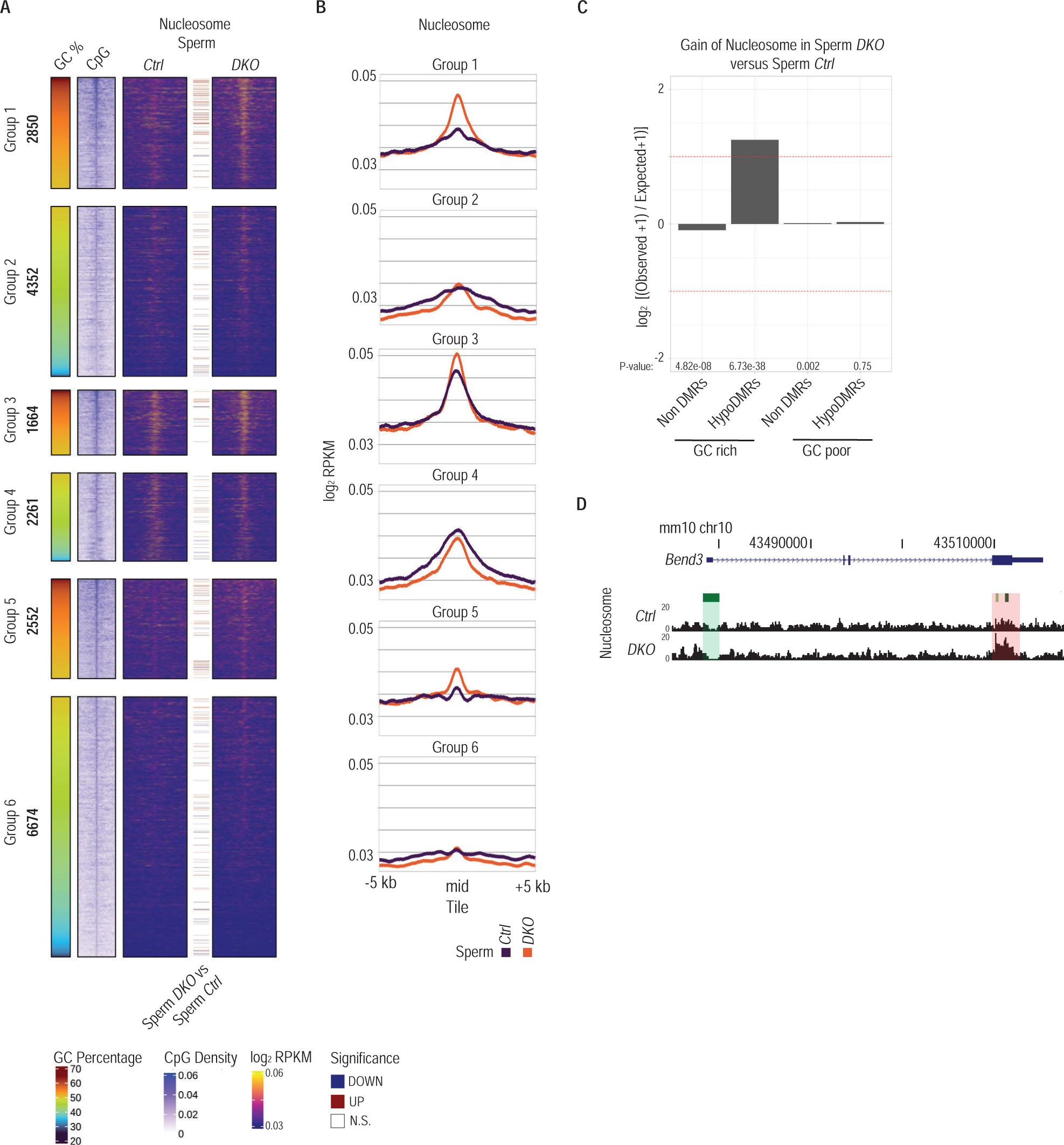
hypoDMRs gain nucleosomes in *DKO* sperm. A) Heatmaps showing nucleosome signal at the center of hypoDMRs (±5kb) assessed by ChIP-seq in *Ctrl* (n=2) and *DKO* (n=2) sperm samples. Colored lines between heatmaps denote hypoDMRs with significantly differential nucleosome occupancy between indicated samples. The GC percentage and CpG density at hypoDMRs (±5 kb) are indicated. B) Line plots showing average nucleosome signal for each group of hypoDMRs (±5kb) assessed by ULI-NChIP-seq in *Ctrl* (n=2) and *DKO* (n=2) sperm samples. C) Bar plots showing log2 enrichments of GC-rich or GC-poor nonDMR and hypoDMR tiles among tiles with significant nucleosome gain in *DKO* sperm versus *Ctrl*. P-values were calculated by Fisher’s exact test. D) UCSC genome browser snapshot of the Bend3 locus, showing tracks for nucleosome occupancy in *Ctrl* and *DKO* sperm. The promoter CGI and the hypoDMR are highlighted in green and red respectively.

### Increased H3K4me3 deposition at ectopically retained nucleosomes in *DKO* sperm

Given the antagonistic relationship between H3K4me3 and DNAme states in spermatogonia (Figure 3) and other cell types (59–61), we generated genome wide profiles of H3K4me3 by ChIP-seq (24) for 2 biological replicates of *Ctrl* and *DKO* sperm (Supplementary Figure 12A). In *Ctrl* sperm, low DNAme hypoDMRs of groups 3 & 4 with some nucleosomal occupancy (Figure 4) also displayed signal for H3K4me3. As for nucleosomal occupancy, the H3K4me3 signal was low or absent at hypoDMRs belonging to high DNAme groups 1,2,5,6 (Figures 5A, 5B). Genome wide differential enrichment analysis showed comparable numbers of regions with increased or decreased H3K4me3 occupancy (Supplementary Figure 12B). Nonetheless, GC-rich but also GC-poor hypoDMRs were significantly overrepresented among tiles with increased H3K4me3 occupancy in *DKO* versus *Ctrl* sperm samples (Figure 5C, Supplementary Figure 12C). Particularly group 1 hypoDMRs, but also group 3 and 5 hypoDMRs, displayed increased H3K4me3 enrichment in *DKO* sperm compared to *Ctrl* sperm (Figures 5A, 5B, 5D). Notably, ∼30% of regions with increased nucleosome signal displayed significantly increased H3K4me3 (Supplementary Figure 12D). These regions showed the greatest reduction of DNAme and the highest gain of nucleosome signal in *DKO* sperm (Supplementary Figures 12E, 12F). Inversely, 87% of H3K4me3 enriched regions displayed no change in nucleosome occupancy (Supplementary Figure 12D). The latter likely reflects a higher immunoprecipitation efficiency by anti-H3K4me3 over anti-nucleosome antibodies, also in line with a wider distribution of anti-H3K4me3 versus anti-nucleosome signals around the center of hypoDMRs (Figures 4A, 4B, 5A, 5B). Increased H3K4me3 occupancy measured in *DKO* sperm may result from enhanced chromatin recruitment of H3K4 methyltransferase(s) via unmethylated CpG dinucleotides (62,63) in spermatogonia.

**Figure 5.**
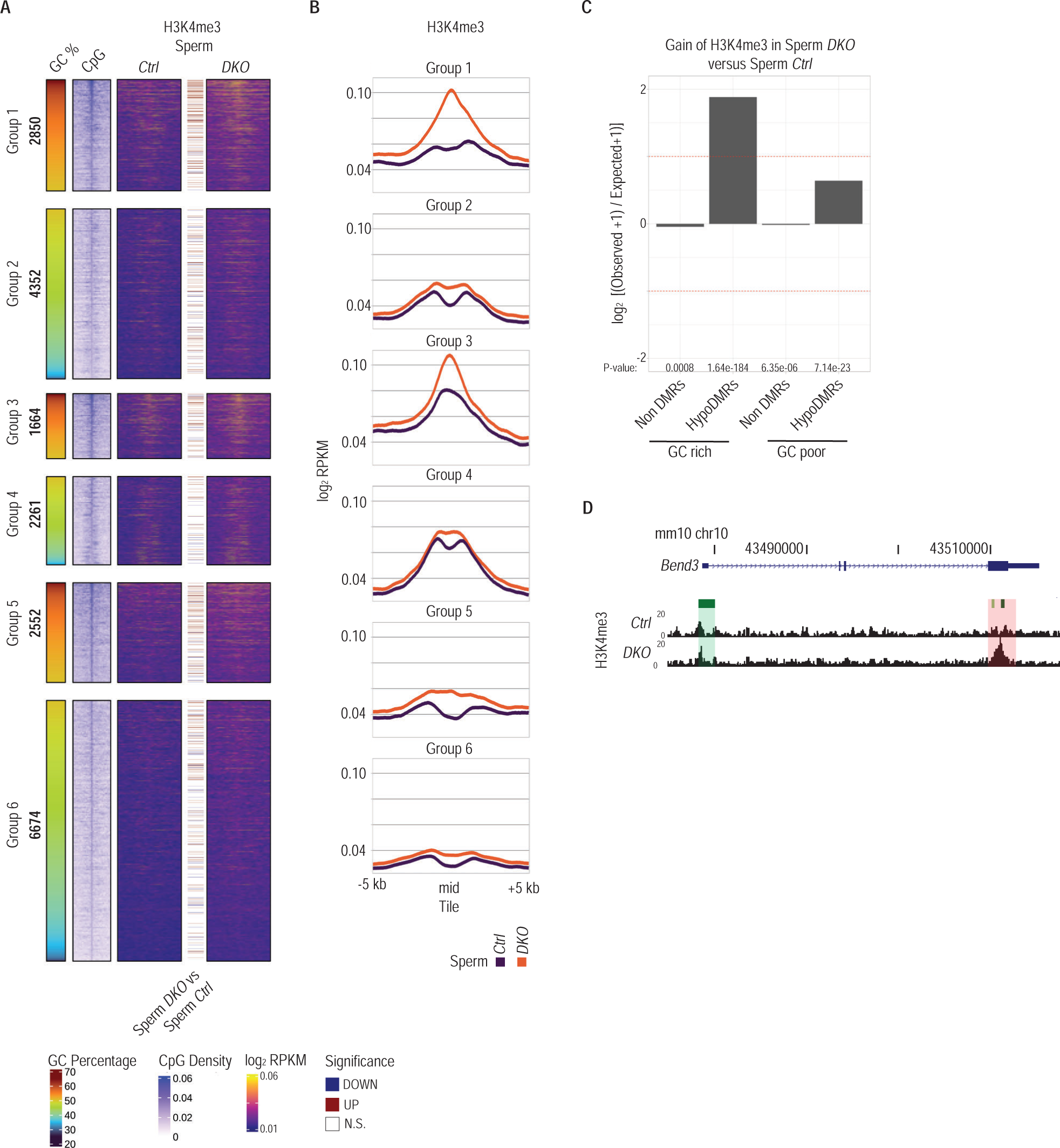
hypoDMRs gain H3K4me3 in *DKO* sperm. A) Heatmaps showing H3K4me3 signal at the center of hypoDMRs (±5kb) assessed by ChIP-seq in *Ctrl* (n=2) and *DKO* (n=2) sperm samples. Colored lines between heatmaps denote hypoDMRs with significantly differential nucleosome occupancy between indicated samples. The GC percentage and CpG density at hypoDMRs (±5 kb) are indicated. B) Line plots showing average H3K4me3 signal for each group of hypoDMRs (±5kb) assessed by ULI-NChIP-seq in *Ctrl* (n=2) and *DKO* (n=2) sperm samples. C) Bar plots showing log2 enrichments of GC-rich or GC-poor nonDMR and hypoDMR tiles among tiles with significant H3K4me3 gain in *DKO* sperm versus *Ctrl* sperm. P-values were calculated by Fisher’s exact test. D) UCSC genome browser snapshot of the Bend3 locus, showing tracks for H3K4me3 occupancy in *Ctrl* and *DKO* sperm. The promoter CGI and the hypoDMR are highlighted in green and red respectively.

### Sperm born DNAme prevents H3K4me3 deposition at hypoDMRs in mouse early embryos

To investigate whether DNAme in sperm affects chromatin establishment in embryos, shortly after fertilization, we developed a new highly sensitive chromatin profiling approach named <u>A</u>ntibody <u>TA</u>rgeted <u>T</u>agmentation <u>A</u>ssay combined with sequencing (ATATA-seq). This assay uses strategies similarly used in CUT&Tag (64) and Stacc-seq (65). It differs, however, (i) in the type of chimeric protein used, (ii) the high salt buffer conditions used during the targeting of antibody-ZZ-Tn5 transposomes to chromatin to limit the targeting to open chromatin, and (iii) the absence of washing steps. Specifically, we generated a Tn5 chimeric protein consisting of two Z domains of staphylococcal protein A fused to the Tn5 transposase (ZZ-Tn5). Next, the fusion protein is loaded with Tn5 sequencing adapters and incubated with antibodies. Finally, the “antibody-ZZ-Tn5 assembled and active transposomes” are applied to permeabilized nuclei to recognize antibody targeted regions and to insert sequencing adapters into the DNA surrounding the chromatin target sites.

We used ATATA-seq to profile H3K4me3 in triplicate pools of 50 early 2-cell stage genetically hybrid embryos, generated by fertilizing JF1/Ms background oocytes with *Ctrl* or *DKO* sperm on a C57BL/6 background allowing parental allelic discrimination based on the presence of single nucleotide polymorphisms (SNPs). The biological replicates of ATATA-seq correlate better with anti-H3K4me3 ChIP seq of 2-cell stage embryos rather than ICM datasets from Liu et al (66), validating the sensitivity of our approach (Supplementary Figure 13A). In *Ctrl* sperm derived embryos H3K4me3 occupancy was predominantly detected at hypoDMRs in groups 3 & 4 and moderately in groups 1,2,5 (Figure 6A, 6B). H3K4me3 occupancy at the maternal alleles was higher compared to the paternal alleles (Figure 6C, 6D, 6E, 6F). Genome wide differential enrichment analysis between *Ctrl* or *DKO* sperm derived embryos showed comparable numbers of regions with increased or decreased H3K4me3 occupancy, with most of the differences arising at paternal alleles rather than at maternal alleles (Supplementary Figure 13B). However, GC-rich hypoDMRs were significantly overrepresented among tiles with increased H3K4me3 occupancy in *DKO* versus *Ctrl* sperm-derived embryos (Figure 6Gi). Importantly, when splitting the maternal and paternal alleles similar enrichment was observed for nonDMRs among tiles with increased H3K4me3, however more than 2 fold enrichment of GC-rich hypoDMRs was evident only for the paternal allele (Figure 6Gii, 6Giii, Supplementary Figure 13C).

**Figure 6.**
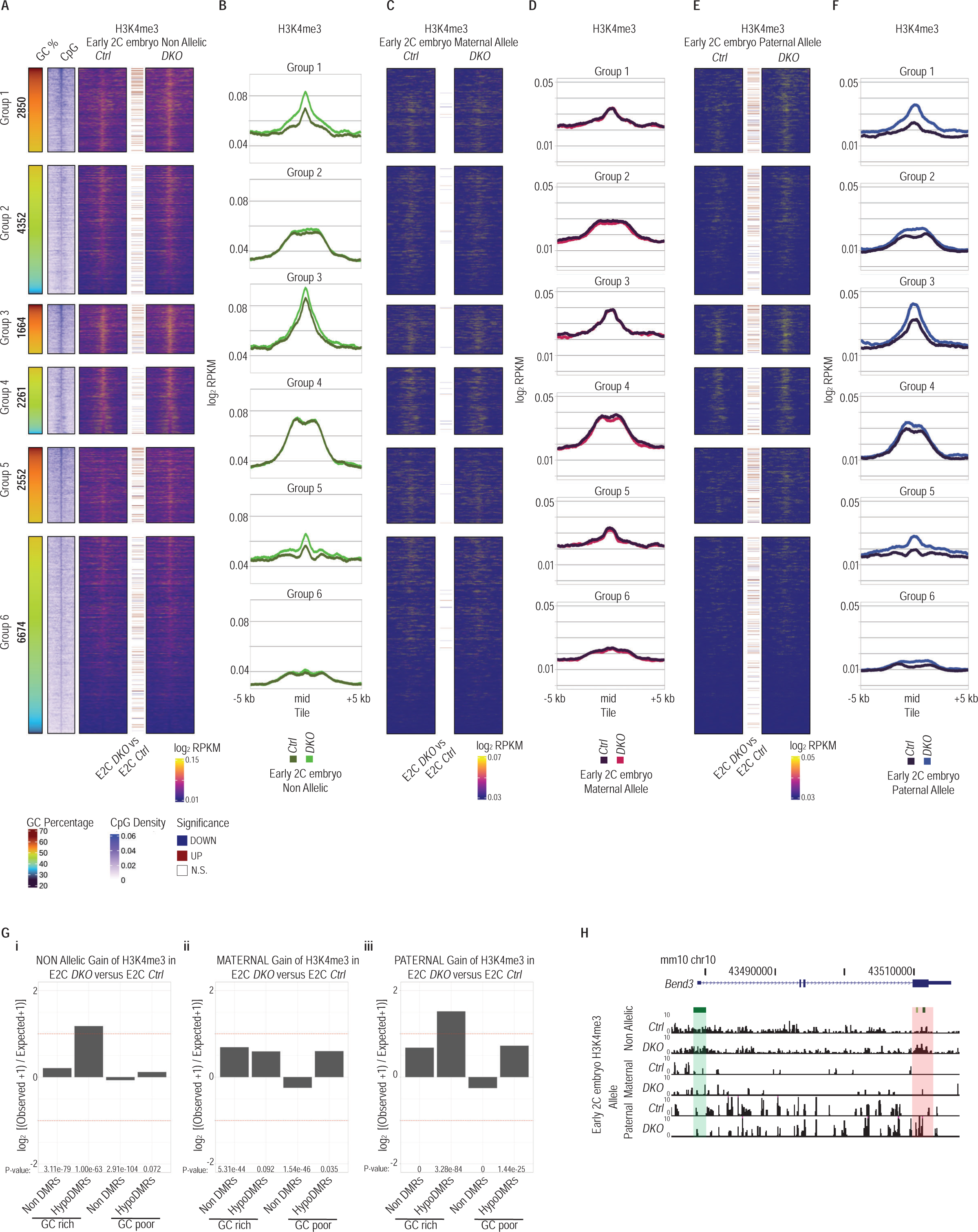
hypoDMRs acquire H3K4me3 in early mouse embryos generated by *DKO* sperm. A) Heatmaps showing H3K4me3 signal at the center of hypoDMRs (±5kb) assessed by ATATA-seq in early 2-cell (E2C) embryos generated by *Ctrl* (n=3) or *DKO* (n=3) sperm. Colored lines between heatmaps denote hypoDMRs with significantly differential H3K4me3 occupancy between indicated samples. The GC percentage and CpG density at hypoDMRs (±5 kb) are indicated. B) Line plots showing average H3K4me3 signal for each group of hypoDMRs (±5kb) assessed by ATATA-seq in early 2-cell (E2C) embryos generated by *Ctrl* (n=3) or *DKO* (n=3) sperm. C, E) Heatmaps showing H3K4me3 signal of maternal (C) or paternal (E) alleles at the center of hypoDMRs (±5kb) assessed by ATATA-seq in early 2-cell (E2C) embryos generated by *Ctrl* (n=3) or *DKO* (n=3) sperm. Colored lines between heatmaps denote hypoDMRs with significantly differential H3K4me3 occupancy between indicated samples. The GC percentage and CpG density at hypoDMRs (±5 kb) are indicated. D, F) Line plots showing average H3K4me3 signal at maternal (D) or paternal (F) alleles for each group of hypoDMRs (±5kb) assessed by ATATA-seq in early 2-cell (E2C) embryos generated by *Ctrl* (n=3) or *DKO* (n=3) sperm. G) Bar plots showing log2 enrichments of GC-rich or GC-poor nonDMR and hypoDMR tiles among tiles with significant H3K4me3 gain at (i) all alleles, (ii) maternal alleles and (iii) paternal alleles in early 2-cell (E2C) embryos generated by *Ctrl* or *DKO* sperm. P-values were calculated by Fisher’s exact test. H) UCSC genome browser snapshot of the Bend3 locus, showing tracks for H3K4me3 occupancy non-allelically, or at maternal or paternal alleles in *Ctrl* and *DKO* sperm. The promoter CGI and the hypoDMR are highlighted in green and red respectively.

Examination of total (non-allelic discrimination) ATATA-seq signal at hypoDMRs revealed an increased H3K4me3 signal primarily at all GC-rich groups in *DKO* sperm-derived embryos (Figure 6A, 6B, 6H). Upon assignment of the parental origin of ATATA-seq reads, we observed that H3K4me3 signals in the maternal genome were largely unchanged in regions examined between embryos generated by sperm of *Ctrl* or *DKO* males (Figure 6C, 6D, 6H). In striking contrast, the paternal genome demonstrated differential enrichments comparable to those observed in the non-allelic data. E.g., paternal alleles from embryos fathered by *DKO* sperm displayed increased H3K4me3 signals at hypoDMRs in GC-rich groups 1,3,5 and moderate increases at group 2,4,6 hypoDMRs, compared to those from *Ctrl* sperm-derived embryos, which had only low occupancy levels (Figure 6E, 6F, 6H). The magnitude of H3K4me3 changes in GC-rich hypoDMRs in *DKO* sired embryos associated to the extent of DNAme loss (Supplementary Figure 13D). These observations indicate that low DNAme in sperm renders paternal alleles in early embryos permissive for H3K4me3 deposition.

### Hypo-DNAme rather than H3K4me3 occupancy in sperm is associated with H3K4me3 deposition in paternal embryonic chromatin

We next asked whether increased H3K4me3 occupancy, or the absence of DNAme at hypoDMRs in *DKO* sperm promotes H3K4me3 deposition at paternal alleles in 2-cell stage embryos. To address this question, we focused on group 1 hypoDMRs displaying the most extensive changes in chromatin variables in *DKO* sperm and *DKO* sperm-derived embryos. We classified these hypoDMRs into three clusters based on increased, unaltered or reduced H3K4me3 occupancies in *DKO* versus *Ctrl* sperm and then quantified corresponding H3K4me3 occupancies in embryos (Figure 7A, 7B). Irrespective of the cluster identity, GC% or CpG densities, paternal alleles displayed similarly increased H3K4me3 levels in *DKO* sperm-derived embryos compared to *Ctrl* (Figure 7A, 7B). Hence, our measurements support the notion that H3K4me3 deposition in embryos does not require H3K4me3 occupancy in sperm, nor that H3K4me3 in sperm promotes H3K4me3 deposition in embryos. Instead, the data indicate that sperm borne DNA methylation restricts early H3K4me3 deposition at CpG-rich regions in paternal chromatin.

**Figure 7.**
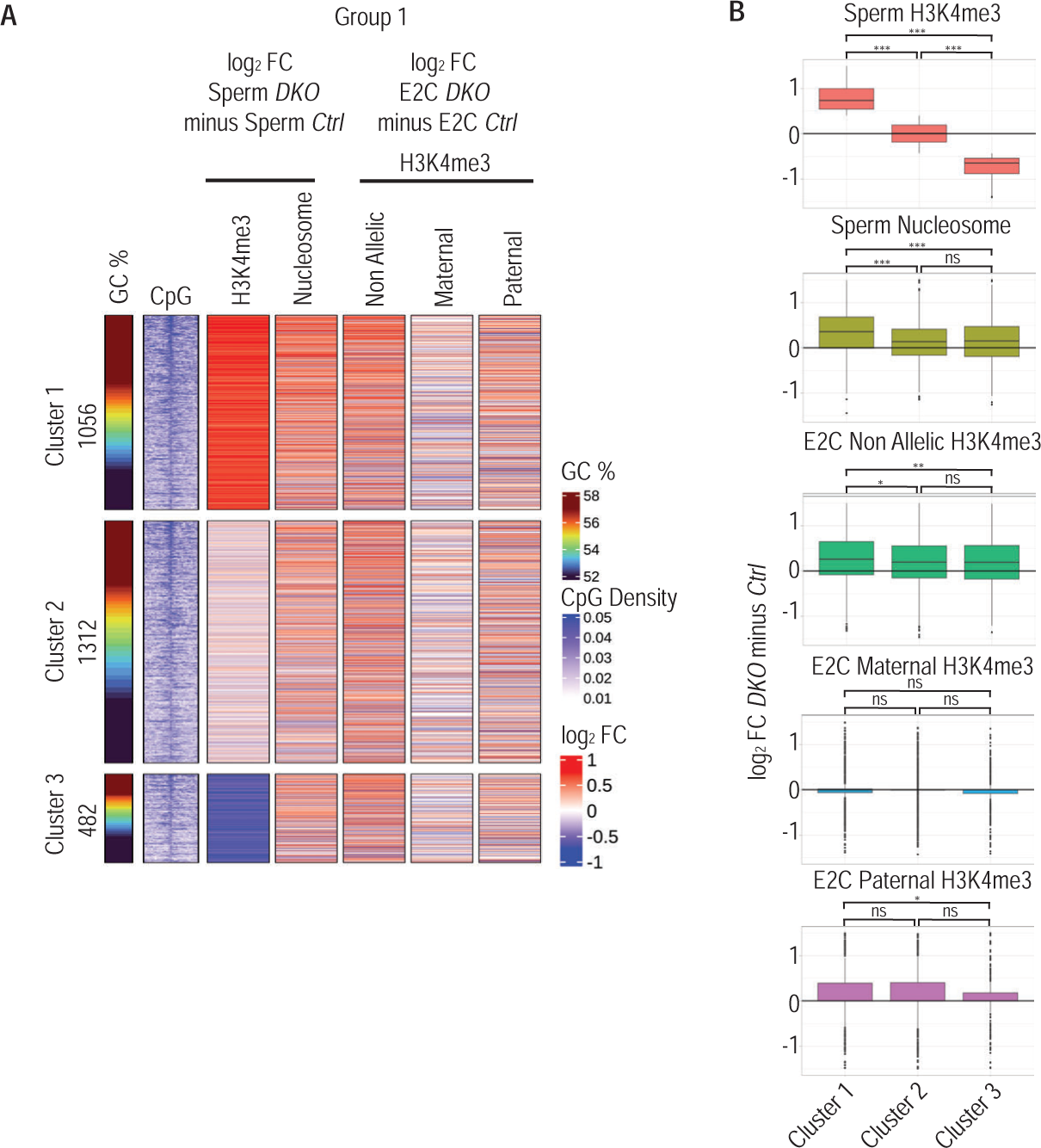
Comparison of intergenerational H3K4me3 changes at hypoDMRs. A) Heatmap showing the differential H3K4me3 and nucleosome occupancy between *DKO* and *Ctrl* sperm samples and differential H3K4me3 in E2C embryos generated by *DKO* and *Ctrl* sperm for Group 1 hypoDMRs. Group 1 hypoDMRs were clustered in 3 subclusters according to H3K4me3 differential enrichments between *DKO* and *Ctrl* sperm samples. B) Boxplots showing the H3K4me3 log2 fold changes between *DKO* and *Ctrl* sperm or E2C embryos generated by *DKO* and *Ctrl* sperm for the 3 subclusters of Group 1 hypoDMRs. Two-sample Wilcoxon tests were performed between indicated groups * P < 0.05, ** P ≤ 0.01, *** P ≤ 0.001.

### Absence of altered transcription around hypoDMRs in *DKO*-sperm-sired early embryos

As described above *DKO* males sired similar numbers of healthy offspring without evident phenotypic abnormalities as compared to *Ctrl* males (Supplementary Figure 1G). Nonetheless, to identify possible functional consequences of an altered paternal DNA methylome on transcription in early embryos, we performed RNA-sequencing analysis of single early 4-cell stage embryos after the occurrence of zygotic genome activation (n= 21 for *Ctrl* and n= 20 for *DKO* sperm derived embryos). Following developmental pseudo-time calculations, we selected age-matched embryos to perform differential gene expression analysis (n= 7 for *Ctrl* and n= 7 for *DKO* sperm derived embryos) (Supplementary Figure 14A). We identified only a handful of differentially expressed genes, which notably didn’t overlap with genes containing hypoDMRs (Supplementary Figure 14B, 14C, 14D).

To identify non-gene related transcriptional changes we quantified Smart-Seq2 reads around hypoDMRs and stratified them according to their expression levels in *Ctrl* embryos into 4 groups. We detected a trend towards increased transcription in lowly expressed regions in *DKO* sperm-derived embryos compared to *Ctrl*. However, this trend was evident in both paternal and maternal alleles suggesting that these transcriptomic alterations may be indirect consequences of the altered DNA methylome in *DKO* sperm (Supplementary Figure 14E). The absence of evidence for direct transcriptional mis-regulation may be due to that the majority of hypoDMRs reside in genomic regions unlikely to serve regulatory functions. For those that could confer regulation, the cofactors necessary to drive their expression may not be present at the 4-cell stage, when we performed our transcriptional profiling.

## DISCUSSION

### Molecular cues of intergenerational epigenetic inheritance

Our study demonstrates that the DNAme state in sperm impacts on the establishment of chromatin states at the paternal genome in early embryos. Sperm borne DNAme at GC-rich regions (Figure 1F) constrains *de novo* deposition of H3K4me3 at paternal alleles in early embryos (Figure 6E, 6F), while its absence at such regions is permissive. Interestingly, global DNAme present in sperm undergoes rapid removal upon fertilization in zygote embryos, except at ICRs and certain repetitive element families (67). Reanalysis of publicly available methylome data shows that most hypoDMR sequences identified in this study undergo such rapid DNAme removal, when paternally inherited (Supplementary Figure 15A, 15B, 15C) (25). In contrast, maternally speaking, the same hypoDMRs methylated in mature oocytes undergo only protracted, presumably replication-related, loss of DNAme during pre-implantation development. Intriguingly, GC-rich hypoDMRs gain H3K4me3 in wild type early embryos, concurrently to DNAme removal (Supplementary Figure 15D, 15E)(66). Given that the extensive erasure of DNAme from the paternal genome occurs prior to replication in one-cell embryos (68,69), we hypothesize that the *de novo* H3K4me3 deposition at hypoDMRs in paternal chromatin of *DKO* sperm-derived embryos occurs even prior to or concurrent with the wave of global DNA demethylation, possibly through precocious recruitment of H3K4 methyltransferases to CpG-rich unmethylated chromatin. Most other CpG-rich sequences, frequently found at gene promoters that remain largely unmethylated in sperm, may undergo a similar process, mechanistically and timing wise. Intriguingly, H3K4 methylation catalyzed on maternally provided H3.3 histones that become incorporated into the paternal genome is essential for the onset of minor ZGA in the paternal pronucleus and subsequent preimplantation development (26). In contrast to the distinct narrow peaks of H3K4me3 on promoters in sperm, paternal alleles in zygotes exhibit large domains of H3K4me3 (70,71), arguing for *de novo* activity of the KMT2A, KMT2B and/or SETD1B/CXXC1 enzymes recognizing unmethylated GC-rich regions via their CXXC domains, possibly followed by spreading *in cis* (72–75). Analogously to paternal transmission, it will be important to determine whether low levels of DNAme in oocyte genome primes H3K4me3 deposition not only in the oocyte (72) but also on maternal chromatin in the embryo.

A multitude of studies investigated the role of H3K4me3 in diet-induced phenotype models of epigenetic inheritance and suggested that paternal H3K4me3 is transmitted to the embryo and influences gene expression and development (32,76). While we detected differential H3K4me3 enrichments at a subset of GC-rich hypoDMRs in sperm, we did not observe a direct correlation of such differential enrichments to differential H3K4me3 enrichments observed at many more hypoDMRs in two-cell embryos (Figure 7A, 7B). Thus, our data argues that H3K4me3 is *de novo* deposited in early zygotes in response to the absence of sperm borne DNAme, rather than instructed by sperm-borne H3K4me3. Some hypoDMRs with clear gains in H3K4me3 in *DKO* sperm exhibited gains of H3K4me3 signal in *DKO* sperm-derived early embryos. Hence, we cannot exclude the possibility that high H3K4me3 occupancy in sperm may potentially enhance the deposition of H3K4me3 by e.g. the maternally provided SETD1B/CXXC1 enzyme complex reading unmethylated CpG dinucleotides and H3K4me3 via the CXXC and PHD domains of the CXXC1 protein, respectively (77–80). PHD-finger domain specific maternal loss-of-function studies of *Setd1b*/*Cxxc1*, or alternatively of *Kmt2a* or *Kmt2b* may contribute to deciphering the contribution of paternal H3K4me3-bearing nucleosomes for paternal inheritance of a H3K4me3 state. Irrespectively, our data support the model that the absence of DNAme serves as an effective intergenerational signal, enabling early H3K4me3 deposition in the next generation.

Over the past years, an increasing body of studies has investigated changes of sperm DNAme in mediating paternal epigenetic inheritance of acquired phenotypes (81,82). Molecular modes of inheritance remain, however, unclear. For example, changes in sperm DNAme upon paternal cigarette smoking were not recapitulated in offspring prefrontal cortex tissues (83). In another study, aged male mice exhibited local hypoDNAme in sperm at sites enriched for REST/NRSF binding motif. These males sired progeny with aberrant neuronal gene expression in forebrains during their embryonic development, which was linked to REST/NRSF motifs. At adulthood, the progeny displayed abnormal vocal communication patterns (84). Building upon our results providing a proof of concept directly linking changes in DNAme at specific CGIs in sperm to an altered H3K4me3 chromatin state in early embryos, the differential impact of REST/NRSF occupancy on DNAme and H3K4me3 in paternal transmission may be an interesting avenue to pursue further (85,86)

### Dynamics of DNAme and interplay with chromatin modifications

We show that the *de novo* DNMT3A and DNMT3B enzymes fulfill distinct non-redundant functions in *de novo* DNAme and even in DNAme maintenance during adult spermatogenesis. Our FACS sorting protocol enabled us to isolate undifferentiated spermatogonia (SgU), including self-renewing spermatogonial stem cells (SSC), and differentiated spermatogonia (SgD). Surprisingly in SgU, hypoDMRs with mid-to-high levels of DNAme were more sensitive to depletion of DNMT3A than DNMT3B (Supplementary Figure 5B), suggesting that DNMT3A is required to maintain DNAme. DNMT3A may possibly counteract TET activity in SgU (87) and/or augment inefficient maintenance by DNMT1. Previously it was shown that depletion of DNMT3A from embryonic male gonocytes resulted in differentiation defects of spermatogonia after birth. In these studies, the depletion resulted in a greatly hypomethylated genome of spermatogonia (10,13). In the current study, most genomic methylation was established before the deletion of *Dnmt3a*. Hence, we interpret that the global pattern of DNAme established in gonocytes is maintained by DNMT1 (43).

In contrast, DNAme levels at hypoDMRs in DNMT3B depleted SgUs resembled those in *Ctrl* SgUs suggesting minimal contribution of this enzyme in undifferentiated spermatogonia. However, the DNAme profile at hypoDMRs in DNMT3B depleted SgDs resembled those measured in *Ctrl* and DNMT3B-depleted SgU cells, suggesting that the developmental gain of DNAme at hypoDMRs during spermatogonial differentiation is defective (Figures 2D, Supplementary Figure 5B). In addition, when SgUs depleted for DNMT3A differentiate to SgD we observed *de novo* gains of DNAme which are mediated by DNMT3B, albeit towards altered levels due to the abnormal initial DNAme status existing in DNMT3A-depleted SgU. Previously it was shown that DNMT3B preferentially located at gene bodies of transcribed genes (52), possibly through interactions between its PWWP domain and the H3K36me3 modification, deposited by the RNA polymerase II interacting histone methyltransferase SETD2. Indeed, genes that contain hypoDMRs were transcriptionally active both at SgU and SgD stages (Supplementary Figure 6A), displaying already mid-high levels of H3K36me3 in SgUs (Supplementary Figure 8A), posing the question why high DNAme levels are established only in SgD. The lowly DNA methylated hypoDMRs are embedded in broader H3K36me3 domains in both cell types, however they are highly enriched in H3K4me3 at the SgU stage (Figure 3A). We postulate a constant tug of war for recruitment of DNMT3 enzymes and subsequent establishment of DNAme via the H3K36me3-rich flanks of hypoDMRs and a repulsion of DNMT3 enzymes from the H3K4me3-rich center of hypoDMRs at SgU stage, which is resolved by the SgD stage with the establishment of high DNAme. In female mice, demethylation of H3K4 by LSD2 (KDM1B) is critical for establishing the DNAme imprints during oogenesis, pointing out that H3K4 demethylases are genetically upstream of DNAme (88). H3K4 demethylases such as LSD2 and KDM5B are recruited to gene bodies by NPAC and MRG15 respectively - two putative H3K36me3 readers (89,90). In oocytes, an interplay between H3K4me3, H3K36me3 and DNAme was observed where loss of H3K36me3 led to failure in establishing the correct DNA methylome, and invasion of H3K4me3 and H3K27me3 into former H3K36me3 territories (56). Additionally, LSD1 (KDM1A) binds *in vitro* to DNMT1 and DNMT3B (91) and functionally impact on DNAme levels in mouse ESCs (92). In analogy to oocytes, we propose that H3K4 demethylases may reprogram H3K4me3 at hypoDMRs during spermatogonial differentiation. Intriguingly, H3K4me3 occupancy persisted at hypoDMRs in SgD in the absence of the DNMT3A/DNMT3B DNAme machinery, suggesting that these enzymes may regulate recruitment of H3K4 demethylases and/or that the absence of DNAme enables H3K4 methyltransferases recruitment and H3K4me3 deposition (Figure 3A). Examining the effect of abnormal H3K4me3 reprogramming on DNAme in the male germ line may, however, be challenging as loss of (93,94)developmental arrest (93,94).

### Sperm chromatin and intergenerational epigenetic inheritance

We previously inferred from MNase protection and ChIP assays for histones and their modifications the genomic localization of nucleosomes/histones in mature mouse and human sperm (3,20). We observed an inverse correlation between high DNAme and nucleosome occupancy at CpG-rich sequences (3). Subsequent MNase studies reported the presence of nucleosomes in intergenic regions and repetitive elements (21,95). Experimental conditions aggravated by a complex chromatin structure in sperm nuclei are likely at the core of these inconsistencies. Among others, they may originate from variability in pretreatments, degrees of protamine reductions and extractions, enzymatic digestions required to solubilize sperm chromatin, and from antibody pulldown efficiencies (17,24,96). Moreover, given the low levels of histones in sperm, purity of spermatozoa samples is critical (20,23,24). Consequently, the distributions and the relative abundances of nucleosomes in mammalian sperm genomes remain unclear.

Here, by FACS sorting highly pure sperm cell samples, as indicated by our DNAme measurements at maternal ICRs (data not shown), and comparing nucleosome ChIP assays on different genetic backgrounds of wild type and *Dnmt3a/Dnmt3b* deficient sperm allowed us to examine the impact of DNAme status on nucleosome occupancy at various genomic sequence composition contexts in a comparative and germ cell specific manner. As before (3), we measured that normally methylated GC-rich regions exhibited low nucleosome signal in *Ctrl* sperm. In *Dnmt3a/Dnmt3b* mutant spermatozoa, the same regions that had become hypomethylated had increased nucleosome signals. We measured only marginal increases in nucleosome signals at GC-poor regions that had become hypomethylated in the mutant or in GC-rich regions that showed only small reduction of DNAme. Together, these data support the model that indeed the combination of GC-richness and low DNAme promote nucleosome retention in sperm (Figure 4A).

The ChIP-seq methods employed in this study examine relative nucleosome enrichments in large bulk populations of sperm limiting our understanding of the heterogeneity among spermatozoa that would contain nucleosomes at a given genomic region. Future studies examining sperm chromatin variability at a single cell resolution will be required to provide more quantitative insights to the extent by which the presence or absence of DNAme would modulate nucleosome eviction during spermiogenesis. Importantly, beyond modulating sperm chromatin, this study provides a molecular paradigm of how the DNA sequence and its methylation status in mature male gametes define the establishment of the chromatin landscape at the paternal genome in the early embryo in terms of H3K4me3 deposition.

## MATERIAL & METHODS

### Mice and early embryos collection

Animal housing, handling and procedures of mice conformed to the Swiss Animal Protection Ordinance and approved by the institutional ethical committee. Mice bearing *Dnmt3a^flox^* (*Dnmt3a^tm3.1Enl^*) and *Dnmt3b^flox^* (*Dnmt3b^tm5.1Enl^*) alleles (97) were provided by Dr. En Li. Mice bearing the iCre expressing transgene under the control of *Stra8* promoter (*Tg(Stra8-icre)1Reb*) (45) were obtained from The Jackson Laboratory (RRID:IMSR_JAX:008208). The JF1/MsJ (Japanese fancy mouse 1) inbred strain (98) was purchased from The Jackson Laboratory (RRID:IMSR_JAX:003720). C57BL/6JRj mice were purchased from Janvier Labs. To generate the *Dnmt3a*; *Dnmt3b*; *Stra8-iCre* conditional mice, we first intercrossed mice from the single *Dnmt3a*^fl/fl^ and *Dnmt3b*^fl/fl^ colonies to generated double *Dnmt3a*^fl/fl^; *Dnmt3b*^fl/fl^ homozygote females which were then crossed with males from *Stra8-iCre* colony to generate *Dnmt3a*^fl/+^; *Dnmt3b*^fl/+^; *Stra8-iCre* male mice. *Dnmt3a*^fl/+^; *Dnmt3b*^fl/+^; *Stra8-iCre* males were backcrossed to C57BL/6JRj females to create *Dnmt3a*^-/+^; *Dnmt3b*^-/+^; *Stra8-iCre* males. *Dnmt3a*^-/+^; *Dnmt3b*^-/+^; *Stra8-iCre* males were then mated with *Dnmt3a*^fl/fl^; *Dnmt3b*^fl/fl^ homozygote females to generate double or single conditional mutants *Dnmt3a*^-/fl^; *Dnmt3b*^-/fl^; *Stra8-iCre* (*DKO*), *Dnmt3a*^-/fl^; *Dnmt3b*^+/fl^; *Stra8-iCre*, (*3aKO*), *Dnmt3a*^+/fl^; *Dnmt3b*^-/fl^; *Stra8-iCre* (*3bKO*), and double heterozygote *Dnmt3a*^+/fl^; *Dnmt3b*^+/fl^; *Stra8-iCre* (HET) or wildtype *Dnmt3a*^+/fl^; *Dnmt3b*^+/fl^ (*Ctrl*) male experimental mice. Single conditional mutant mice for one Dnmt3 paralogue are heterozygous for the other.

To obtain early embryos, 7-10-week-old JF1 female mice were injected intraperitoneally with HyperOva (Cosmo Bio, KYD-010-EX-X5) 63 hours pre collection and human chorionic gonadotropin (hCG, Chorulon MSD) 15 hours pre collection. Oviducts were dissected and oocytes from 2 animals were collected in 30 μl drops of HTF medium (EmbryoMax® Human Tubal Fluid Millipore, MR-070-D). Sperm was collected from 2 cauda epididymides in 50 μl drops of HTF medium. 2-5 μl of sperm sample was transferred to the oocyte drops and incubated for 3 hours at 37 °C. Zygotes with visible pronuclei washed 3 times in HTF medium and transferred to KSOM medium (Millipore, MR-106-D) and cultured in low oxygen chamber filled with the mixture gas (5% O2/ 5% CO2). Early 2 cell stage embryos were collected 22 hours post fertilization. Early 4 cell stage embryos were collected 50-55 hours post fertilization.

### Fluorescence Activated Cell Sorting (FACS) of spermatogonia and sperm

Testicular cell suspension was prepared by incubation of seminiferous tubules in 200 U/ml Collagenase type I (Worthington Biochemical LS004196), 5 μg/ml DNAse I (Roche 10104159001), and 0.05%Trypsin (Gibco 25200056) in GBSS (Sigma G9779) as described in (99). Cells were stained with 1/200 anti-CD117/c-kit PE (eBioscience 12-1171-83), anti-CD324/E-Cadherin eFluor 660 (eBioscience 50-3249-82) and anti-CD49f/Integrin alpha 6 PE-Cyanine7 (eBioscience 25-0459-82) for 1 hour at 32°C with constant shaking and protected from light. After washing cells were incubated with 20 μg/ml with Hoechst 33342 (Thermo Fischer Scientific H3570) for 1 hour at 32°C with constant shaking and protected from light and then with 30nM DRAQ7 (Biostatus DR71000) for 5 minutes on bench. The stained cells were strained through a 40 μm nylon filter into 5 ml polypropylene tubes and were sorted on a BD FACSAria III cell sorter fitted with a 70 μm nozzle (Becton Dickinson) using a 375nm laser to excite Hoechst 33342, a 561nm laser to excite PE and PE-Cy7 and a 633nm laser to excite DRAQ7 and eFluor600. Cells were first gated for FSC and SSC to exclude debris. Then the live cells were selected based on the absence of DRAQ7 signal detected using 755LP, 780/60BP. Then we gated for cells positive for Hoechst 33342 emission which was detected using a 670LP (Hoechst-Red) and 450/20 BP (Hoechst-Blue). We gated for the Hoechst-Red^low^ and Hoechst-Blue^mid^ population which is enriched for 2N spermatogonia. Fluorescence for PE was detected using a 582/15BP filter, for PE-Cy7 was detected using 735LP,780/60BP filter and for eFluor660 using a 660/20BP filter. Undifferentiated spermatogonia were sorted as CD324^high^, CD49f^high^, CD117^low^ and differentiated spermatogonia were sorted as CD324^low^, CD49f^low^, CD117^high^.

For sperm cell sorting, mouse cauda epididymides were dissected into a Petri dish and fat patches were removed with forceps and scissors. Each epididymis was punctured with a needle and carefully squeezed into 100 ul of PBS with the help of two forceps. Sperm cell suspension was transferred into an Eppendorf tube and was allowed to liquefy at room temperature for 5 minutes. To break sperm tails, cells were briefly sonicated with a Brandson Tip digital sonicator with 10% amplitude and 3 cycles of 0.5 sec ON / 2 sec OFF. Cells were stained with 2 ul/ml Hoechst 33342 (H3570, ThermoFisher) for 1h at 25°C with constant shaking and protected from light. Cells were filtered through a 40 μm Nylon filter into 5 ml polypropylene tubes prior to sorting into PBS with a BD FACS Aria III instrument. Cells were first gated for FSC and SSC to exclude debris. Then sperm cells were sorted according to their 1C DNA content.

### Reduced Representation Bisulfite Sequencing (RRBS)

Pellets of 1 million sperm cells were incubated in 200μl of DNA extraction buffer (20mM Tris-HCl pH 7.5, 300mM NaCl, 2% SDS, 10mM EDTA, 200 μg/ml Proteinase K) for 16 hours at 55°C. An equal volume of phenol:chloroform:isoamyl alcohol (25:24:1) solution was added to the samples. After a vigorous vortexing for 30 seconds and centrifugation at 16.000g for 10 minutes, the upper aqueous phase was transferred to new tubes. 0.05 μg/μl glycogen, 1/10 volume of 3M sodium acetate and 2.5 volumes of 100% ethanol were added, and the samples were incubated for 2 hours at −80°C. DNA precipitated by centrifugation at 16000g for 30m minutes at 4°C. DNA pellet was washed once with 70% ethanol and resuspended in 20 μl. DNA was further purified using 2 volumes of Ampure XP beads (Beckman Coulter). DNA was extracted from 10000 undifferentiated or differentiated spermatogonia cell samples by QIAamp DNA Micro Kit (Qiagen # 56304) according to the manufacturer’s instructions. We prepared RRBS libraries using 10-20 ng of DNA and following the previously published method (100). Briefly, DNA was digested with 0.9μl MspI Fast Digest (Thermo Scientific FD0544) in 1X Tango buffer in a volume of 18 μl for 6 hours at 37°C. DNA ends were repaired by adding 1 μl Klenow Fragment, exo– (Thermo Scientific EP0421) and 1µl of nucleotide end-repair mix (1mM dATP; 0.1mM dGTP; 0.1mM dCTP prepared in 2X Tango buffer) and incubated for 45min at 37ᵒC, followed by enzyme heat inactivation at 75ᵒC for 15min. Then 6.25nM of 5mC sequencing adapter (NEB E7535S) was ligated with 1 μl T4 DNA Ligase, HC (Thermo Scientific EL0013) and 0.5 μl ATP solution (Thermo Scientific R0441) for 16 hours at room temperature. The looped adaptor ligated DNA fragments were subjected to USER (NEB M5508) digestion at 37 ᵒC for 20 minutes and used directly for bisulfite treatment using the Imprint DNA Modification Kit (Sigma MOD50) according to the manufacturer’s 2-step modification procedure. The modified DNA was amplified with 13 cycles of PCR using Kapa U+ polymerase (Roche 07959052001) and indexed primers (NEB) and purified with 1.8X Ampure XP beads. Libraries were sequenced on a HiSeq 2500 sequencing platform generating 50 bp single end reads, per the manufacturer’s recommendations.

### Whole genome methylation sequencing

Genomic DNA (100ng per reaction) was sheared to an average fragment size of 400 bp using the Covaris S220 instrument. The sheared DNA was used as input for the NEBNext Enzymatic Methyl-seq (NEB E7120) following the manufacturer’s instructions except for doubling the reaction incubation times. The libraries were amplified for 5 cycles and sequenced in 2×75-bp paired-end mode with NextSeq 500 sequencing technology (Illumina), per the manufacturer’s recommendations.

### Chromatin Immunoprecipitation (ChIP)

To profile histone modifications in spermatogonial cells we followed the previously published ultra-low input native ChIP-seq (ULI-NChIP-seq) protocol (57) with minor modifications. Briefly 5000-7500 spermatogonia were FACS-sorted directly in 20 μl nuclear isolation buffer (Sigma-Aldrich NUC101) in Eppendorf® LoBind microcentrifuge tubes. The nuclei were centrifuged at 500g for 5 minutes at 4°C in a swing bucket centrifuge and the volume of the sample was reduced to 10 μl. The sample was snap frozen in liquid nitrogen and stored at –80°C freezer until further processing. The sample was thawed on ice and 1 μl of TS buffer (5% Triton X-100, 5% Sodium deoxycholate) was added for 10 minutes to ensure permeabilization of nuclei. 40 μl of MNAse digestion buffer (3.33 gelUnits/μl Micrococcal Nuclease (NEB M0247S), 1.25X Micrococcal Nuclease buffer, 6.25% PEG 6000, 0.85mM DTT) was added to the sample for 10 minutes at 25 °C to fragment the chromatin. Reaction was stopped by addition of 5 μl 100 uM EDTA and the sample was placed on ice. The sample was diluted by addition of 145 μl ChIP buffer (20 mM Tris-HCl pH 8.0, 2 mM EDTA, 150 mM NaCl, 0.1% Triton X-100, 1X EDTA-free protease inhibitor cocktail). To ensure release of chromatin fragments, the sample was sonicated using the Diagenode Bioruptor Plus® for 3 cycles of 5 seconds ON and 30 seconds OFF at low output. Then chromatin was pre-cleared with 10 μl of 1:1 protein A:protein G Dynabeads (Thermo Fisher Scientific, 10001D and 10003D). The precleared chromatin sample was incubated with 0.5 μg of anti-H3K4me3 (Millipore 17-614) or 0.75 μg anti-H3K36me3 (Cell Signaling #4909) conjugated with 10 μl of 1:1 protein A: protein G Dynabeads overnight at 4 °C on a rotator. Chromatin-Antibody-Dynabeads complexes were washed once with 500 μl of ChIP buffer, 3 times with 1ml of Low Salt wash buffer (20 mM Tris-HCl (pH 8.0), 0.05% SDS, 1% Triton X-100, 2 mM EDTA and 150 mM NaCl) and 3 times with 1 ml High Salt buffer (20 mM Tris-HCl (pH 8.0), 0.05% SDS, 1% Triton X-100, 2 mM EDTA and 250 mM NaCl). Chromatin was eluted in 40 μl of Elution buffer (100 mM NaHCO3 and 1% SDS) for 90 minutes at 65 °C. DNA from eluted material was purified by 2X volumes of Ampure XP DNA purification beads (Beckman Coulter, A63881) and resuspended in 25 μl nuclease free H2O.

To profile histone modifications in spermatozoa, we followed the previously published native ChIP-seq protocol (24) with modifications as follows. Spermatozoa (500.000 per reaction) were FACS sorted in Eppendorf® LoBind microcentrifuge tubes. The cells were pelleted at 6000g for 5 minutes at 4°C in a benchtop centrifuge. The pellet was snap frozen in liquid nitrogen and stored at –80°C freezer until further processing. The pellet was thawed on ice and resuspended in 500 μl PBS. 50mM DTT was added for 2 hours at 21°C to reduce disulfide bonds of protamine. 100mM N-Ethylmaleinimid was added for 30 minutes at 21°C to modify the cysteine residues. The cells were pelleted at 6000g for 5 minutes at 4°C in a benchtop centrifuge and washed once with 600 μl PBS. Cells were resuspended in 50 μl Lysis buffer (20 mM Tris-HCl pH 7.5, 60 mM KCl, 5mM MgCl2, 0.1 mM EGTA, 300 mM Sucrose, 0.5 mM DTT, 0.25 % Igepal, 0.5% sodium deoxycholate, 1X EDTA-free protease inhibitor cocktail) and incubated for 30 minutes at 21°C. 50 μl of MNAse digestion buffer (5 gelUnits/μl Micrococcal Nuclease (NEB M0247S), 2X Micrococcal Nuclease buffer) was added to the sample for 15 minutes at 37°C to fragment the chromatin. Reaction was stopped by addition of 5 μl 500 uM EDTA and the sample was centrifuged at 16000g for 10 minutes at 4°C in a benchtop centrifuge. The supernatant chromatin was pre-cleared with 10 μl of 1:1 protein A: protein G Dynabeads (Thermo Fisher Scientific, 10001D and 10003D). The precleared chromatin sample was incubated with 0.5 μg of anti-H3K4me3 (Millipore 17-614) or 1 μg anti-nucleosome (58) provided by J. van der Vlag) or 1 μg anti-H3.3 (Cosmo Bio CE-040B) or 1 μg anti-H3.1/2/t (101) provided by J. van der Vlag) conjugated with 10 μl of 1:1 protein A:protein G Dynabeads overnight at 4 °C on a rotator. Chromatin-Antibody-Dynabeads complexes were washed once with 500 μl of ChIP buffer, 3 times with 1ml of Low Salt wash buffer (20 mM Tris-HCl (pH 8.0), 0.05% SDS, 1% Triton X-100, 2 mM EDTA and 150 mM NaCl) and 3 times with 1 ml High Salt buffer (20 mM Tris-HCl (pH 8.0), 0.05% SDS, 1% Triton X-100, 2 mM EDTA and 250 mM NaCl). Chromatin was eluted in 40 μl of Elution buffer (100 mM NaHCO3 and 1% SDS) for 90 minutes at 65 °C. DNA from eluted material was purified by phenol-chloroform, ethanol-precipitation and resuspended in 25 μl nuclease free H2O.

For library construction the ChIP purified DNA was used as input for the NEBNext® Ultra™ II DNA Library Prep Kit for Illumina® (NEB E7645L) with modifications of the manufacturer instruction as we assembled half volume reactions, and we doubled reaction incubation times. For adaptor ligation we used 1:15 adaptor dilution. We amplified the ligated fragments for 12 cycles using single indexed primers. Libraries were pooled and sequenced in 50-bp single-end mode with HiSeq 2500 sequencing technology (Illumina), per the manufacturer’s recommendations.

### Antibody Targeted Tagmentation (ATATA)

ATATA-seq was based on the Cut&Tag strategy (64). We fused N terminally the Tn5 transposase to two tandem IgG binding Z domains to specifically target the ZZ-Tn5 to antibody bound regions. Early 2 cell embryos (50 embryos per reaction) were incubated in 30 ul LIB buffer (20mM HEPES pH 7.5, 300mM NaCl, 0.5mM Spermidine, 0.05% Digitonin, 0.05% Triton X-100, 1X EDTA-free protease inhibitor cocktail) on ice for 10 mins for nuclei permeabilization. Active transposomes were generated by incubating ZZ-Tn5 (8μM) with Mosaic End double-stranded (MEDS) oligonucleotides (12μM) for 30 minutes at 37°C. Active transposomes (0.4 μM) were incubated with anti-H3K4me3 antibody or normal rabbit IgG (1μM) in LIB buffer for 30 minutes at 4°C with continuous mixing. Antibody-ZZ-transposomes complexes (final concentration 0.1 μM) were added to the embryo samples and incubated for 1 hour at 4°C with continuous mixing to allow targeting to the chromatin. Antibody-ZZ-transposomes were activated by addition of TAB buffer (20mM HEPES, 5%PEG6000, 7.5mM MgCl2, 300mM NaCl) and incubated for 1 hour at 37°C with continuous mixing. The tagmentation reaction was stopped with addition of Stop buffer (0.2% SDS, 10mM EDTA, 100ug/ml ProteinaseK) and incubated at 65°C for 15 minutes. DNA was purified by phenol-chloroform, ethanol precipitation and libraries were amplified for 16 cycles. The libraries were purified by 0.5X-1.5X volumes left-right side of Ampure XP DNA purification beads and sequenced in 50-bp single-end mode with HiSeq 2500 sequencing technology (Illumina), per the manufacturer’s recommendations.

### RNA sequencing

For 4-cell stage embryos RNA-seq libraries were prepared following the previously published Smart-Seq2 method (102). Each 4-cell stage embryo was collected at 50-55 hours after IVF, was washed in PBS + 0,02% PVA, transferred into a well of a 96-well plate containing SS2 Lysis buffer (0,09% Triton-X 100, 0,5U/μl SUPERAseIN (Life Technologies AM2696), 2.5mM oligodT primer (Microsynth), 2.5mM dNTP mix(Promega), ERCC RNA Spike-In Mix (1:3.2×10^7^ Thermo Fischer scientific 4456740)), snap frozen on dry ice and then stored in −80°C until further use. The sample was incubated at 72°C for 3 minutes and immediately put back on ice. Then SS2 RT mix (10U/µl SuperScript II reverse transcriptase (Thermo Fischer 18064014), 0.25U/μl SUPERAseIN, 1X SuperScript II first-strand buffer, 5mM DT, 1M Betaine (Sigma B0300-1VL), 6mM MgCl2, 1mM TSO (Exiqon)) was added to the sample and incubated for 90 minutes at 42°C, then 10 cycles of 2 minutes at 50°C and 2 minutes at 42°C and finally for 15 minutes at 70°C. Sample was preamplified for 16 cycles using 1X KAPA HiFi HotStart ReadyMix (KAPA Biosystems KK2602) and 0.1µM ISPCR primers (Microsynth). DNA was purified using 1X volumes of Ampure XP DNA purification beads. 1ng of pre-amplified DNA was added to Tn5 tagmentation mix (1x TAPS-DMF buffer, homemade Tn5 (1:1,200)) in total volume 20μl and incubated at 55°C for 7minutes. Then the reaction was stopped by adding 5μl of 0.2% SDS and kept at 25°C for 7 min. Adapter-ligated fragment amplification was done using Nextera XT index kit (Illumina) in a total volume 50μl (1x Phusion HF Buffer, 2 µl of Phusion High Fidelity DNA Polymerase (Thermo Fischer, F530L), dNTP mix (0.3mM each) (Promega)) with 10 cycles of PCR. The library was purified by 1X volumes of Ampure XP DNA purification beads and resuspended in 25 μl nuclease free H2O. Sequencing was performed on an Illumina HiSeq 2500 machine with single-end 50-bp read length (Illumina), per the manufacturer’s recommendations.

RNA from Spermatogonia (10.000 per sample) was extracted with Single Cell RNA Purification Kit (Norgen, 51800) and libraries were prepared using the Ovation® SoLo RNA-Seq Library Preparation Kit (Nugen, 0501-32) following the manufacturer’s instructions. Libraries were sequenced in 2×38-bp paired-end mode with NextSeq 500 sequencing technology (Illumina), per the manufacturer’s recommendations.

### Whole mount seminiferous tubules immunohistochemistry

Seminiferous tubules were gently detangled and washed 2-3 times with PBS, to remove the interstitial cells. Then, the tubules were fixed with 4% PFA in PBS for 1 hour at 4°C. After fixation, the tubules were washed 3 times in PBS and then 3 times in PBS containing 0.04% Tween-20 (PBST) in a cell strainer, each wash for 10 mins. Then, the tubules were dehydrated through a graded series of 25%, 50%, 75% methanol containing PBST and 100% methanol at 4°C for 7 mins, respectively. The samples were stored at −80°C until later usage. At the time of observation, the samples were rehydrated through a graded series of 75%, 50%, 25% methanol containing PBST for 7 mins at 4°C and then washed 3 times with PBST for 10 mins each. Rehydrated tubules were blocked in PBST containing 4% Normal Donkey Serum (Abcam ab7475) for 1 hour. After blocking, the tubules were incubated with primary antibodies against cKit (1:1000 R&D systems AF1356), DNMT3a (1:1000 Imgenex IMG-268A) and DNMT3b (1:1000 Imgenex IMG-184A) for 3 hours at room temperature. Afterwards, the tubules were washed 3 times for 10 mins in blocking buffer followed by incubation with species specific secondary Alexafluor-conjugated antibodies (1:1000, ThermoFischer Scientific) for 2 hours. Finally, tubules were incubated in 0.001 mg/ml DAPI solution (Sigma D9542) for 10 mins and then washed 3 times with PBST for 10 mins. For imaging, tubules were oriented in PBST between a microscope slide and a coverslip separated by a 0.12 mm thick SecureSeal™ Imaging Spacers (Grace Bio Labs). Images were obtained using spinning disk confocal scanning unit Yokogawa CSU W1 Dual T2 with 100x/1.4 oil immersion objective. Representative regions were selected using Fiji software (103).

### Histopathology

Fresh whole testis samples were fixed in 5ml of Bouin’s solution (Sigma HT10132) for at least 48-72 hours and then stored in 70% ethanol until further processing. The fixed samples were embedded in paraffin using an automated tissue processing center (TPC 15 Duo, Medite) with standard settings. Sectioning was done at 3 um thickness using the automatic microtome (HM355S, Thermo Fisher Scientific). Sections were mounted onto Superfrost Plus Adhesion Microscope Slides (J1800AMNZ, Thermo Fisher Scientific) and dried at 37°C overnight. For staining, sections were deparaffinized by incubating in xylene solution (Sigma 534056) 2 times for 5 mins and rehydrated in a series of decreasing concentrations of ethanol (2x 100%, 95%, 70%, 3 min each) to deionized water. Rehydrated tissues were immersed in Periodic Acid Solution (Sigma, 395132) for 5 min at RT, rinsed several times in deionized water, immersed in Schiff’s Reagent (Sigma 3952016) for 15 min at RT and then rinsed in tap water for 5 min. Samples were counterstained with Mayer’s Hematoxylin Solution (MHS32, Sigma) for 2 mins and rinsed in tap water for 5 mins. Finally, samples were dehydrated in a series of increasing concentrations of ethanol (70%, 95%, 2×100%, 3 min each), cleared in xylenes solution (2 x 3min) and mounted with PermountTM mounting media (ThernmoFischer, SP15-100). Images were acquired using motorized automated slide Scanner Zeiss Axioscan Z1 with 40x air objective and analyzed with ZEN blue software (version 2.3, Zeiss).

### Computational Analysis of Sequencing data

Computational analyses were performed using custom made R scripts (R version 4.2.1). Mouse genome BSgenome.Mmusculus.UCSC.mm10 was tiled in 500bp non overlapping tiles. Tiles that overlap with blacklisted regions from (104) were removed from the subsequent analysis. Tiles containing less than 4 CpGs were excluded from the analysis as non-informative. For annotation of genomic features, coordinates of genes (including exons and introns) were extracted from TxDb.Mmusculus.UCSC.mm10.knownGene (version 3.10.0) and the promoter regions were defined as genes TSS +/- 1kb (Supplementary Figure 2B). CpG islands and repetitive element coordinates were downloaded from the UCSC “CpG Islands” and “RepeatMasker” tables respectively. Gametic DMR coordinates were obtained from (105).

EMseq (this study) and PBAT (from (47)) reads were first processed using TrimGalore (version 0.6.2) to trim adaptor and low-quality reads with settings (--stringency 3). EMseq and PBAT trimmed reads were then aligned to the mouse genome build mm10 using the qAlign function from the QuasR package and Bismark package, respectively. Methylated and unmethylated read counts for each CpG were extracted from the BAM files using the function qMeth from the QuasR package. Tiles covered with less than 25 reads on their CpGs were discarded from downstream analysis. The summed read count of methylated and unmethylated CpGs was calculated for each retained tile. The resulting read count data were processed by edgeR to identify differentially methylated tiles between experimental groups as described previously (106). Tiles with methylation percentage change >25% or <-25% and false discovery rate-adjusted FDR < 0.05 were considered to be significantly changed. The percentage of methylation was calculated as the fraction of methylated CpG read counts to the total CpG read counts for tile.

RRBS reads were first processed using TrimGalore (version 0.6.2) to trim adaptor and low-quality reads with settings (--rrbs, --stringency 3). Trimmed reads were then aligned to the mouse genome build mm10 using the qAlign function from the QuasR package. Methylated and unmethylated read counts for each CpG were extracted from the BAM files using the function qMeth from the QuasR package. CpGs covered with less than 5 reads were discarded from downstream analysis. In addition, tiles covered with less than 20 reads on their CpGs were discarded from downstream analysis. The summed read count of methylated and unmethylated CpGs was calculated for each retained tile. The resulting read count data were processed by edgeR to identify differentially methylated tiles among experimental groups. Tiles with methylation percentage change >20% or <-20% and false discovery rate-adjusted FDR < 0.05 were considered as significantly changed. The percentage of methylation was calculated as the fraction of methylated CpG read counts to the total CpG read counts for tile.

ChIP-seq and ATATA reads were first processed using TrimGalore (version 0.6.2) to trim adaptor and low-quality reads with settings (--stringency 3). Trimmed reads were then aligned to the mouse genome build mm10 using STAR (version 2.5.0a) with settings (-- alignIntronMin 1 --alignIntronMax 1 --alignEndsType EndToEnd --alignMatesGapMax 1000 -- outFilterMatchNminOverLread 0.85). ATATA samples for hybrid JF1/MsJ x C57BL/6JRj 2-cell embryos were separately aligned to C57BL/6J and JF1/MsJ genomes obtained by incorporating JF1 single-nucleotide polymorphisms (SNPs) into reference mm10 genome using previously published SNP table from (107). Reads were categorized as maternal (JF1/MsJ), paternal (C57BL/6JRj) or undefined based on minimal number of mismatches in alignments to both genomes. Aligned reads were deduplicated using SAMtools (version 1.10) with standard settings. Uniquely mapped reads were counted on genomic regions using the function qCount (mapqMin = 255L) from QuasR package (version 1.36.0). log2RPKM values were calculated for each tile and normalized between biological samples of mutant and control using the function normalizeBetweenArrays (method = "cyclicloess",cyclic.method = "fast") from the package limma (version 3.52.2). For allelic analysis, total number of maternal and paternal reads was used as library size for calculating RPKM values. Normalized log2RPKM values were transformed back to read counts and were processed by edgeR to identify differential enrichment of histone modification on tiles among experimental groups and calculate log2 fold changes with standard parameters (prior.count = 1). Statistical significance cutoffs for a tile to be considered as differentially enriched were set as follows: log2FC > 1 or < −1 and P-Value < 0.05. For heatmap plots the uniquely mapped reads were quantified 5kb upstream and downstream of the middle of each genomic tile in 51bp bins using the qProfile function from QuasR package. Log2RPKM values of each bin were calculated and normalized as above. We performed smoothening by taking the mean value of 10 bins upstream and downstream of a given bin. The data were plotted using the package ComplexHeatmap (version 2.12.1) or ggplot2 (version 3.3.6).

Nugen Ovation Solo RNA-seq reads were first processed using TrimGalore (version 0.6.2) to trim adaptor and low-quality reads with settings (--clip_R1 3, --stringency 3). Trimmed reads were then aligned to the mouse genome build mm10 using STAR (version 2.5.0a) with settings (--outFilterMismatchNmax 6). Duplicated reads were marked/removed using nudup.py (version 2.2) with settings (-s 8, −l 8). Uniquely mapped reads were counted on genes (TxDb.Mmusculus.UCSC.mm10.knownGene) using the function qCount (mapqMin = 255L) from QuasR package (version 1.36.0). log2RPKM values for each gene were calculated. The resulting read count data were processed by edgeR to identify differentially expressed genes among experimental groups. Genes with CPM< 1 were excluded from the analysis. Genes with log2FC >1 or <-1 and false discovery rate-adjusted FDR < 0.05 were considered to be significantly changed.

Differential expression analysis for early 4-cell mouse embryos RNA-Seq data was done using generalized linear model (GLM) with basis functions for natural splines included into the model to regress out expression differences that might be explained by possible developmental delay. First, we performed pseudotime ordering of knock-out and *Ctrl* embryos using in-house RNA-seq data for several time points of pre-implantation development as a benchmark. Pseudotime for each embryo was estimated using R package SCORPIUS (v1.0.8)(108). Next, we constructed a model matrix for GLM considering genotypes as covariate SmartSeq2 RNA-seq reads were first processed using TrimGalore (version 0.6.2) to trim adaptor and low-quality reads with settings (--stringency 3). Trimmed reads were then aligned to the mouse genome build mm10 using STAR (version 2.5.0a) with settings (-- outFilterMismatchNmax 6). SmartSeq2 RNA-seq samples for hybrid JF1/MsJ x C57BL/6JRj 4-cell embryos were separately aligned to C57BL/6J and JF1/MsJ genomes obtained by incorporating JF1 single-nucleotide polymorphisms (SNPs) into reference mm10 genome using previously published SNP table from (107). Reads were categorized as maternal (JF1/MsJ), paternal (C57BL/6JRj) or undefined based on minimal number of mismatches in alignments to both genomes. Aligned reads were deduplicated using SAMtools (version 1.10) with standard settings. Uniquely mapped reads were counted on genes (TxDb.Mmusculus.UCSC.mm10.ensGene) and tiles using the function qCount (mapqMin = 255L) from QuasR package (version 1.36.0). log2RPKM values for each gene or tile were calculated.

and including basis functions for natural splines with 3 components generated by ns function in R package splines (version 3.5.1) using pseudotime as knots to regress out effects of possible developmental delays. More explicitly, design matrix for GLM was generated using model.matrix function with formula ∼ 0 + genotype + ns(PsT,3). To control possible overfitting by splines, the same model was fit for samples with randomly permuted pseudotime estimates. Expression changes and FDR were calculated for difference between paternal knock-out and control using log-likelihood test and Benjamini-Hochberg method for multiple testing correction. Differential expression analysis for allele specific expression was done similarly, taking read counts for respective allele, and normalizing by total number of allelic reads. In addition, uniquely mapped reads were quantified 5kb upstream and downstream of the middle of each genomic tile in 51bp bins using the qProfile function from QuasR package. Since genomic tiles do not have any particular orientation, we have artificially oriented the tiles to exhibit higher amount of RNA tags on their right side bins. We performed smoothening by taking the mean value of 10 bins upstream and downstream of a given bin.

Bigwig files for all sequencing experiments were generated using qExportWig function from QuasR package. For principal component analysis in Supplementary Figure 5A the package PCAtools (version 2.8.0) was used. Clustering of tiles in Figure 7A was performed with hclust and dist functions from the stats package (version 4.2.1) using the “Ward.D” method and “eucledian” distance. Gene ontology analysis was performed using the Mouse GO slim subset and the package topGO (version 2.48.0).

Analysis of enrichments of GC-poor/GC-rich hypoDMR/nonDMR tiles among tiles with significant changes in particular chromatin mark, e.g. gain or loss of nucleosome ChIP enrichment, was done by constructing 2×2 contingency tables with tile counts belonging/not belonging to particular class of tiles, e.g. GC-poor hypoDMR, versus having/not having significant change in particular chromatin mark and running Fisher’s exact test using R function “fisher.test”. log2-enrichment values were calculated by dividing observed to expected tile counts calculated using R function “chisq.test” (after adding pseudocount 1) and taking log2, i.e. log2(Observed+1)/(Expected+1).

## Data availability

The datasets produced in this study are available in the following database: Gene Expression Omnibus GSE229246 (https://www.ncbi.nlm.nih.gov/geo/query/acc.cgi?acc=GSE229246). Publicly available datasets used in this study can be found at Gene Expression Omnibus database with the following accession IDs: GSE148150 (47), GSE56697 (25) and GSE73952 (66).

## AUTHOR CONTIRBUTIONS

G.F. and A.H.F.M.P. conceived the study, designed the experiments, interpreted the data, and wrote the manuscript with input from all authors. G.F. performed genomic experiments and analyzed the data with the help of E.A.O.. G.F. and S.A.S. developed the ATATA-seq methodology. L.G-.T. and G.F. performed ChIP-seq experiment of sperm samples. P.A.K. and G.F. performed imaging experiments.

## ACKNOWLEDGEMENTS

We thank Y.K. Kawamura for expert advice on embryo culture experiments, M.B. Stadler for advice on computational data analysis and interpretation, H. Kohler for cell sorting, J. Keusch for support in Tn5-ZZ purification, L. Plantard for microscopy and imaging support, members of the animal facility for support, E. Li for *Dnmt3a* and *Dnmt3b* floxed mouse lines and J. van der Vlag for providing antibodies. We thank J. Brind’Amour and M. Lorincz for training on using the ULI-NChIP protocol. We acknowledge M. Gill and other members of the group for feedback on the manuscript. This research was supported by the Novartis Research Foundation, the Swiss National Science Foundation (31003A-172873) and the European Research Council (ERC) under the European Union’s Horizon 2020 research and innovation programme (grant agreement ERC-AdG no. 695288 – Totipotency).

## REFERENCES

1. Deaton, A.M. and Bird, A. (2011) CpG islands and the regulation of transcription. Genes Dev, 25, 1010–1022.

2. Reik, W., Dean, W. and Walter, J. (2001) Epigenetic reprogramming in mammalian development. Science, 293, 1089–1093.

3. Erkek, S., Hisano, M., Liang, C.Y., Gill, M., Murr, R., Dieker, J., Schubeler, D., van der Vlag, J., Stadler, M.B. and Peters, A.H. (2013) Molecular determinants of nucleosome retention at CpG-rich sequences in mouse spermatozoa. Nat Struct Mol Biol, 20, 868–875.

4. Heard, E. and Martienssen, R.A. (2014) Transgenerational epigenetic inheritance: myths and mechanisms. Cell, 157, 95–109.

5. Boskovic, A. and Rando, O.J. (2018) Transgenerational Epigenetic Inheritance. Annu Rev Genet, 52, 21–41.

6. Kobayashi, H., Sakurai, T., Miura, F., Imai, M., Mochiduki, K., Yanagisawa, E., Sakashita, A., Wakai, T., Suzuki, Y., Ito, T. et al. (2013) High-resolution DNA methylome analysis of primordial germ cells identifies gender-specific reprogramming in mice. Genome Res, 23, 616–627.

7. Kubo, N., Toh, H., Shirane, K., Shirakawa, T., Kobayashi, H., Sato, T., Sone, H., Sato, Y., Tomizawa, S., Tsurusaki, Y. et al. (2015) DNA methylation and gene expression dynamics during spermatogonial stem cell differentiation in the early postnatal mouse testis. BMC Genomics, 16, 624.

8. Seisenberger, S., Andrews, S., Krueger, F., Arand, J., Walter, J., Santos, F., Popp, C., Thienpont, B., Dean, W. and Reik, W. (2012) The dynamics of genome-wide DNA methylation reprogramming in mouse primordial germ cells. Mol Cell, 48, 849–862.

9. Mattei, A.L., Bailly, N. and Meissner, A. (2022) DNA methylation: a historical perspective. Trends Genet, 38, 676–707.

10. Dura, M., Teissandier, A., Armand, M., Barau, J., Lapoujade, C., Fouchet, P., Bonneville, L., Schulz, M., Weber, M., Baudrin, L.G. et al. (2022) DNMT3A-dependent DNA methylation is required for spermatogonial stem cells to commit to spermatogenesis. Nat Genet, 54, 469–480.

11. Bourc’his, D. and Bestor, T.H. (2004) Meiotic catastrophe and retrotransposon reactivation in male germ cells lacking Dnmt3L. Nature, 431, 96–99.

12. Barau, J., Teissandier, A., Zamudio, N., Roy, S., Nalesso, V., Herault, Y., Guillou, F. and Bourc’his, D. (2016) The DNA methyltransferase DNMT3C protects male germ cells from transposon activity. Science, 354, 909–912.

13. Kaneda, M., Okano, M., Hata, K., Sado, T., Tsujimoto, N., Li, E. and Sasaki, H. (2004) Essential role for de novo DNA methyltransferase Dnmt3a in paternal and maternal imprinting. Nature, 429, 900–903.

14. Davis, T.L., Yang, G.J., McCarrey, J.R. and Bartolomei, M.S. (2000) The H19 methylation imprint is erased and re-established differentially on the parental alleles during male germ cell development. Hum Mol Genet, 9, 2885–2894.

15. Davis, T.L., Trasler, J.M., Moss, S.B., Yang, G.J. and Bartolomei, M.S. (1999) Acquisition of the H19 methylation imprint occurs differentially on the parental alleles during spermatogenesis. Genomics, 58, 18–28.

16. Kobayashi, H., Sakurai, T., Imai, M., Takahashi, N., Fukuda, A., Yayoi, O., Sato, S., Nakabayashi, K., Hata, K., Sotomaru, Y. et al. (2012) Contribution of intragenic DNA methylation in mouse gametic DNA methylomes to establish oocyte-specific heritable marks. PLoS Genet, 8, e1002440.

17. Gaspa-Toneu, L. and Peters, A.H. (2023) Nucleosomes in mammalian sperm: conveying paternal epigenetic inheritance or subject to reprogramming between generations? Curr Opin Genet Dev, 79, 102034.

18. Balhorn, R., Gledhill, B.L. and Wyrobek, A.J. (1977) Mouse sperm chromatin proteins: quantitative isolation and partial characterization. Biochemistry, 16, 4074–4080.

19. Hammoud, S.S., Nix, D.A., Zhang, H., Purwar, J., Carrell, D.T. and Cairns, B.R. (2009) Distinctive chromatin in human sperm packages genes for embryo development. Nature, 460, 473–478.

20. Brykczynska, U., Hisano, M., Erkek, S., Ramos, L., Oakeley, E.J., Roloff, T.C., Beisel, C., Schubeler, D., Stadler, M.B. and Peters, A.H. (2010) Repressive and active histone methylation mark distinct promoters in human and mouse spermatozoa. Nat Struct Mol Biol, 17, 679–687.

21. Carone, B.R., Hung, J.H., Hainer, S.J., Chou, M.T., Carone, D.M., Weng, Z., Fazzio, T.G. and Rando, O.J. (2014) High-resolution mapping of chromatin packaging in mouse embryonic stem cells and sperm. Dev Cell, 30, 11–22.

22. Yamaguchi, K., Hada, M., Fukuda, Y., Inoue, E., Makino, Y., Katou, Y., Shirahige, K. and Okada, Y. (2018) Re-evaluating the Localization of Sperm-Retained Histones Revealed the Modification-Dependent Accumulation in Specific Genome Regions. Cell Rep, 23, 3920–3932.

23. Yin, Q., Yang, C.H., Strelkova, O.S., Wu, J., Sun, Y., Gopalan, S., Yang, L., Dekker, J., Fazzio, T.G., Li, X.Z. et al. (2023) Revisiting chromatin packaging in mouse sperm. Genome Res, 33, 2079–2093.

24. Hisano, M., Erkek, S., Dessus-Babus, S., Ramos, L., Stadler, M.B. and Peters, A.H. (2013) Genome-wide chromatin analysis in mature mouse and human spermatozoa. Nat Protoc, 8, 2449–2470.

25. Wang, L., Zhang, J., Duan, J., Gao, X., Zhu, W., Lu, X., Yang, L., Zhang, J., Li, G., Ci, W. et al. (2014) Programming and inheritance of parental DNA methylomes in mammals. Cell, 157, 979–991.

26. Aoshima, K., Inoue, E., Sawa, H. and Okada, Y. (2015) Paternal H3K4 methylation is required for minor zygotic gene activation and early mouse embryonic development. EMBO Rep, 16, 803–812.

27. Ferguson-Smith, A.C. (2011) Genomic imprinting: the emergence of an epigenetic paradigm. Nat Rev Genet, 12, 565–575.

28. Inoue, A., Jiang, L., Lu, F., Suzuki, T. and Zhang, Y. (2017) Maternal H3K27me3 controls DNA methylation-independent imprinting. Nature, 547, 419–424.

29. Lismer, A., Siklenka, K., Lafleur, C., Dumeaux, V. and Kimmins, S. (2020) Sperm histone H3 lysine 4 trimethylation is altered in a genetic mouse model of transgenerational epigenetic inheritance. Nucleic Acids Res, 48, 11380–11393.

30. Ben Maamar, M., Sadler-Riggleman, I., Beck, D. and Skinner, M.K. (2018) Epigenetic Transgenerational Inheritance of Altered Sperm Histone Retention Sites. Sci Rep, 8, 5308.

31. Ihara, M., Meyer-Ficca, M.L., Leu, N.A., Rao, S., Li, F., Gregory, B.D., Zalenskaya, I.A., Schultz, R.M. and Meyer, R.G. (2014) Paternal poly (ADP-ribose) metabolism modulates retention of inheritable sperm histones and early embryonic gene expression. PLoS Genet, 10, e1004317.

32. Lismer, A., Dumeaux, V., Lafleur, C., Lambrot, R., Brind’Amour, J., Lorincz, M.C. and Kimmins, S. (2021) Histone H3 lysine 4 trimethylation in sperm is transmitted to the embryo and associated with diet-induced phenotypes in the offspring. Dev Cell, 56, 671–686 e676.

33. Siklenka, K., Erkek, S., Godmann, M., Lambrot, R., McGraw, S., Lafleur, C., Cohen, T., Xia, J., Suderman, M., Hallett, M. et al. (2015) Disruption of histone methylation in developing sperm impairs offspring health transgenerationally. Science, 350, aab2006.

34. Yoshida, K., Muratani, M., Araki, H., Miura, F., Suzuki, T., Dohmae, N., Katou, Y., Shirahige, K., Ito, T. and Ishii, S. (2018) Mapping of histone-binding sites in histone replacement-completed spermatozoa. Nat Commun, 9, 3885.

35. Karahan, G., Chan, D., Shirane, K., McClatchie, T., Janssen, S., Baltz, J.M., Lorincz, M. and Trasler, J. (2021) Paternal MTHFR deficiency leads to hypomethylation of young retrotransposons and reproductive decline across two successive generations. Development, 148.

36. Aarabi, M., Christensen, K.E., Chan, D., Leclerc, D., Landry, M., Ly, L., Rozen, R. and Trasler, J. (2018) Testicular MTHFR deficiency may explain sperm DNA hypomethylation associated with high dose folic acid supplementation. Hum Mol Genet, 27, 1123–1135.

37. Ooi, S.K., Qiu, C., Bernstein, E., Li, K., Jia, D., Yang, Z., Erdjument-Bromage, H., Tempst, P., Lin, S.P., Allis, C.D. et al. (2007) DNMT3L connects unmethylated lysine 4 of histone H3 to de novo methylation of DNA. Nature, 448, 714–717.

38. Otani, J., Nankumo, T., Arita, K., Inamoto, S., Ariyoshi, M. and Shirakawa, M. (2009) Structural basis for recognition of H3K4 methylation status by the DNA methyltransferase 3A ATRX-DNMT3-DNMT3L domain. EMBO Rep, 10, 1235–1241.

39. Stewart, K.R., Veselovska, L., Kim, J., Huang, J., Saadeh, H., Tomizawa, S., Smallwood, S.A., Chen, T. and Kelsey, G. (2015) Dynamic changes in histone modifications precede de novo DNA methylation in oocytes. Genes Dev, 29, 2449–2462.

40. Zhang, Y., Jurkowska, R., Soeroes, S., Rajavelu, A., Dhayalan, A., Bock, I., Rathert, P., Brandt, O., Reinhardt, R., Fischle, W. et al. (2010) Chromatin methylation activity of Dnmt3a and Dnmt3a/3L is guided by interaction of the ADD domain with the histone H3 tail. Nucleic Acids Res, 38, 4246–4253.

41. Kato, Y., Kaneda, M., Hata, K., Kumaki, K., Hisano, M., Kohara, Y., Okano, M., Li, E., Nozaki, M. and Sasaki, H. (2007) Role of the Dnmt3 family in de novo methylation of imprinted and repetitive sequences during male germ cell development in the mouse. Hum Mol Genet, 16, 2272–2280.

42. Dong, J., Wang, X., Cao, C., Wen, Y., Sakashita, A., Chen, S., Zhang, J., Zhang, Y., Zhou, L., Luo, M. et al. (2019) UHRF1 suppresses retrotransposons and cooperates with PRMT5 and PIWI proteins in male germ cells. Nat Commun, 10, 4705.

43. Takada, Y., Yaman-Deveci, R., Shirakawa, T., Sharif, J., Tomizawa, S.I., Miura, F., Ito, T., Ono, M., Nakajima, K., Koseki, Y. et al. (2021) Maintenance DNA methylation in pre-meiotic germ cells regulates meiotic prophase by facilitating homologous chromosome pairing. Development, 148.

44. Shirakawa, T., Yaman-Deveci, R., Tomizawa, S., Kamizato, Y., Nakajima, K., Sone, H., Sato, Y., Sharif, J., Yamashita, A., Takada-Horisawa, Y. et al. (2013) An epigenetic switch is crucial for spermatogonia to exit the undifferentiated state toward a Kit-positive identity. Development, 140, 3565–3576.

45. Sadate-Ngatchou, P.I., Payne, C.J., Dearth, A.T. and Braun, R.E. (2008) Cre recombinase activity specific to postnatal, premeiotic male germ cells in transgenic mice. Genesis, 46, 738–742.

46. Vaisvila, R., Ponnaluri, V.K.C., Sun, Z., Langhorst, B.W., Saleh, L., Guan, S., Dai, N., Campbell, M.A., Sexton, B.S., Marks, K. et al. (2021) Enzymatic methyl sequencing detects DNA methylation at single-base resolution from picograms of DNA. Genome Res, 31, 1280–1289.

47. Shirane, K., Miura, F., Ito, T. and Lorincz, M.C. (2020) NSD1-deposited H3K36me2 directs de novo methylation in the mouse male germline and counteracts Polycomb-associated silencing. Nat Genet, 52, 1088–1098.

48. Tokuda, M., Kadokawa, Y., Kurahashi, H. and Marunouchi, T. (2007) CDH1 is a specific marker for undifferentiated spermatogonia in mouse testes. Biol Reprod, 76, 130–141.

49. Shinohara, T., Avarbock, M.R. and Brinster, R.L. (1999) beta1- and alpha6-integrin are surface markers on mouse spermatogonial stem cells. Proc Natl Acad Sci U S A, 96, 5504–5509.

50. Yoshinaga, K., Nishikawa, S., Ogawa, M., Hayashi, S., Kunisada, T., Fujimoto, T. and Nishikawa, S. (1991) Role of c-kit in mouse spermatogenesis: identification of spermatogonia as a specific site of c-kit expression and function. Development, 113, 689–699.

51. Weinberg, D.N., Papillon-Cavanagh, S., Chen, H., Yue, Y., Chen, X., Rajagopalan, K.N., Horth, C., McGuire, J.T., Xu, X., Nikbakht, H. et al. (2019) The histone mark H3K36me2 recruits DNMT3A and shapes the intergenic DNA methylation landscape. Nature, 573, 281–286.

52. Baubec, T., Colombo, D.F., Wirbelauer, C., Schmidt, J., Burger, L., Krebs, A.R., Akalin, A. and Schubeler, D. (2015) Genomic profiling of DNA methyltransferases reveals a role for DNMT3B in genic methylation. Nature, 520, 243–247.

53. Dhayalan, A., Rajavelu, A., Rathert, P., Tamas, R., Jurkowska, R.Z., Ragozin, S. and Jeltsch, A. (2010) The Dnmt3a PWWP domain reads histone 3 lysine 36 trimethylation and guides DNA methylation. J Biol Chem, 285, 26114–26120.

54. Rondelet, G., Dal Maso, T., Willems, L. and Wouters, J. (2016) Structural basis for recognition of histone H3K36me3 nucleosome by human de novo DNA methyltransferases 3A and 3B. J Struct Biol, 194, 357–367.

55. Lerner, A.M., Hepperla, A.J., Keele, G.R., Meriesh, H.A., Yumerefendi, H., Restrepo, D., Zimmerman, S., Bear, J.E., Kuhlman, B., Davis, I.J. et al. (2020) An optogenetic switch for the Set2 methyltransferase provides evidence for transcription-dependent and -independent dynamics of H3K36 methylation. Genome Res, 30, 1605–1617.

56. Xu, Q., Xiang, Y., Wang, Q., Wang, L., Brind’Amour, J., Bogutz, A.B., Zhang, Y., Zhang, B., Yu, G., Xia, W. et al. (2019) SETD2 regulates the maternal epigenome, genomic imprinting and embryonic development. Nat Genet, 51, 844–856.

57. Brind’Amour, J., Liu, S., Hudson, M., Chen, C., Karimi, M.M. and Lorincz, M.C. (2015) An ultra-low-input native ChIP-seq protocol for genome-wide profiling of rare cell populations. Nat Commun, 6, 6033.

58. Kramers, K., Stemmer, C., Monestier, M., van Bruggen, M.C., Rijke-Schilder, T.P., Hylkema, M.N., Smeenk, R.J., Muller, S. and Berden, J.H. (1996) Specificity of monoclonal anti-nucleosome auto-antibodies derived from lupus mice. J Autoimmun, 9, 723–729.

59. Gahurova, L., Tomizawa, S.I., Smallwood, S.A., Stewart-Morgan, K.R., Saadeh, H., Kim, J., Andrews, S.R., Chen, T. and Kelsey, G. (2017) Transcription and chromatin determinants of de novo DNA methylation timing in oocytes. Epigenetics Chromatin, 10, 25.

60. Meissner, A., Mikkelsen, T.S., Gu, H., Wernig, M., Hanna, J., Sivachenko, A., Zhang, X., Bernstein, B.E., Nusbaum, C., Jaffe, D.B. et al. (2008) Genome-scale DNA methylation maps of pluripotent and differentiated cells. Nature, 454, 766–770.

61. Singh, P., Li, A.X., Tran, D.A., Oates, N., Kang, E.R., Wu, X. and Szabo, P.E. (2013) De novo DNA methylation in the male germ line occurs by default but is excluded at sites of H3K4 methylation. Cell Rep, 4, 205–219.

62. Birke, M., Schreiner, S., Garcia-Cuellar, M.P., Mahr, K., Titgemeyer, F. and Slany, R.K. (2002) The MT domain of the proto-oncoprotein MLL binds to CpG-containing DNA and discriminates against methylation. Nucleic Acids Res, 30, 958–965.

63. Lee, J.H., Voo, K.S. and Skalnik, D.G. (2001) Identification and characterization of the DNA binding domain of CpG-binding protein. J Biol Chem, 276, 44669–44676.

64. Kaya-Okur, H.S., Wu, S.J., Codomo, C.A., Pledger, E.S., Bryson, T.D., Henikoff, J.G., Ahmad, K. and Henikoff, S. (2019) CUT&Tag for efficient epigenomic profiling of small samples and single cells. Nat Commun, 10, 1930.

65. Liu, B., Xu, Q., Wang, Q., Feng, S., Lai, F., Wang, P., Zheng, F., Xiang, Y., Wu, J., Nie, J. et al. (2020) The landscape of RNA Pol II binding reveals a stepwise transition during ZGA. Nature, 587, 139–144.

66. Liu, X., Wang, C., Liu, W., Li, J., Li, C., Kou, X., Chen, J., Zhao, Y., Gao, H., Wang, H. et al. (2016) Distinct features of H3K4me3 and H3K27me3 chromatin domains in pre-implantation embryos. Nature, 537, 558–562.

67. Messerschmidt, D.M., Knowles, B.B. and Solter, D. (2014) DNA methylation dynamics during epigenetic reprogramming in the germline and preimplantation embryos. Genes Dev, 28, 812–828.

68. Amouroux, R., Nashun, B., Shirane, K., Nakagawa, S., Hill, P.W., D’Souza, Z., Nakayama, M., Matsuda, M., Turp, A., Ndjetehe, E. et al. (2016) De novo DNA methylation drives 5hmC accumulation in mouse zygotes. Nat Cell Biol, 18, 225–233.

69. Mayer, W., Niveleau, A., Walter, J., Fundele, R. and Haaf, T. (2000) Demethylation of the zygotic paternal genome. Nature, 403, 501–502.

70. Zhang, B., Zheng, H., Huang, B., Li, W., Xiang, Y., Peng, X., Ming, J., Wu, X., Zhang, Y., Xu, Q. et al. (2016) Allelic reprogramming of the histone modification H3K4me3 in early mammalian development. Nature, 537, 553–557.

71. Dahl, J.A., Jung, I., Aanes, H., Greggains, G.D., Manaf, A., Lerdrup, M., Li, G., Kuan, S., Li, B., Lee, A.Y. et al. (2016) Broad histone H3K4me3 domains in mouse oocytes modulate maternal-to-zygotic transition. Nature, 537, 548–552.

72. Hanna, C.W., Taudt, A., Huang, J., Gahurova, L., Kranz, A., Andrews, S., Dean, W., Stewart, A.F., Colome-Tatche, M. and Kelsey, G. (2018) MLL2 conveys transcription-independent H3K4 trimethylation in oocytes. Nat Struct Mol Biol, 25, 73–82.

73. Hanna, C.W., Huang, J., Belton, C., Reinhardt, S., Dahl, A., Andrews, S., Stewart, A.F., Kranz, A. and Kelsey, G. (2022) Loss of histone methyltransferase SETD1B in oogenesis results in the redistribution of genomic histone 3 lysine 4 trimethylation. Nucleic Acids Res, 50, 1993–2004.

74. Yu, C., Fan, X., Sha, Q.Q., Wang, H.H., Li, B.T., Dai, X.X., Shen, L., Liu, J., Wang, L., Liu, K. et al. (2017) CFP1 Regulates Histone H3K4 Trimethylation and Developmental Potential in Mouse Oocytes. Cell Rep, 20, 1161–1172.

75. Andreu-Vieyra, C.V., Chen, R., Agno, J.E., Glaser, S., Anastassiadis, K., Stewart, A.F. and Matzuk, M.M. (2010) MLL2 is required in oocytes for bulk histone 3 lysine 4 trimethylation and transcriptional silencing. PLoS Biol, 8.

76. Pepin, A.S., Lafleur, C., Lambrot, R., Dumeaux, V. and Kimmins, S. (2022) Sperm histone H3 lysine 4 tri-methylation serves as a metabolic sensor of paternal obesity and is associated with the inheritance of metabolic dysfunction. Mol Metab, 59, 101463.

77. Brown, D.A., Di Cerbo, V., Feldmann, A., Ahn, J., Ito, S., Blackledge, N.P., Nakayama, M., McClellan, M., Dimitrova, E., Turberfield, A.H., et al. (2017) The SET1 Complex Selects Actively Transcribed Target Genes via Multivalent Interaction with CpG Island Chromatin. Cell Rep, 20, 2313–2327.

78. Clouaire, T., Webb, S., Skene, P., Illingworth, R., Kerr, A., Andrews, R., Lee, J.H., Skalnik, D. and Bird, A. (2012) Cfp1 integrates both CpG content and gene activity for accurate H3K4me3 deposition in embryonic stem cells. Genes Dev, 26, 1714–1728.

79. Thomson, J.P., Skene, P.J., Selfridge, J., Clouaire, T., Guy, J., Webb, S., Kerr, A.R., Deaton, A., Andrews, R., James, K.D. et al. (2010) CpG islands influence chromatin structure via the CpG-binding protein Cfp1. Nature, 464, 1082–1086.

80. Voo, K.S., Carlone, D.L., Jacobsen, B.M., Flodin, A. and Skalnik, D.G. (2000) Cloning of a mammalian transcriptional activator that binds unmethylated CpG motifs and shares a CXXC domain with DNA methyltransferase, human trithorax, and methyl-CpG binding domain protein 1. Mol Cell Biol, 20, 2108–2121.

81. Ly, L., Chan, D., Aarabi, M., Landry, M., Behan, N.A., MacFarlane, A.J. and Trasler, J. (2017) Intergenerational impact of paternal lifetime exposures to both folic acid deficiency and supplementation on reproductive outcomes and imprinted gene methylation. Mol Hum Reprod, 23, 461–477.

82. Chan, D., Ly, L., Rebolledo, E.M.D., Martel, J., Landry, M., Scott-Boyer, M.P., Droit, A. and Trasler, J.M. (2023) Transgenerational impact of grand-paternal lifetime exposures to both folic acid deficiency and supplementation on genome-wide DNA methylation in male germ cells. Andrology, 11, 927–942.

83. Murphy, P.J., Guo, J., Jenkins, T.G., James, E.R., Hoidal, J.R., Huecksteadt, T., Broberg, D.S., Hotaling, J.M., Alonso, D.F., Carrell, D.T. et al. (2020) NRF2 loss recapitulates heritable impacts of paternal cigarette smoke exposure. PLoS Genet, 16, e1008756.

84. Yoshizaki, K., Kimura, R., Kobayashi, H., Oki, S., Kikkawa, T., Mai, L., Koike, K., Mochizuki, K., Inada, H., Matsui, Y. et al. (2021) Paternal age affects offspring via an epigenetic mechanism involving REST/NRSF. EMBO Rep, 22, e51524.

85. McGann, J.C., Oyer, J.A., Garg, S., Yao, H., Liu, J., Feng, X., Liao, L., Yates, J.R., 3rd and Mandel, G. (2014) Polycomb- and REST-associated histone deacetylases are independent pathways toward a mature neuronal phenotype. Elife, 3, e04235.

86. Stadler, M.B., Murr, R., Burger, L., Ivanek, R., Lienert, F., Scholer, A., van Nimwegen, E., Wirbelauer, C., Oakeley, E.J., Gaidatzis, D., et al. (2011) DNA-binding factors shape the mouse methylome at distal regulatory regions. Nature, 480, 490–495.

87. Huang, G., Liu, L., Wang, H., Gou, M., Gong, P., Tian, C., Deng, W., Yang, J., Zhou, T.T., Xu, G.L. et al. (2020) Tet1 Deficiency Leads to Premature Reproductive Aging by Reducing Spermatogonia Stem Cells and Germ Cell Differentiation. iScience, 23, 100908.

88. Ciccone, D.N., Su, H., Hevi, S., Gay, F., Lei, H., Bajko, J., Xu, G., Li, E. and Chen, T. (2009) KDM1B is a histone H3K4 demethylase required to establish maternal genomic imprints. Nature, 461, 415–418.

89. Fang, R., Chen, F., Dong, Z., Hu, D., Barbera, A.J., Clark, E.A., Fang, J., Yang, Y., Mei, P., Rutenberg, M. et al. (2013) LSD2/KDM1B and its cofactor NPAC/GLYR1 endow a structural and molecular model for regulation of H3K4 demethylation. Mol Cell, 49, 558–570.

90. Xie, L., Pelz, C., Wang, W., Bashar, A., Varlamova, O., Shadle, S. and Impey, S. (2011) KDM5B regulates embryonic stem cell self-renewal and represses cryptic intragenic transcription. EMBO J, 30, 1473–1484.

91. Brenner, C., Luciani, J., Bizet, M., Ndlovu, M., Josseaux, E., Dedeurwaerder, S., Calonne, E., Putmans, P., Cartron, P.F., Defrance, M. et al. (2016) The interplay between the lysine demethylase KDM1A and DNA methyltransferases in cancer cells is cell cycle dependent. Oncotarget, 7, 58939–58952.

92. Wang, J., Hevi, S., Kurash, J.K., Lei, H., Gay, F., Bajko, J., Su, H., Sun, W., Chang, H., Xu, G. et al. (2009) The lysine demethylase LSD1 (KDM1) is required for maintenance of global DNA methylation. Nat Genet, 41, 125–129.

93. Lambrot, R., Lafleur, C. and Kimmins, S. (2015) The histone demethylase KDM1A is essential for the maintenance and differentiation of spermatogonial stem cells and progenitors. FASEB J, 29, 4402–4416.

94. Tomizawa, S.I., Kobayashi, Y., Shirakawa, T., Watanabe, K., Mizoguchi, K., Hoshi, I., Nakajima, K., Nakabayashi, J., Singh, S., Dahl, A. et al. (2018) Kmt2b conveys monovalent and bivalent H3K4me3 in mouse spermatogonial stem cells at germline and embryonic promoters. Development, 145.

95. Samans, B., Yang, Y., Krebs, S., Sarode, G.V., Blum, H., Reichenbach, M., Wolf, E., Steger, K., Dansranjavin, T. and Schagdarsurengin, U. (2014) Uniformity of nucleosome preservation pattern in Mammalian sperm and its connection to repetitive DNA elements. Dev Cell, 30, 23–35.

96. Saitou, M. and Kurimoto, K. (2014) Paternal nucleosomes: are they retained in developmental promoters or gene deserts? Dev Cell, 30, 6–8.

97. Okano, M., Bell, D.W., Haber, D.A. and Li, E. (1999) DNA methyltransferases Dnmt3a and Dnmt3b are essential for de novo methylation and mammalian development. Cell, 99, 247–257.

98. Koide, T., Moriwaki, K., Uchida, K., Mita, A., Sagai, T., Yonekawa, H., Katoh, H., Miyashita, N., Tsuchiya, K., Nielsen, T.J. et al. (1998) A new inbred strain JF1 established from Japanese fancy mouse carrying the classic piebald allele. Mamm Genome, 9, 15–19.

99. Gaysinskaya, V., Soh, I.Y., van der Heijden, G.W. and Bortvin, A. (2014) Optimized flow cytometry isolation of murine spermatocytes. Cytometry A, 85, 556–565.

100. Smallwood, S.A., Tomizawa, S., Krueger, F., Ruf, N., Carli, N., Segonds-Pichon, A., Sato, S., Hata, K., Andrews, S.R. and Kelsey, G. (2011) Dynamic CpG island methylation landscape in oocytes and preimplantation embryos. Nat Genet, 43, 811–814.

101. van der Heijden, G.W., Dieker, J.W., Derijck, A.A., Muller, S., Berden, J.H., Braat, D.D., van der Vlag, J. and de Boer, P. (2005) Asymmetry in histone H3 variants and lysine methylation between paternal and maternal chromatin of the early mouse zygote. Mech Dev, 122, 1008–1022.

102. Picelli, S., Faridani, O.R., Bjorklund, A.K., Winberg, G., Sagasser, S. and Sandberg, R. (2014) Full-length RNA-seq from single cells using Smart-seq2. Nat Protoc, 9, 171–181.

103. Schindelin, J., Arganda-Carreras, I., Frise, E., Kaynig, V., Longair, M., Pietzsch, T., Preibisch, S., Rueden, C., Saalfeld, S., Schmid, B. et al. (2012) Fiji: an open-source platform for biological-image analysis. Nat Methods, 9, 676–682.

104. Amemiya, H.M., Kundaje, A. and Boyle, A.P. (2019) The ENCODE Blacklist: Identification of Problematic Regions of the Genome. Sci Rep, 9, 9354.

105. Strogantsev, R., Krueger, F., Yamazawa, K., Shi, H., Gould, P., Goldman-Roberts, M., McEwen, K., Sun, B., Pedersen, R. and Ferguson-Smith, A.C. (2015) Allele-specific binding of ZFP57 in the epigenetic regulation of imprinted and non-imprinted monoallelic expression. Genome Biol, 16, 112.

106. Chen, Y., Pal, B., Visvader, J.E. and Smyth, G.K. (2017) Differential methylation analysis of reduced representation bisulfite sequencing experiments using edgeR. F1000Res, 6, 2055.

107. Takada, T., Ebata, T., Noguchi, H., Keane, T.M., Adams, D.J., Narita, T., Shin, I.T., Fujisawa, H., Toyoda, A., Abe, K. et al. (2013) The ancestor of extant Japanese fancy mice contributed to the mosaic genomes of classical inbred strains. Genome Res, 23, 1329–1338.

108. Cannoodt, R., Saelens, W., Sichien, D., Tavernier, S., Janssens, S., Guilliams, M., Lambrecht, B., Preter, K.D. and Saeys, Y. (2016). bioRxiv.

