## Supplementary Figures 1 to 15 for "DNA methylation modulates nucleosome retention in sperm and H3K4 methylation deposition in early mouse embryos"

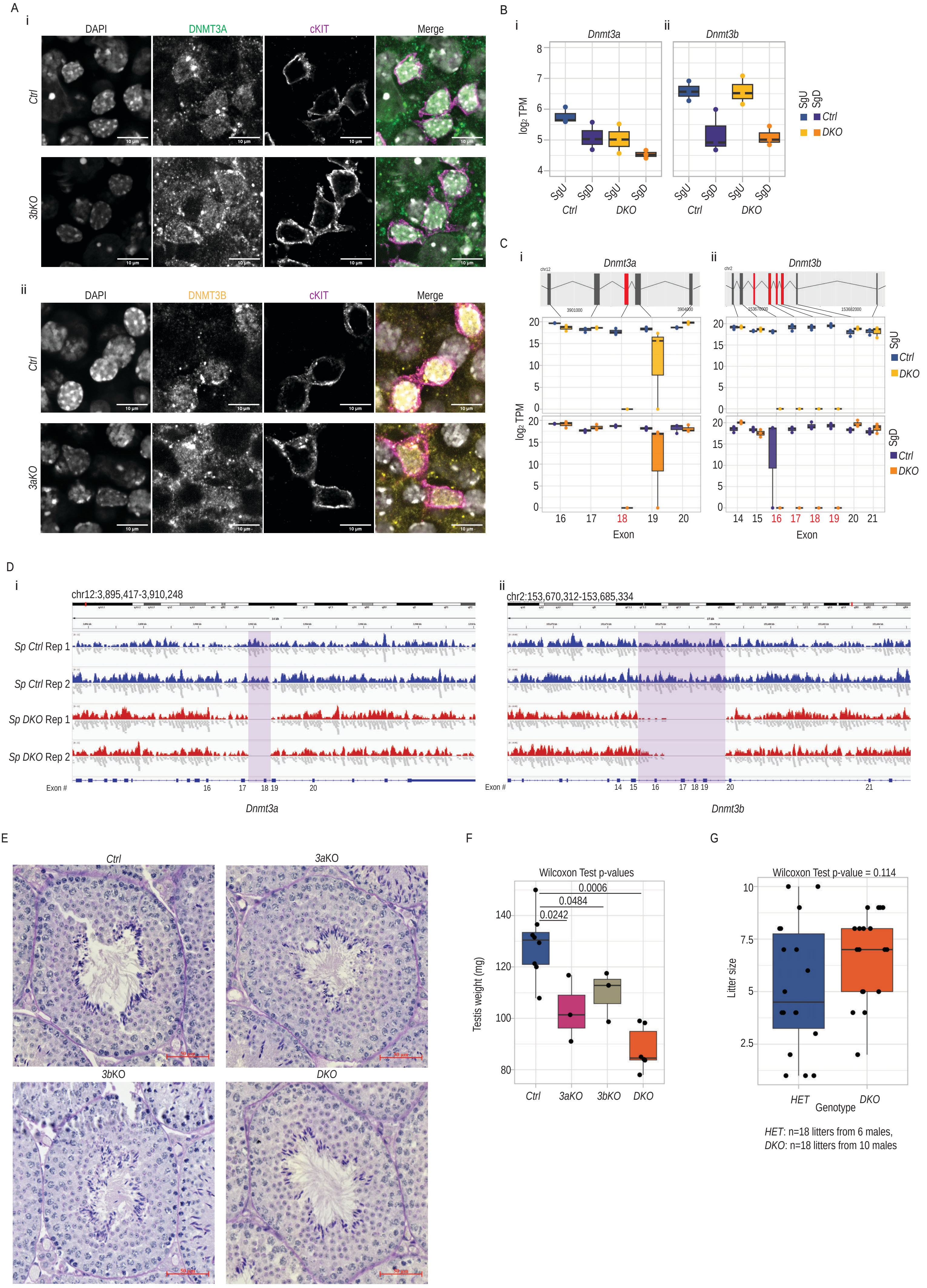
A
i
DAPI
DNMT3A
cKIT
Merge
Ctrl
3bKO
10 µm
ii
DAPI
DNMT3B
cKIT
Merge
Ctrl
3aKO
B
i
Dnmt3a
ii
Dnmt3b
log2 TPM
SgU
SgD
Ctrl
DKO
C
i
Dnmt3a
ii
Dnmt3b
chr12
3901000
3904000
chr2
153670000
153682000
log2 TPM
SgU
SgD
Ctrl
DKO
Exon
D
i
chr12:3,895,417-3,910,248
ii
chr2:153,670,312-153,685,334
Sp Ctrl Rep 1
Sp Ctrl Rep 2
Sp DKO Rep 1
Sp DKO Rep 2
Exon #
Dnmt3a
Dnmt3b
E
Ctrl
3aKO
3bKO
DKO
F
Wilcoxon Test p-values
0.0006
0.0484
0.0242
Testis weight (mg)
Ctrl
3aKO
3bKO
DKO
G
Wilcoxon Test p-value = 0.114
Litter size
HET
DKO
Genotype
HET: n=18 litters from 6 males,
DKO: n=18 litters from 10 males

**Supplementary Figure 1. Characterization of Dnmt3a and/or Dnmt3b conditional depletion in male germ cells.**

A) Whole mount immunofluorescence staining of (i) DNMT3A in seminiferous tubules of *Ctrl* and *3bKO* testes, and (ii) DNMT3B in seminiferous tubules of *Ctrl* and *3aKO* testes. All samples were co-stained for cKIT and DNA was visualized by DAPI. Maximum projections of multiple confocal z-stacks are shown. Scale bars = 10 µm.

B) Boxplots showing the log2 TPM values for the (i) Dnmt3a and (ii) Dnmt3b genes in undifferentiated spermatogonia (SgU) and differentiated spermatogonia (SgD) from *Ctrl* and *DKO* samples (n=3 for each biological sample type).

C) Boxplots showing the log2 RPKM values for the floxed exons (in red) and their surrounding exons (in gray) of (i) Dnmt3a and (ii) Dnmt3b genes in SgU and SgD from *Ctrl* and *DKO* samples (n=3 for each biological sample type).

D) IGV genome browser view of aligned whole genome EM-seq reads across the (i) Dnmt3a and (ii) Dnmt3b genes from 2 replicates of *Ctrl* and 2 replicates of *DKO* sperm samples. The genomic loci flanked by loxP sites are highlighted in purple.

E) Representative staining with periodic acid-Schiff's (PAS) and hematoxylin of stage IV-V seminiferous tubule sections from *Ctrl*, *3aKO*, *3bKO* and *DKO* testes. Scale bars = 50 µm.

F) Boxplots plot showing testicular weight from *Ctrl* (n=8), *3aKO* (n=3), *3bKO* (n=3) and *DKO* (n=6) animals. P-values between biological groups were calculated by Wilcoxon Test.

G) Boxplots plot showing number of pups per litter sired either from Het (n=18 litters from 6 sires) and *DKO* (n=18 litters from 10 sires) animals. P-values between biological groups were calculated by Wilcoxon Test.

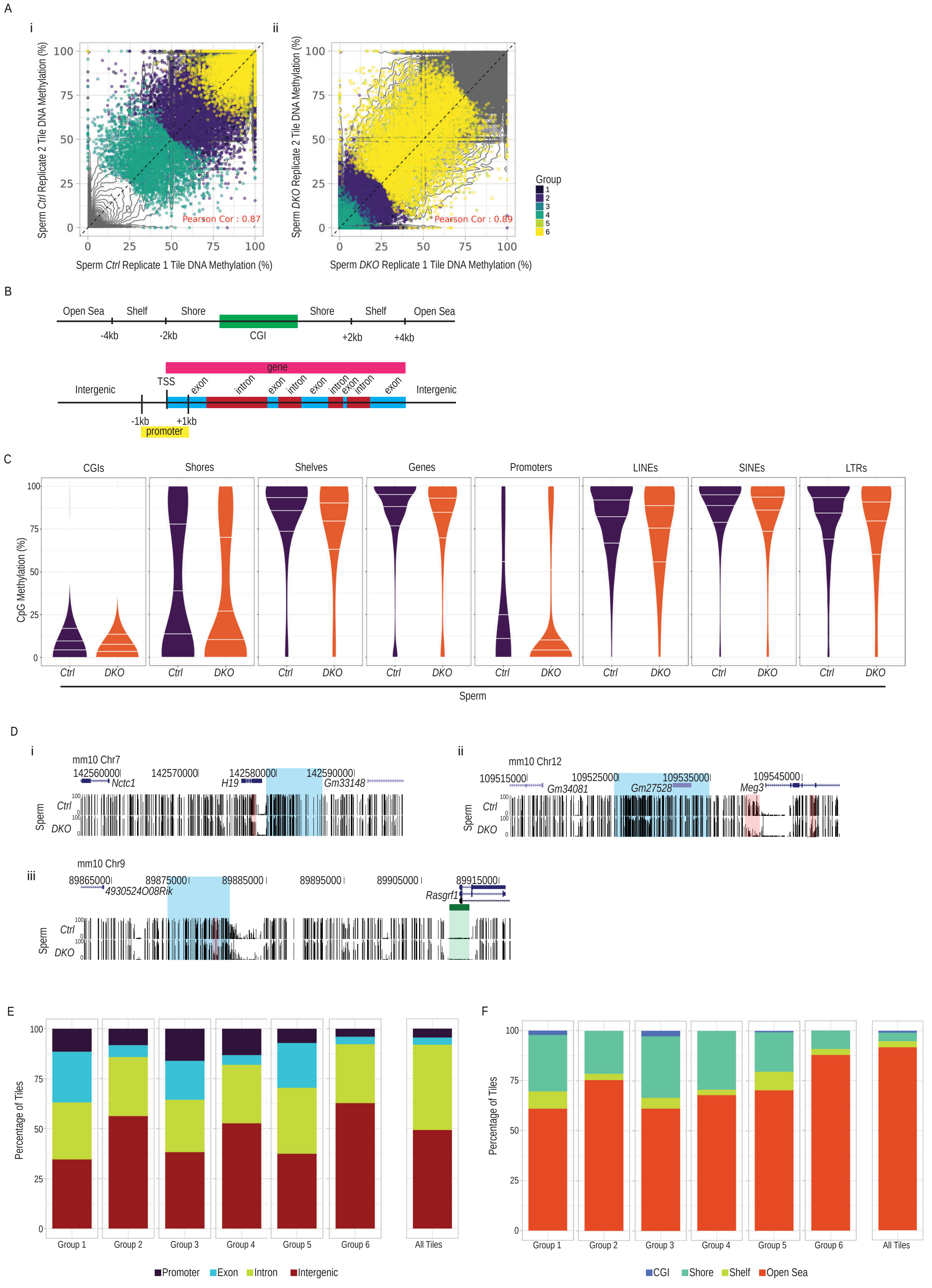
A
i
ii
Sperm Ctrl Replicate 2 Tile DNA Methylation (%)
Sperm Ctrl Replicate 1 Tile DNA Methylation (%)
Pearson Cor : 0.87
Sperm DKO Replicate 2 Tile DNA Methylation (%)
Sperm DKO Replicate 1 Tile DNA Methylation (%)
Pearson Cor : 0.89
Group
1
2
3
4
5
6
B
Open Sea
Shelf
Shore
CGI
Shore
Shelf
Open Sea
-4kb
-2kb
+2kb
+4kb
gene
Intergenic
TSS
exon
intron
Intergenic
-1kb
+1kb
promoter
C
CGIs
Shores
Shelves
Genes
Promoters
LINEs
SINEs
LTRs
CpG Methylation (%)
Ctrl
DKO
Sperm
D
i
mm10 Chr7
142560000
142570000
142580000
142590000
Nctc1
H19
Gm33148
Sperm
Ctrl
DKO
ii
mm10 Chr12
109515000
109525000
109535000
109545000
Gm34081
Gm27528
Meg3
iii
mm10 Chr9
89865000
89875000
89885000
89895000
89905000
89915000
4930524O08Rik
Rasgrf1
E
Percentage of Tiles
Group 1
Group 2
Group 3
Group 4
Group 5
Group 6
All Tiles
Promoter
Exon
Intron
Intergenic
F
Percentage of Tiles
CGI
Shore
Shelf
Open Sea

**Supplementary Figure 2. DNAme of various genomic features and paternal gametic DMRs in *DKO* sperm.**

A) Scatterplot showing percentage DNAme of 500bp genomic tiles assessed by EMseq in (i) replicate 1 *Ctrl* (x-axis) and replicate 2 *Ctrl* (y-axis) sperm and (ii) replicate 1 *DKO* (x-axis) and replicate 2 *DKO* (y-axis) sperm. HypoDMR tiles are highlighted according to the group that they belong according to Figure 1E. Contour lines denote the density of NonDMR tiles. The Pearson correlation coefficient and R-squared values of linear modelling rep1~rep2 for each genotype are presented on upper left corner.

B) Schematic representation of genomic feature annotation related to CpG islands (top) and genes (bottom).

C) Violin plots showing the percentage of DNAme of genomic CpGs that overlap with CGI, gene and repeat related genomic features in *Ctrl* (n=2) and *DKO* (n=2) sperm. The center lines in the violins represent median values. The upper and lower lines indicate the interquartile range (IQR; from the 25th to 75th percentile).

D) UCSC genome browser snapshot of the (i) H19/Igf2, (ii) Dlk1-Gtl2 and (iii) Rasgrf1 paternally methylated gametic DMRs, showing tracks for DNAme levels in *Ctrl* and *DKO* sperm. Gametic DMRs are highlighted in blue. The promoter CGI and the hypoDMR are highlighted in green and red respectively.

E) Bar plot showing the percentage of hypoDMRs in each group as defined in Figure 1E overlapping with gene related features. As a reference, the percentage of gene related features at all genomic tiles is shown on the right.

F) Bar plot showing the percentage of hypoDMRs in each group as defined in Figure 1E overlapping with CpG island related features. As a reference, the percentage of CpG island related features at all genomic tiles is shown on the right.

Fanourgakis et al Supplementary Figure 3

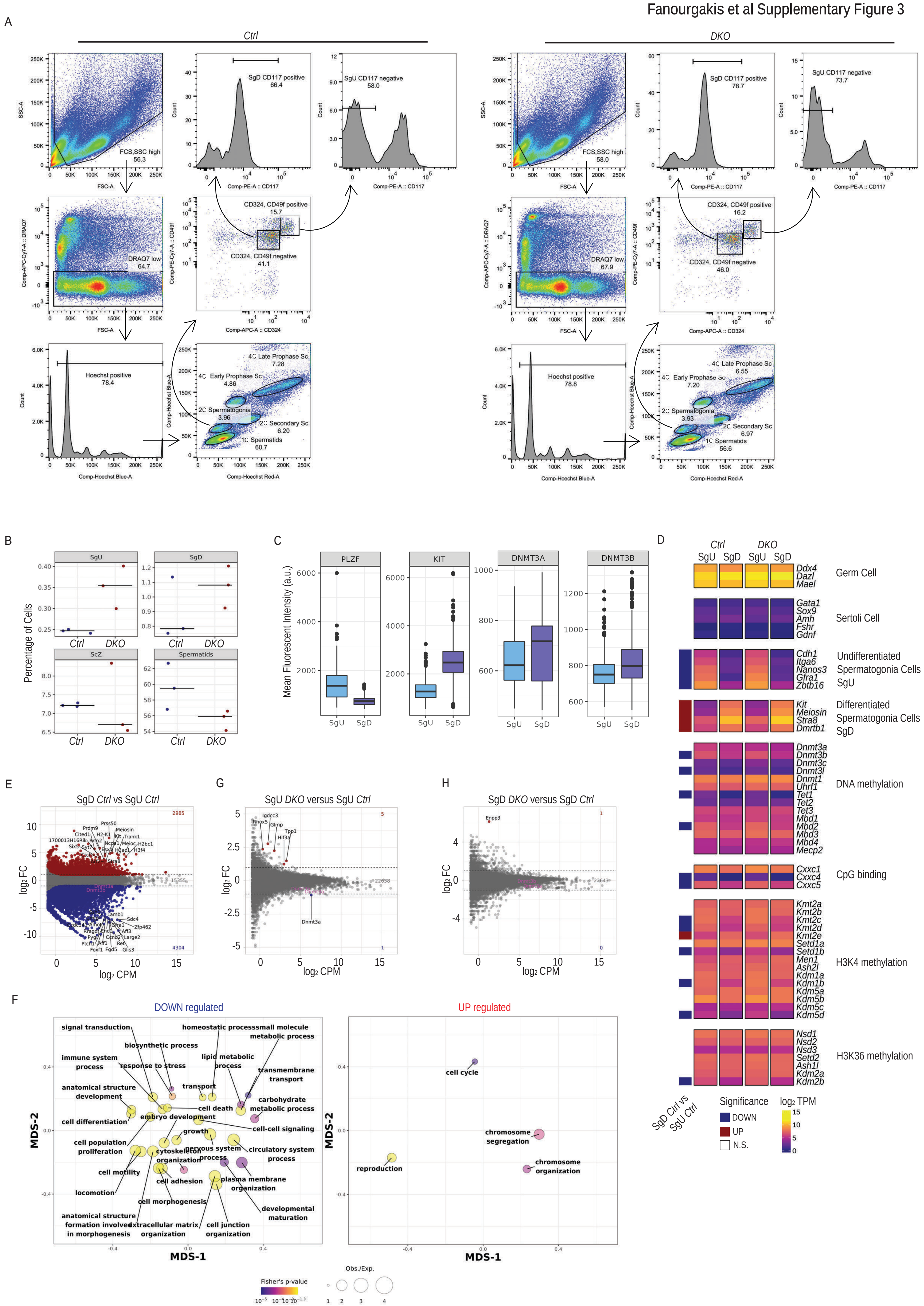

**Supplementary Figure 3. Transcriptional effects of Dnmt3a/Dnmt3b depletion in spermatogonia.**

A) Representative FACS sorting profiles of germ cells with gating parameters from adult *Ctrl* (left) and *DKO* (right) male mice. FSC versus SSC plot indicating gate excluding the debris. FCS versus DRAQ7 plot indicating gate including viable cells. Hoechst-Blue histogram indicate gate with stained cells. Hoechst-Red versus Hoechst-Blue plot indicating gates containing populations of spermatogonia, meiotic spermatocytes and post-meiotic cells. CD324 (E-CADHERIN) versus CD49f (INTEGRIN ALPHA 6) plot showing gates containing undifferentiated spermatogonia SgU (CD324high, CD49fhigh) and differentiated spermatogonia SgD (CD324low, CD49flow). CD117 (c-KIT) histogram indicates the gate with SgU spermatogonia (Hoechst-Bluemid, Hoechst-Redlow, CD324high, CD49fhigh, CD117low). CD117 (c-KIT) histogram indicates the gate with SgD spermatogonia (Hoechst-Bluemid, Hoechst-Redlow, CD324low, CD49flow, CD117high).

B) Dot plots representing the relative percentages of sorted SgU, SgD, Zygotene Spermatocytes (ScZ) and spermatids cells to all viable Hoechst positive cells from *Ctrl* (n=3) and *DKO* (n=3) animals.

C) Boxplots showing from left to right the mean fluorescent intensity value of a-PLZF, a-cKIT, a-DNMT3A and a-DNMT3B immunofluorescence staining of nuclei from FACS sorted SgU and SgD from *Ctrl* animals.

D) Heatmap showing the log2 TPM values of selected gene markers in SgU and SgD from *Ctrl* (n=3) and *DKO* (n=3) samples.

E) Scatter plots showing gene expression log2 fold changes among *Ctrl* SgD (n=3) versus *Ctrl* SgU (n=3) samples were plotted against log2 CPM gene expression levels. The numbers of significantly upregulated (red) or downregulated (blue) genes are displayed on the left and genes names of the most highly significant differentially expressed genes are shown.

F) Bubbleplots showing significantly enriched GO-terms associated with downregulated (left) and upregulated (right) genes in *Ctrl* SgD versus *Ctrl* SgU samples. The size of the bubble denotes enrichment and color denotes significance. Distance of the bubbles centers represents that similarity of genes contained in each GO-term.

G) Scatter plots showing gene expression log2 fold changes among *DKO* SgU (n=3) versus *Ctrl* SgU (n=3) samples were plotted against log2 CPM gene expression levels. The numbers of significantly upregulated (red) or downregulated (blue) genes are displayed on the left and genes names of the most highly significant differentially expressed genes are shown.

H) Scatter plots showing gene expression log2 fold changes among *DKO* SgD (n=3) versus *Ctrl* SgD (n=3) samples were plotted against log2 CPM gene expression levels. The numbers of significantly upregulated (red) or downregulated (blue) genes are displayed on the left and genes names of the most highly significant differentially expressed genes are shown.

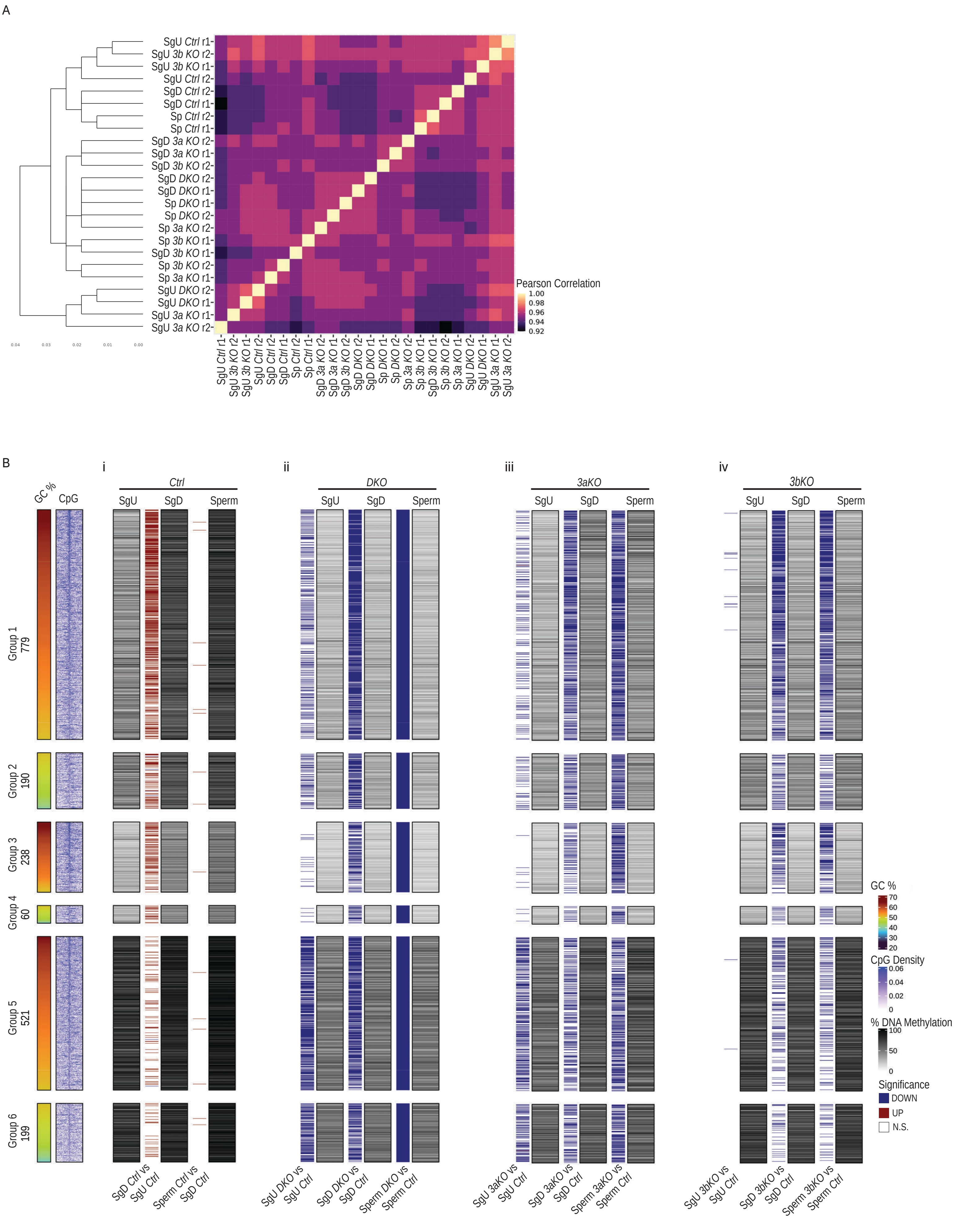
A
Pearson Correlation
1.00
0.98
0.96
0.94
0.92
SgU Ctrl r1
SgU 3b KO r2
SgU 3b KO r1
SgU Ctrl r2
SgD Ctrl r2
SgD Ctrl r1
Sp Ctrl r2
Sp Ctrl r1
SgD 3a KO r2
SgD 3a KO r1
SgD 3b KO r2
SgD DKO r2
SgD DKO r1
Sp DKO r1
Sp DKO r2
Sp 3a KO r2
Sp 3b KO r1
SgD 3b KO r1
Sp 3b KO r2
Sp 3a KO r1
SgU DKO r2
SgU DKO r1
SgU 3a KO r1
SgU 3a KO r2
0.04
0.03
0.02
0.01
0.00
B
i
ii
iii
iv
GC %
CpG
Ctrl
DKO
3aKO
3bKO
SgU
SgD
Sperm
Group 1
779
Group 2
190
Group 3
238
Group 4
60
Group 5
521
Group 6
199
SgD Ctrl vs SgU Ctrl
Sperm Ctrl vs SgD Ctrl
SgU DKO vs SgU Ctrl
SgD DKO vs SgD Ctrl
Sperm DKO vs Sperm Ctrl
SgU 3aKO vs SgU Ctrl
SgD 3aKO vs SgD Ctrl
Sperm 3aKO vs Sperm Ctrl
SgU 3bKO vs SgU Ctrl
SgD 3bKO vs SgD Ctrl
Sperm 3bKO vs Sperm Ctrl
GC %
70
60
50
40
30
20
CpG Density
0.06
0.04
0.02
0
% DNA Methylation
100
50
0
Significance
DOWN
UP
N.S.

**Supplementary Figure 4. DNAme levels of hypoDMRs during spermatogonia development upon Dnmt3a and/or Dnmt3b depletion.**

A) Heatmap showing the Pearson correlation coefficient of DNAme percentage at all genomic tiles detected by RRBS between the biological replicates of undifferentiated spermatogonia (SgU), differentiated spermatogonia (SgD) and sperm from *Ctrl*, *DKO*, *3aKO* and *3bKO* samples.

B) Heatmap showing the DNAme percentage of hypoDMRs detected by RRBS in undifferentiated spermatogonia (SgU), differentiated spermatogonia (SgD) and sperm from *Ctrl*, *DKO*, *3aKO* and *3bKO* samples. Grouping and ordering of hypoDMRs is based on Figure 1E. Dark to bright color scale denotes high to low DNAme percentage. Color bands between the DNAme heatmaps denote hypoDMRs with significantly differential DNAme in the contrasts mentioned below them. The density of CpGs at hypoDMRs (±5 kb from their center) is indicated.

A

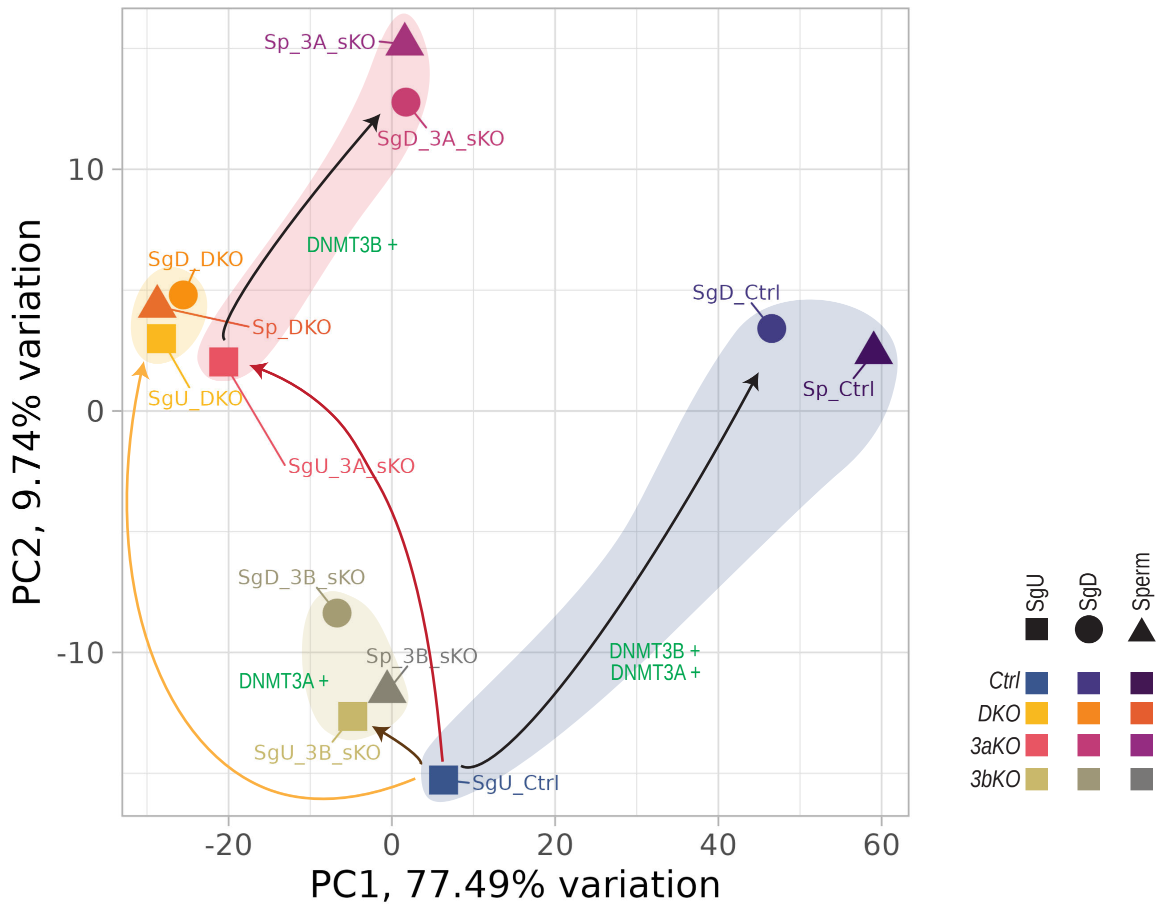
Sp_3A_sKO
SgD_3A_sKO
SgD_DKO
DNMT3B +
Sp_DKO
SgU_DKO
SgU_3A_sKO
SgD_Ctrl
Sp_Ctrl
SgD_3B_sKO
Sp_3B_sKO
DNMT3A +
SgU_3B_sKO
SgU_Ctrl
DNMT3B +
DNMT3A +
PC2, 9.74% variation
PC1, 77.49% variation
SgU
SgD
Sperm
Ctrl
DKO
3aKO
3bKO

B

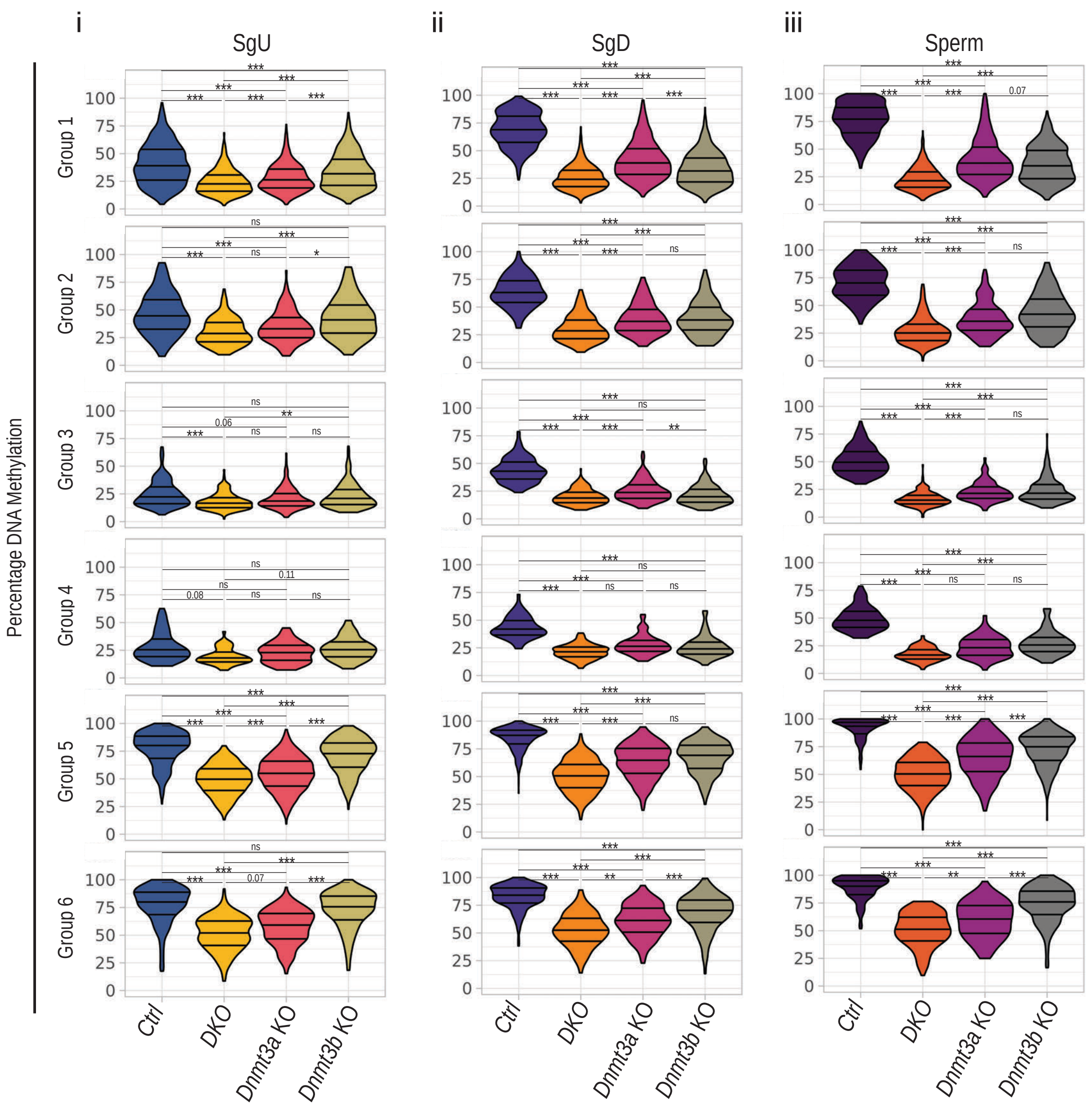
i
SgU
ii
SgD
iii
Sperm
Group 1
Group 2
Group 3
Group 4
Group 5
Group 6
Percentage DNA Methylation
Ctrl
DKO
Dnmt3a KO
Dnmt3b KO

**Supplementary Figure 5. Comparison of DNAme levels of hypoDMRs during spermatogonia development upon Dnmt3a and/or Dnmt3b depletion.**

A) Principal component analysis (PCA) of DNAme levels at hypoDMRS in SgU, SgD and sperm samples from *Ctrl*, *3aKO*, *3bKO* and *DKO* animals.

B) Violin plots showing percentages DNAme of hypoDMRs assessed by RRBS in (i) undifferentiated spermatogonia (SgU) for *Ctrl* (n=2), *3aKO* (n=2), *3bKO* (n=2) and *DKO* (n=2) samples, (ii) differentiated spermatogonia for *Ctrl* (n=2), *3aKO* (n=2), *3bKO* (n=2) and *DKO* (n=2) samples and (iii) sperm for *Ctrl* (n=2), *3aKO* (n=2), *3bKO* (n=2) and *DKO* (n=2) samples. Two-sample Wilcoxon tests were performed between indicated groups * $P < 0.05$, ** $P \leq 0.01$, *** $P \leq 0.001$.

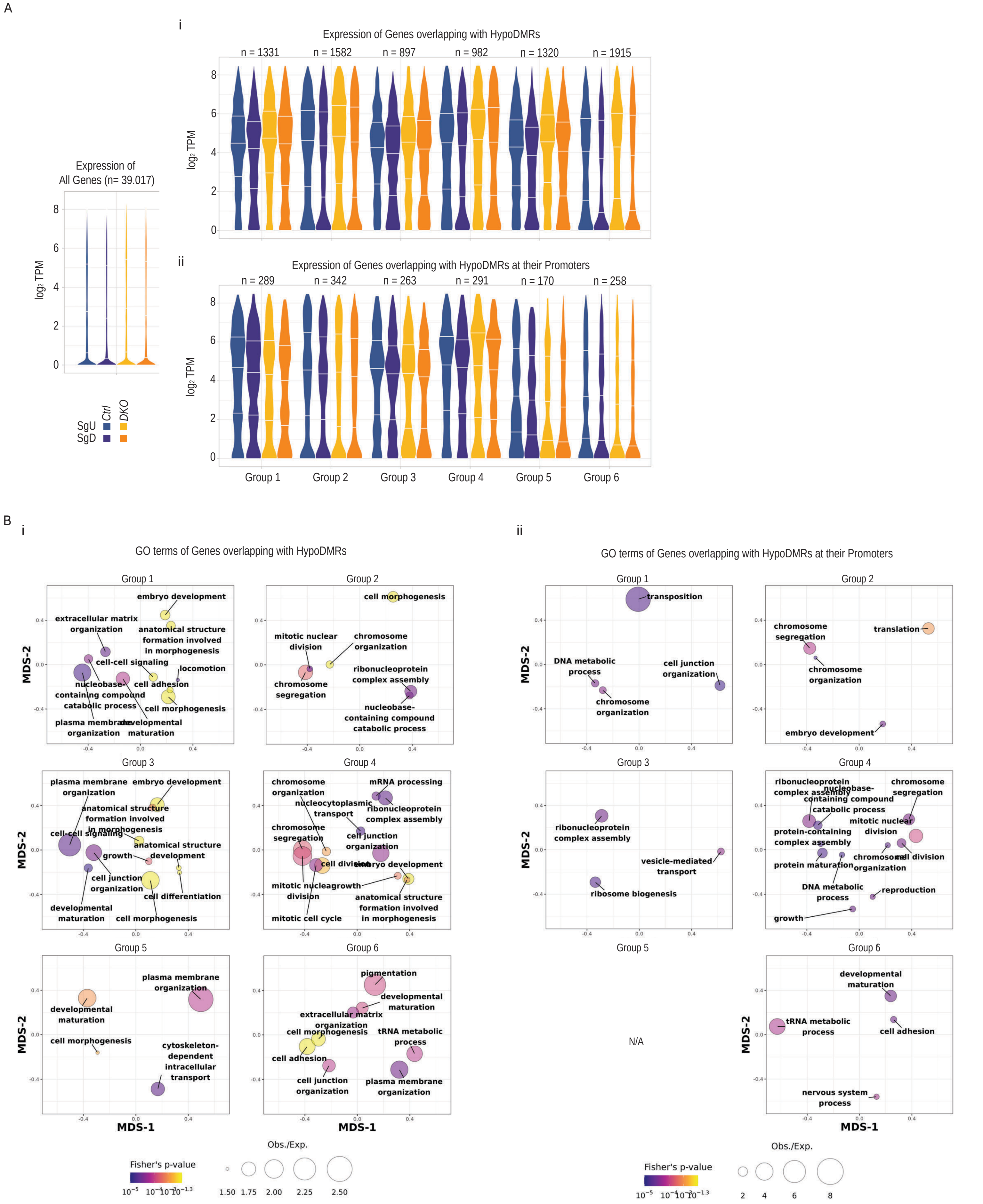
A
i
Expression of Genes overlapping with HypoDMRs
n = 1331
n = 1582
n = 897
n = 982
n = 1320
n = 1915
log2 TPM
Expression of
All Genes (n= 39.017)
ii
Expression of Genes overlapping with HypoDMRs at their Promoters
n = 289
n = 342
n = 263
n = 291
n = 170
n = 258
Group 1
Group 2
Group 3
Group 4
Group 5
Group 6
Ctrl
DKO
SgU
SgD
B
i
GO terms of Genes overlapping with HypoDMRs
ii
GO terms of Genes overlapping with HypoDMRs at their Promoters
MDS-1
MDS-2
embryo development
extracellular matrix organization
anatomical structure formation involved in morphogenesis
cell-cell signaling
locomotion
nucleobase-containing compound catabolic process
cell adhesion
cell morphogenesis
plasma membrane organization
developmental maturation
mitotic nuclear division
chromosome organization
chromosome segregation
ribonucleoprotein complex assembly
nucleocytoplasmic transport
mRNA processing
cell junction organization
growth
anatomical structure development
cell differentiation
cell division
mitotic cell cycle
pigmentation
tRNA metabolic process
cytoskeleton-dependent intracellular transport
transposition
DNA metabolic process
translation
vesicle-mediated transport
ribosome biogenesis
protein-containing complex assembly
protein maturation
reproduction
nervous system process
N/A
Fisher's p-value
Obs./Exp.
1.50
1.75
2.00
2.25
2.50
2
4
6
8

**Supplementary Figure 6. Transcriptional expression and gene ontology of hypoDMRs associated genes.**

A) Violin plot showing log2 TPM values of (i) genes overlapping with hypoDMRs grouped according to Figure 1E and (ii) genes overlapping with promoter associated hypoDMRs grouped according to Figure 1E in *Ctrl* SgU (n=3), *Ctrl* SgD (n=3), *DKO* SgU (n=3) and *DKO* SgD (n=3). For reference log2 TPM values of all genes are shown in the left.

B) Bubbleplots showing significantly enriched GO-terms associated with (i) genes overlapping with hypoDMRs and (ii) genes overlapping with promoter associated hypoDMRs. The size of the bubble denotes enrichment and color denotes significance. Distance of the bubbles centers represents that similarity of genes contained in each GO-term.

Fanourgakis et al Supplementary Figure 7

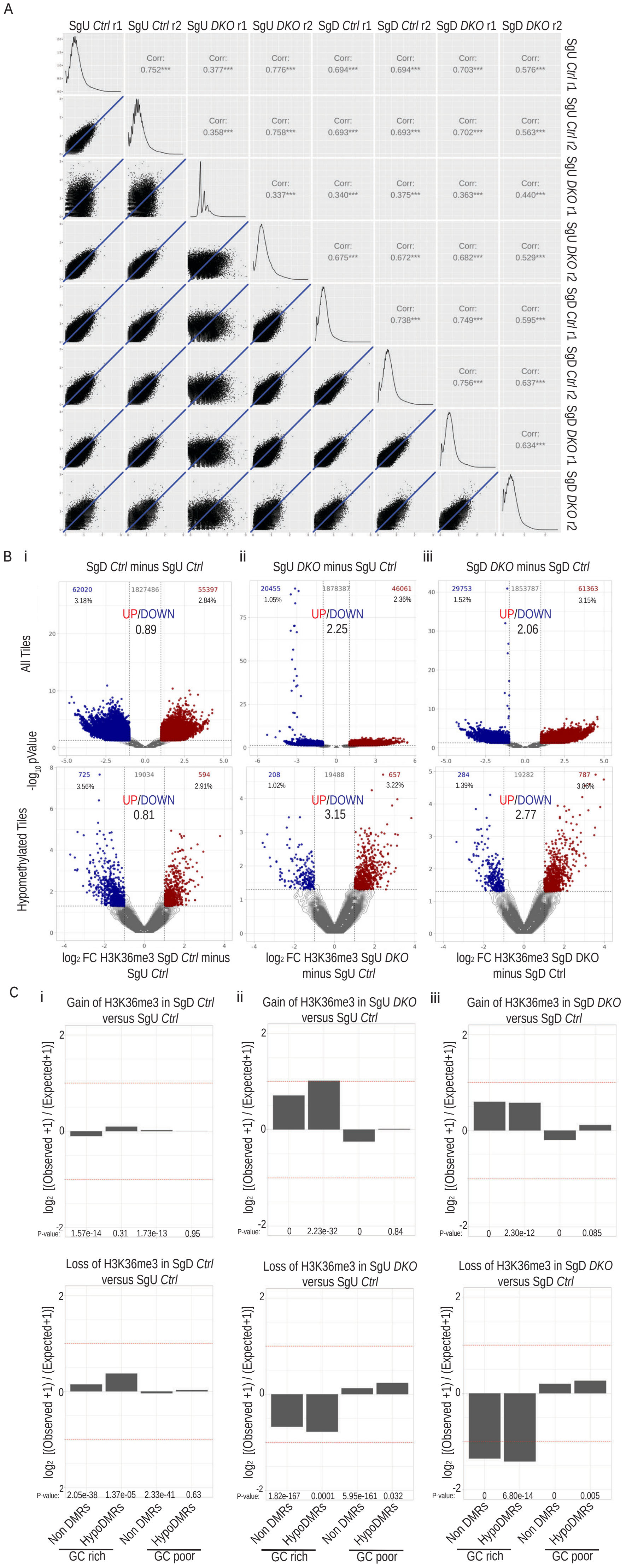

**Supplementary Figure 7. Differential H3K36me3 analysis in *DKO* spermatogonia.**

A) Scatterplots [left side] show the H3K36me3 log2 RPKM values at 100.000 randomly selected genomic tiles detected by ULI-NChIP-seq between pairs of biological replicated samples of undifferentiated spermatogonia (SgU), differentiated spermatogonia (SgD) from *Ctrl* and *DKO* animals. The diagonal line plots show the distribution of the H3K36me3 log2 RPKM values in each replicate. The Pearson correlation coefficient [right side] of H3K36me3 log2 RPKM values at 100.000 randomly selected genomic tiles detected by ULI-NChIP-seq between the biological replicates of undifferentiated spermatogonia (SgU), differentiated spermatogonia (SgD) from *Ctrl* and *DKO* samples.

B) Volcano plots showing differential enrichment for H3K36me3 in (i) *Ctrl* SgD (n=2) versus *Ctrl* SgU (n=2), (ii) *DKO* SgU (n=2) versus *Ctrl* SgU (n=2) and (iii) *DKO* SgD (n=2) versus *Ctrl* SgD (n=2) (right) contrasts for all genomic tiles (upper) and hypoDMR tiles (lower). Tiles with statistically significant gain and loss of the H3K36me3 signals are highlighted in red (upregulated) and blue (downregulated) and their numbers and percentage of differentially are shown on the upper right and left corners respectively. The ratio between upregulated versus downregulated tiles in each plot is shown in the middle.

C) Bar plots showing log2 enrichments of GC-rich or GC-poor nonDMR and hypoDMR tiles among tiles with significant H3K36me3 gain (upper) or significant H3K36me3 loss (lower) in (i) *Ctrl* SgD versus *Ctrl* SgU, (ii) *DKO* SgU versus *Ctrl* SgU and (iii) *DKO* SgD versus *Ctrl* SgD. P-values were calculated by Fisher's exact test.

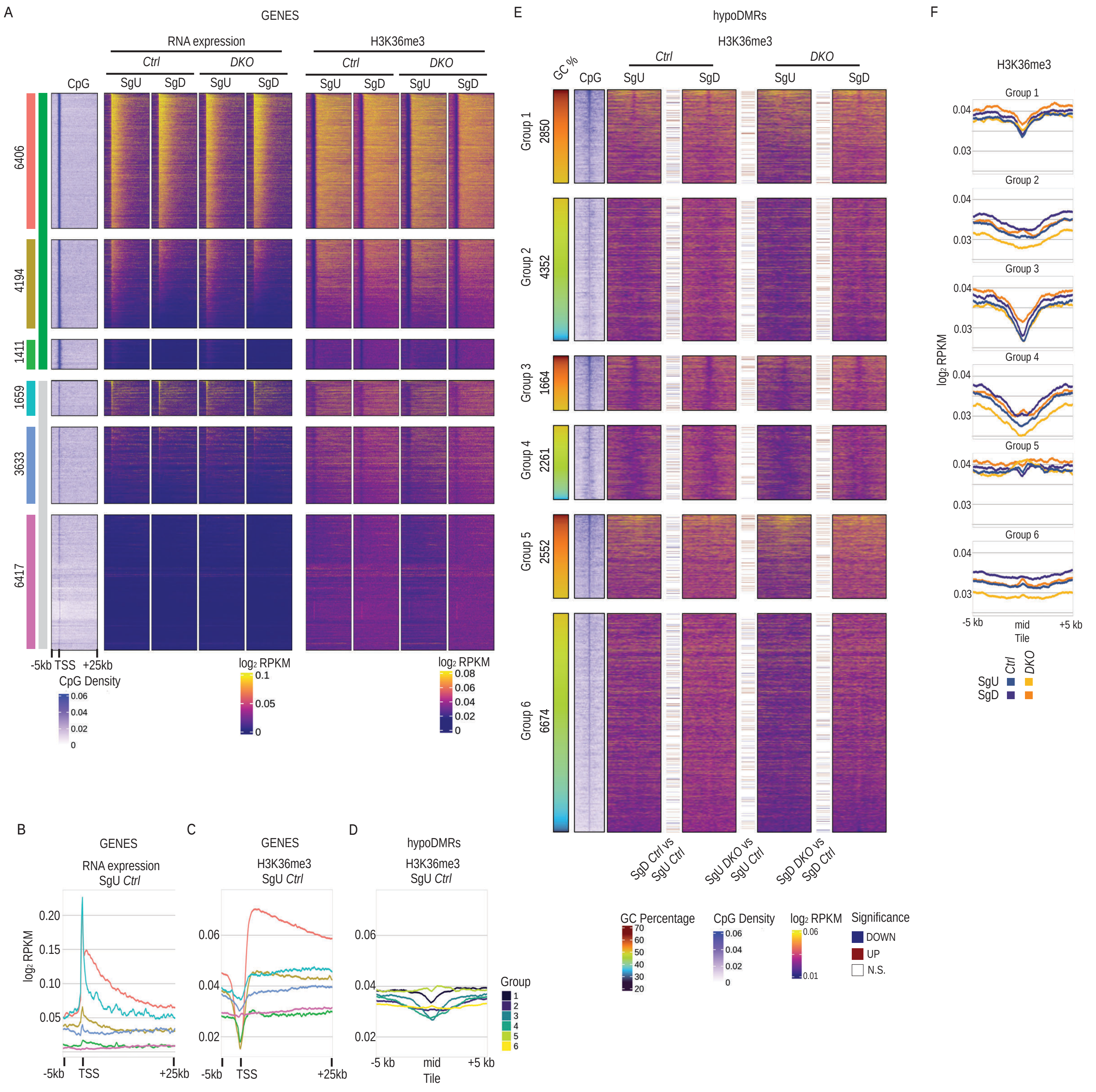
A
GENES
RNA expression
H3K36me3
Ctrl
DKO
CpG
SgU
SgD
6406
4194
1411
1659
3633
6417
-5kb TSS +25kb
CpG Density
0.06
0.04
0.02
0
log2 RPKM
0.1
0.05
0
0.08
0.06
0.04
0.02
0
E
hypoDMRs
H3K36me3
GC %
Group 1 2850
Group 2 4352
Group 3 1664
Group 4 2261
Group 5 2552
Group 6 6674
SgD Ctrl vs SgU Ctrl
SgU DKO vs SgU Ctrl
SgD DKO vs SgD Ctrl
GC Percentage
70
60
50
40
30
20
Significance
DOWN
UP
N.S.
F
Group 1
Group 2
Group 3
Group 4
Group 5
Group 6
log2 RPKM
0.04
0.03
-5 kb
mid
Tile
+5 kb
B
RNA expression
SgU Ctrl
0.20
0.15
0.10
0.05
-5kb
TSS
+25kb
C
D
Group
1
2
3
4
5
6

**Supplementary Figure 8. H3K36me3 levels of hypoDMRs in *DKO* spermatogonia.**

A) Heatmaps showing RNA and H3K36me3 occupancy at gene transcriptional start sites (TSS) (-5/+25kb) in *Ctrl* and *DKO* SgU and SgD (n=2 for each sample). The CpG density at gene TSS (-5/+25kb) is indicated. Genes have been grouped according to the presence of a CpG island at their TSS +/- 1kb and ordered by decreasing expression in *Ctrl* SgU.

B) Line plots showing average RNA occupancy at gene TSS (-5/+25kb) in *Ctrl* SgU (n=2) for each expression group.

C) Line plots showing average H3K36me3 occupancy at gene TSS (-5/+25kb) in *Ctrl* SgU (n=2) for each expression group.

D) Line plots showing average H3K36me3 occupancy at Groups 1-6 hypoDMRs (±5kb) in *Ctrl* SgU (n=2).

E) Heatmaps showing H3K36me3 occupancy at center of hypoDMRs (±5kb) belonging to Groups 1-6 in *Ctrl* and *DKO* SgU and SgD (n=2 for each sample). Colored lines between heatmaps denote hypoDMRs with significantly differential H3K36me3 occupancy in the indicated contrasts. The GC percentage and CpG density at hypoDMRs (±5 kb) are indicated.

F) Line plots showing average H3K36me3 occupancy at Groups 1-6 hypoDMRs (±5kb) in *Ctrl* and *DKO* SgU and SgD (n=2 for each sample).

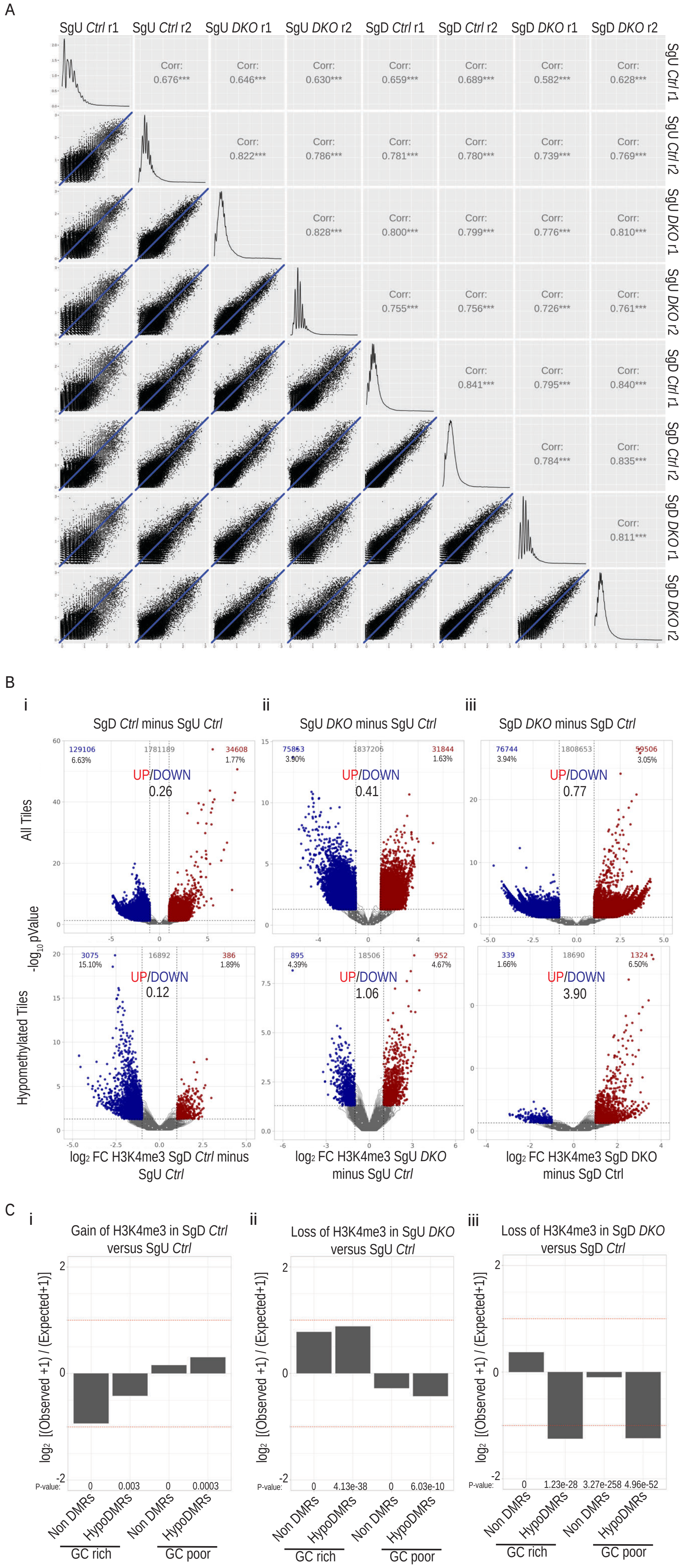
A
SgU Ctrl r1
SgU Ctrl r2
SgU DKO r1
SgU DKO r2
SgD Ctrl r1
SgD Ctrl r2
SgD DKO r1
SgD DKO r2
Corr: 0.676***
Corr: 0.646***
Corr: 0.630***
Corr: 0.659***
Corr: 0.689***
Corr: 0.582***
Corr: 0.628***
Corr: 0.822***
Corr: 0.786***
Corr: 0.781***
Corr: 0.780***
Corr: 0.739***
Corr: 0.769***
Corr: 0.828***
Corr: 0.800***
Corr: 0.799***
Corr: 0.776***
Corr: 0.810***
Corr: 0.755***
Corr: 0.756***
Corr: 0.726***
Corr: 0.761***
Corr: 0.841***
Corr: 0.795***
Corr: 0.840***
Corr: 0.784***
Corr: 0.835***
Corr: 0.811***
B
i
SgD Ctrl minus SgU Ctrl
ii
SgU DKO minus SgU Ctrl
iii
SgD DKO minus SgD Ctrl
All Tiles
-log10 pValue
Hypomethylated Tiles
129106 6.63%
1781189
34608 1.77%
UP/DOWN 0.26
75863 3.90%
1837206
31844 1.63%
UP/DOWN 0.41
76744 3.94%
1808653
59506 3.05%
UP/DOWN 0.77
3075 15.10%
16892
386 1.89%
UP/DOWN 0.12
895 4.39%
18506
952 4.67%
UP/DOWN 1.06
339 1.66%
18690
1324 6.50%
UP/DOWN 3.90
log2 FC H3K4me3 SgD Ctrl minus SgU Ctrl
log2 FC H3K4me3 SgU DKO minus SgU Ctrl
log2 FC H3K4me3 SgD DKO minus SgD Ctrl
C
i
Gain of H3K4me3 in SgD Ctrl versus SgU Ctrl
ii
Loss of H3K4me3 in SgU DKO versus SgU Ctrl
iii
Loss of H3K4me3 in SgD DKO versus SgD Ctrl
log2 [(Observed +1) / (Expected+1)]
P-value: 0 0.003 0 0.0003
P-value: 0 4.13e-38 0 6.03e-10
P-value: 0 1.23e-28 3.27e-258 4.96e-52
Non DMRs
HypoDMRs
GC rich
GC poor

**Supplementary Figure 9. Differential H3K4me3 analysis in *DKO* spermatogonia.**

A) Scatterplots [left side] show the H3K4me3 log2 RPKM values at 100.000 randomly selected genomic tiles detected by ULI-NChIP-seq between pairs of biological replicated samples of undifferentiated spermatogonia (SgU), differentiated spermatogonia (SgD) from *Ctrl* and *DKO* animals. The diagonal line plots show the distribution of the H3K4me3 log2 RPKM values in each replicate. The Pearson correlation coefficient [right side] of H3K4me3 log2 RPKM values at 100.000 randomly selected genomic tiles detected by ULI-NChIP-seq between the biological replicates of undifferentiated spermatogonia (SgU), differentiated spermatogonia (SgD) from *Ctrl* and *DKO* samples.

B) Volcano plots showing differential enrichment for H3K4me3 in (i) *Ctrl* SgD (n=2) versus *Ctrl* SgU (n=2), (ii) *DKO* SgU (n=2) versus *Ctrl* SgU (n=2) and (iii) *DKO* SgD (n=2) versus *Ctrl* SgD (n=2) (right) contrasts for all genomic tiles (upper) and hypoDMR tiles (lower). Tiles with statistically significant gain and loss of the H3K4me3 signals are highlighted in red (upregulated) and blue (downregulated) and their numbers and percentage of differentially are shown on the upper right and left corners respectively. The ratio between upregulated versus downregulated tiles in each plot is shown in the middle.

C) Bar plots showing log2 enrichments of GC-rich or GC-poor nonDMR and hypoDMR tiles among tiles with (i) significant H3K4me3 gain in *Ctrl* SgD versus *Ctrl* SgU, (ii) significant H3K4me3 loss in *DKO* SgU versus *Ctrl* SgU and (iii) significant H3K4me3 loss in *DKO* SgD versus *Ctrl* SgD. P-values were calculated by Fisher's exact test.

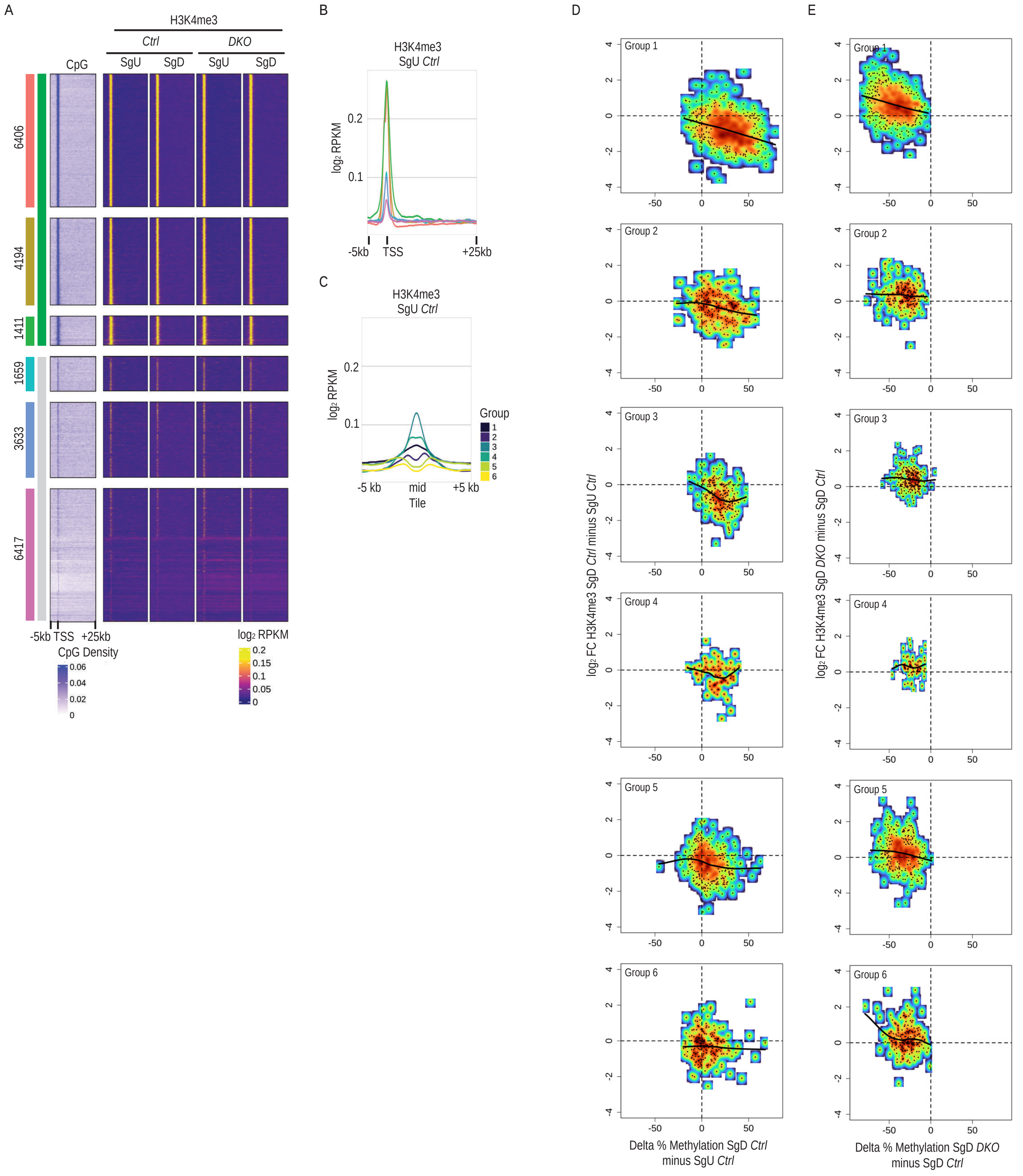
A
H3K4me3
Ctrl
DKO
CpG
SgU
SgD
SgU
SgD
6406
4194
1411
1659
3633
6417
-5kb TSS +25kb
CpG Density
0.06
0.04
0.02
0
log2 RPKM
0.2
0.15
0.1
0.05
0
B
H3K4me3
SgU Ctrl
log2 RPKM
-5kb
TSS
+25kb
C
H3K4me3
SgU Ctrl
Group
-5 kb
mid
+5 kb
Tile
D
E
Group 1
Group 2
Group 3
Group 4
Group 5
Group 6
log2 FC H3K4me3 SgD Ctrl minus SgU Ctrl
log2 FC H3K4me3 SgD DKO minus SgD Ctrl
Delta % Methylation SgD Ctrl minus SgU Ctrl
Delta % Methylation SgD DKO minus SgD Ctrl

**Supplementary Figure 10. H3K4me3 levels of hypoDMRs in *DKO* spermatogonia.**

A) Heatmaps showing H3K4me3 occupancy at gene transcriptional start sites (TSS) (-5/+25kb) in *Ctrl* and *DKO* SgU and SgD (n=2 for each sample). The CpG density at gene TSS (-5/+25kb) is indicated. Genes have been grouped according to the presence of a CpG island at their TSS +/- 1kb and ordered by decreasing expression in *Ctrl* SgU.

B) Line plots showing average H3K4me3 occupancy at gene TSS (-5/+25kb) in *Ctrl* SgU (n=2) for each expression group.

C) Line plots showing average H3K4me3 occupancy at Groups 1-6 hypoDMRs (±5kb) in *Ctrl* SgU (n=2).

D) Scatterplots showing DNAme difference between *Ctrl* SgD minus *Ctrl* SgU (x-axis) versus log2 fold change of H3K4me3 signal between *Ctrl* SgD minus *Ctrl* SgU (y-axis) for hypoDMRs in Groups 1-6. Black lines represent LOESS weighted linearly regressed data.

E) Scatterplots showing DNAme difference between *DKO* SgD minus *Ctrl* SgD (x-axis) versus log2 fold change of H3K4me3 signal between *DKO* SgD minus *Ctrl* SgD (y-axis) for hypoDMRs in Groups 1-6. Black lines represent LOESS weighted linearly regressed data.

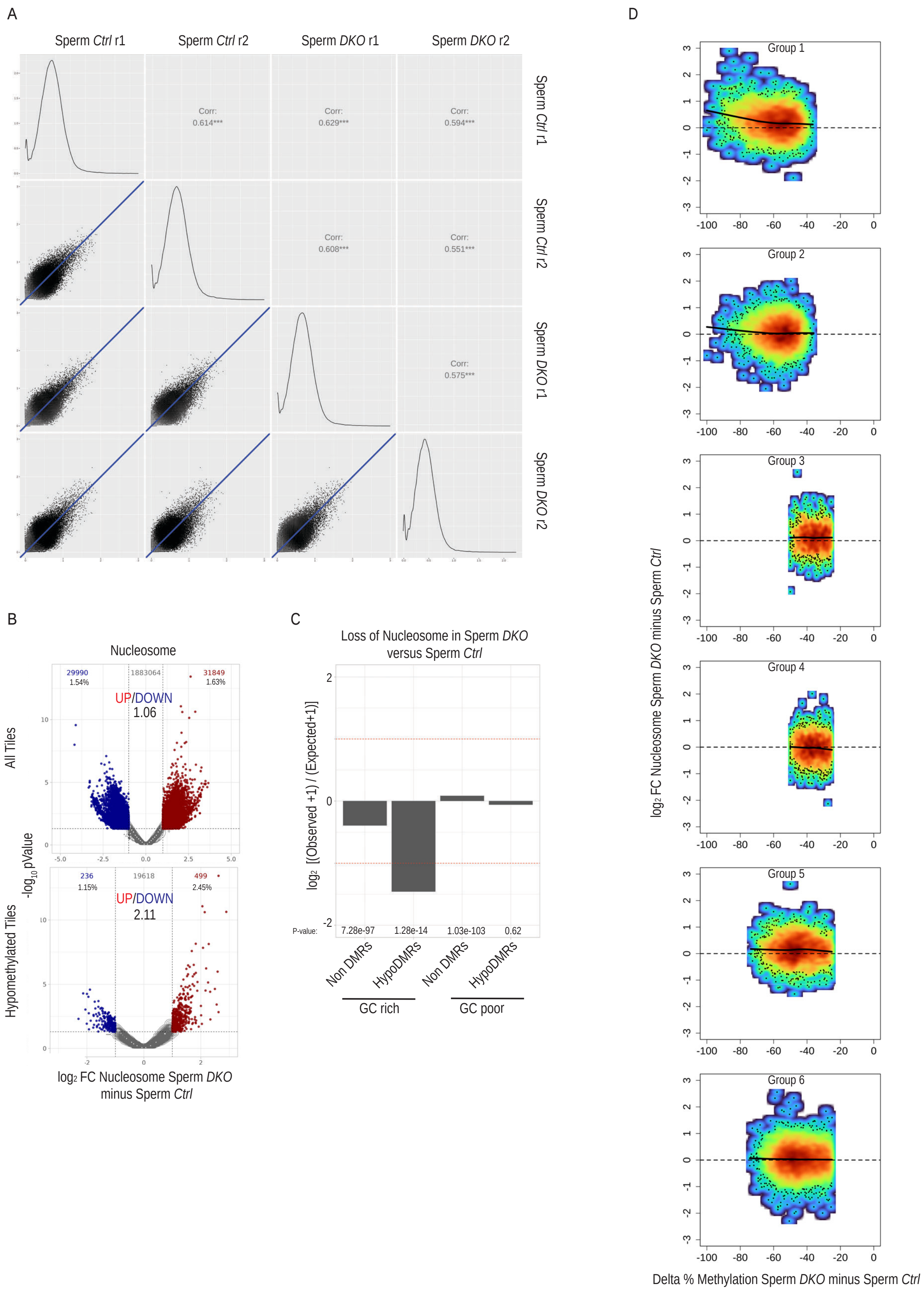
A
Sperm Ctrl r1
Sperm Ctrl r2
Sperm DKO r1
Sperm DKO r2
Corr: 0.614***
Corr: 0.629***
Corr: 0.594***
Corr: 0.608***
Corr: 0.551***
Corr: 0.575***
B
Nucleosome
All Tiles
29990
1.54%
1883064
31849
1.63%
UP/DOWN
1.06
Hypomethylated Tiles
236
1.15%
19618
499
2.45%
UP/DOWN
2.11
-log10 pValue
log2 FC Nucleosome Sperm DKO minus Sperm Ctrl
C
Loss of Nucleosome in Sperm DKO versus Sperm Ctrl
log2 [(Observed +1) / (Expected+1)]
P-value:
7.28e-97
1.28e-14
1.03e-103
0.62
Non DMRs
HypoDMRs
Non DMRs
HypoDMRs
GC rich
GC poor
D
Group 1
Group 2
Group 3
Group 4
Group 5
Group 6
log2 FC Nucleosome Sperm DKO minus Sperm Ctrl
Delta % Methylation Sperm DKO minus Sperm Ctrl

**Supplementary Figure 11. Differential nucleosome analysis in *DKO* sperm.**

A) Scatterplots [left side] show the nucleosome log2 RPKM values at 100.000 randomly selected genomic tiles detected by ULI-NChIP-seq between pairs of biological replicated samples of sperm from *Ctrl* and *DKO* animals. The diagonal line plots show the distribution of the nucleosome log2 RPKM values in each replicate. The Pearson correlation coefficient [right side] of nucleosome log2 RPKM values at 100.000 randomly selected genomic tiles detected by ULI-NChIP-seq between the biological replicates of sperm from *Ctrl* and *DKO* samples.

B) Volcano plots showing differential enrichment for nucleosome in *DKO* (n=2) versus *Ctrl* (n=2) sperm contrast for all genomic tiles (upper) and hypoDMR tiles (lower). Tiles with statistically significant gain and loss of the nucleosomal signals are highlighted in red (upregulated) and blue (downregulated) and their numbers and percentage of differentially are shown on the upper right and left corners respectively. The ratio between upregulated versus downregulated tiles in each plot is shown in the middle.

C) Bar plots showing log2 enrichments of GC-rich or GC-poor nonDMR and hypoDMR tiles among tiles with significant nucleosome loss in *DKO* sperm versus *Ctrl*. P-values were calculated by Fisher's exact test.

D) Scatterplots showing DNAme difference between *DKO* minus *Ctrl* sperm (x-axis) versus log2 fold change of nucleosome signal between *DKO* minus *Ctrl* sperm (y-axis) for hypoDMRs in Groups 1-6. Black lines represent LOESS weighted linearly regressed data.

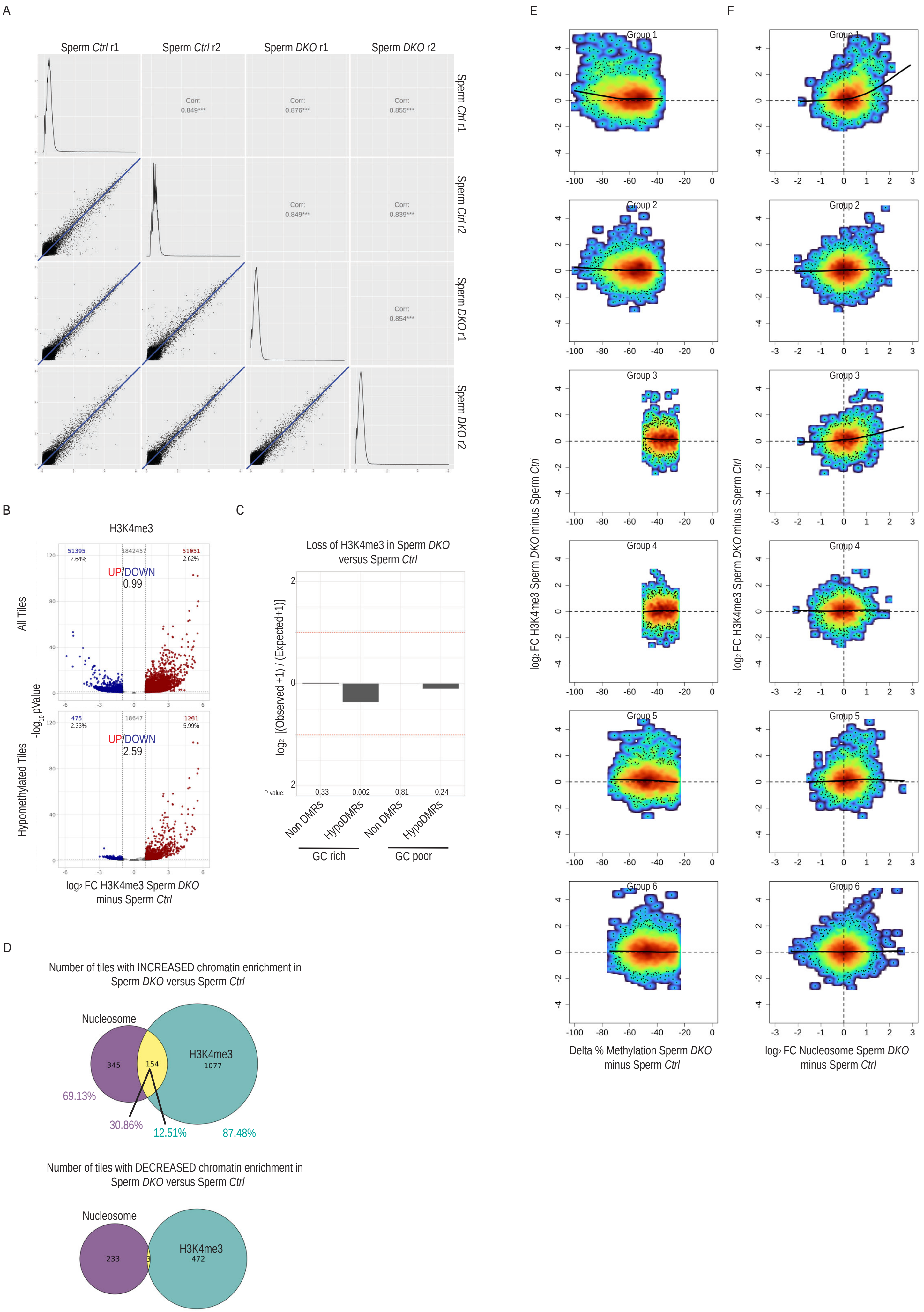
A
Sperm Ctrl r1
Sperm Ctrl r2
Sperm DKO r1
Sperm DKO r2
Corr: 0.849***
Corr: 0.876***
Corr: 0.855***
Corr: 0.849***
Corr: 0.839***
Corr: 0.854***
B
H3K4me3
All Tiles
51395 2.64%
1842457
51851 2.62%
UP/DOWN 0.99
Hypomethylated Tiles
475 2.33%
18647
1231 5.99%
UP/DOWN 2.59
-log10 pValue
log2 FC H3K4me3 Sperm DKO minus Sperm Ctrl
C
Loss of H3K4me3 in Sperm DKO versus Sperm Ctrl
log2 [(Observed +1) / (Expected+1)]
P-value: 0.33 0.002 0.81 0.24
Non DMRs
HypoDMRs
Non DMRs
HypoDMRs
GC rich
GC poor
D
Number of tiles with INCREASED chromatin enrichment in Sperm DKO versus Sperm Ctrl
Nucleosome
H3K4me3
345
154
1077
69.13%
30.86%
12.51%
87.48%
Number of tiles with DECREASED chromatin enrichment in Sperm DKO versus Sperm Ctrl
Nucleosome
H3K4me3
233
3
472
E
F
Group 1
Group 2
Group 3
Group 4
Group 5
Group 6
log2 FC H3K4me3 Sperm DKO minus Sperm Ctrl
Delta % Methylation Sperm DKO minus Sperm Ctrl
log2 FC Nucleosome Sperm DKO minus Sperm Ctrl

**Supplementary Figure 12. Differential H3K4me3 analysis in *DKO* sperm.**

A) Scatterplots [left side] show the H3K4me3 log2 RPKM values at 100.000 randomly selected genomic tiles detected by ULI-NChIP-seq between pairs of biological replicated samples of sperm from *Ctrl* and *DKO* animals. The diagonal line plots show the distribution of the H3K4me3 log2 RPKM values in each replicate. The Pearson correlation coefficient [right side] of H3K4me3 log2 RPKM values at 100.000 randomly selected genomic tiles detected by ULI-NChIP-seq between the biological replicates of sperm from *Ctrl* and *DKO* samples.

B) Volcano plots showing differential enrichment for H3K4me3 in *DKO* (n=2) versus *Ctrl* (n=2) sperm contrast for all genomic tiles (upper) and hypoDMR tiles (lower). Tiles with statistically significant gain and loss of the nucleosomal signals are highlighted in red (upregulated) and blue (downregulated) and their numbers and percentage of differentially are shown on the upper right and left corners respectively. The ratio between upregulated versus downregulated tiles in each plot is shown in the middle.

C) Bar plots showing log2 enrichments of GC-rich or GC-poor nonDMR and hypoDMR tiles among tiles with significant H3K4me3 loss in *DKO* sperm versus *Ctrl*. P-values were calculated by Fisher's exact test.

D) Euler diagrams representing the number of hypoDMRs that are differentially gain enrichment in *DKO* versus *Ctrl* sperm either for nucleosome or H3K4me3 or both (left) and the number of hypoDMRs that are differentially lose enrichment in *DKO* versus *Ctrl* sperm either for nucleosome or H3K4me3 or both (right).

E) Scatterplots showing DNAme difference between *DKO* minus *Ctrl* sperm (x-axis) versus log2 fold change of H3K4me3 signal between *DKO* minus *Ctrl* sperm (y-axis) for hypoDMRs in Groups 1-6. Black lines represent LOESS weighted linearly regressed data.

E) Scatterplots showing log2 fold change of nucleosome signal between *DKO* minus *Ctrl* sperm (x-axis) versus log2 fold change of H3K4me3 signal between *DKO* minus *Ctrl* sperm (y-axis) for hypoDMRs in Groups 1-6. Black lines represent LOESS weighted linearly regressed data.

Fanourgakis et al Supplementary Figure 13

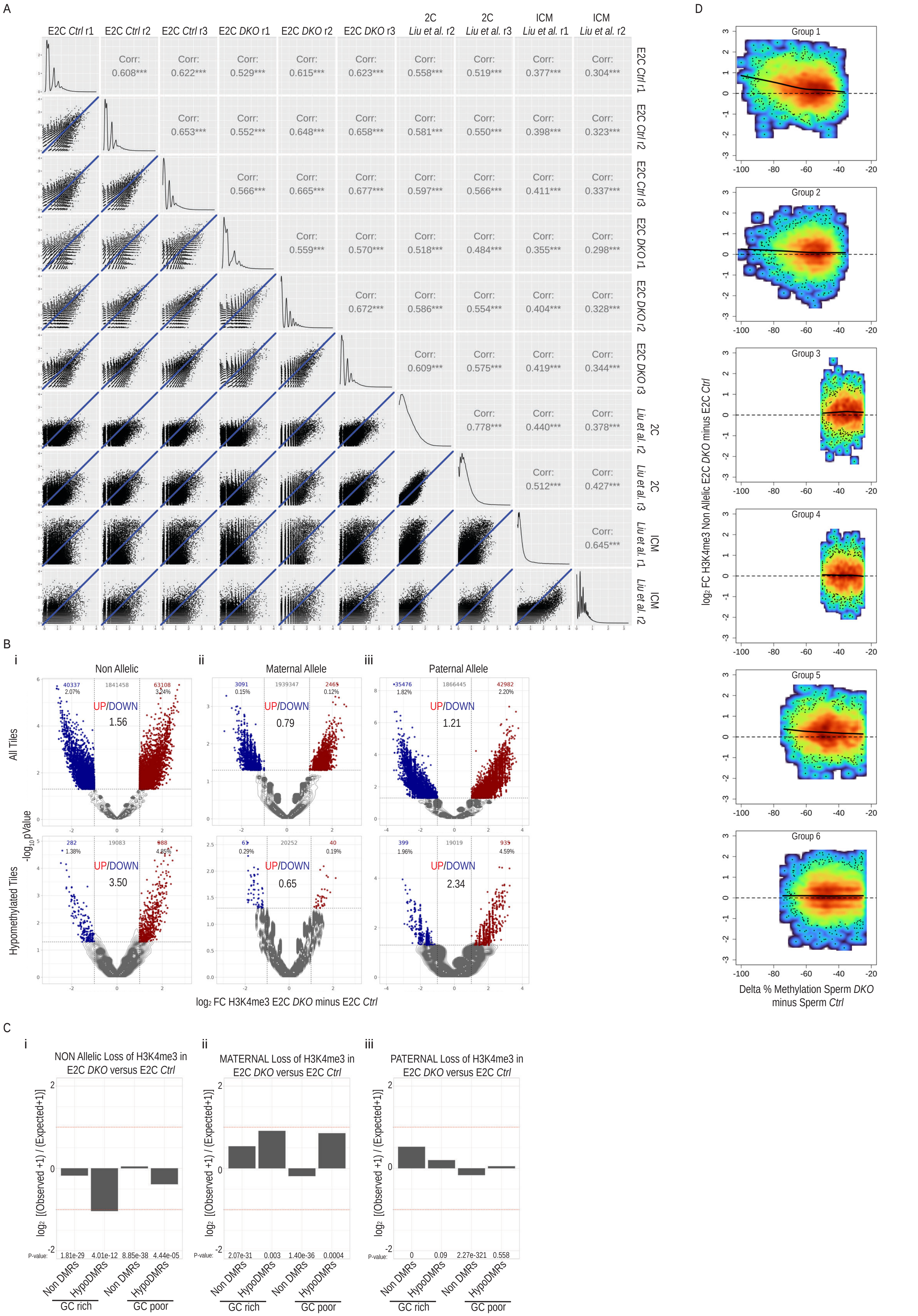

**Supplementary Figure 13. Differential H3K4me3 analysis in *DKO* sperm derived early 2 cell stage (E2C) embryos.**

A) Scatterplots [left side] show the H3K4me3 log2 RPKM values at 100.000 randomly selected genomic tiles detected by ATATA-seq between pairs of biological replicated samples of early 2-cell stage (E2C) embryos generated by *Ctrl* or *DKO* sperm and detected by CHIP-seq in wild type 2 cell stage embryos and blastocyst stage ICM samples (from Liu et al., 2016) as a reference. The diagonal line plots show the distribution of the H3K4me3 log2 RPKM values in each replicate. The Pearson correlation coefficient [right side] of H3K4me3 log2 RPKM values at 100.000 randomly selected genomic tiles detected by ATATA-seq between the biological replicated samples of early 2-cell stage (E2C) embryos generated by *Ctrl* or *DKO* sperm and detected by CHIP-seq in wild type 2 cell stage embryos and blastocyst stage ICM samples (from Liu et al., 2016) as a reference.

B) Volcano plots showing differential enrichment for H3K4me3 in *DKO* (n=3) versus *Ctrl* (n=3) sperm derived E2C embryos contrast (i) without allelic assignment, (ii) at the maternal allele or (iii) at the paternal allele for all genomic tiles (upper) and hypoDMR tiles (lower). Tiles with statistically significant gain and loss of the H3K4me3 signals are highlighted in red (upregulated) and blue (downregulated) and their numbers and percentage of differentially are shown on the upper right and left corners respectively. The ratio between upregulated versus downregulated tiles in each plot is shown in the middle.

C) Bar plots showing log2 enrichments of GC-rich or GC-poor nonDMR and hypoDMR tiles among tiles with significant H3K4me3 loss at (i) all alleles, (ii) maternal alleles and (iii) paternal alleles in early 2-cell (E2C) embryos generated by *Ctrl* or *DKO* sperm. P-values were calculated by Fisher's exact test.

D) Scatterplots showing DNAme difference between *DKO* minus *Ctrl* sperm (x-axis) versus log2 fold change of H3K4me3 signal between *DKO* minus *Ctrl* sperm derived early 2-cell (E2C) embryos (y-axis) for hypoDMRs in Groups 1-6. Black lines represent LOESS weighted linearly regressed data.

Fanourgakis et al Supplementary Figure 14

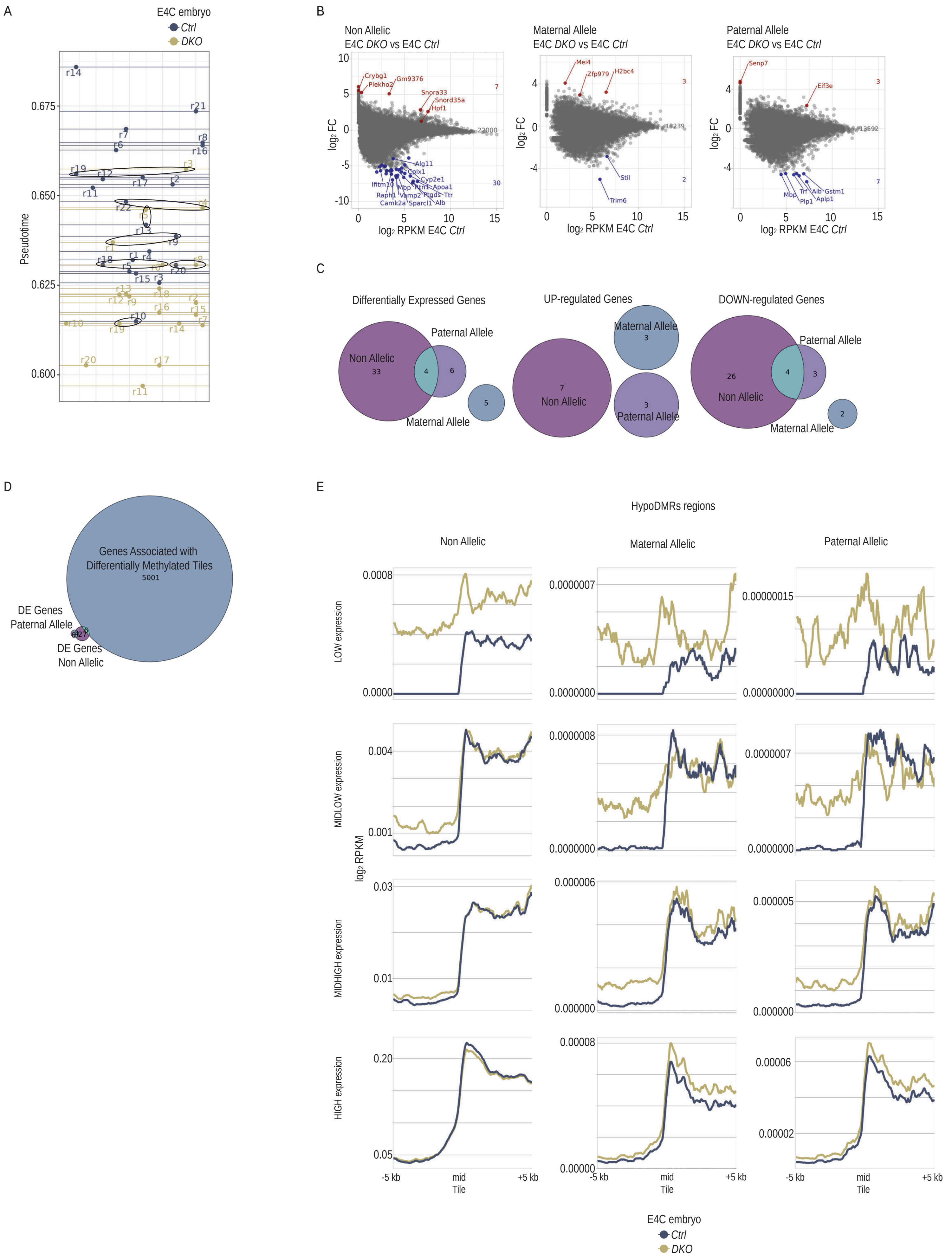

**Supplementary Figure 14. Transcriptional effects of Dnmt3a/Dnmt3b *DKO* and *Ctrl* sperm derived embryos.**

A) Dotplot shows the assigned developmental pseudotime of each embryo (n= 21 for *Ctrl* and n= 20 for *DKO* sperm derived embryos). Circles denote paired samples that were used for downstream differential gene expression analysis (n= 7 for *Ctrl* and n= 7 for *DKO* sperm derived embryos).

B) Scatter plots showing gene expression log2 fold changes among *DKO* (n=7) versus *Ctrl* (n=7) sperm derived 4-cell stage embryos samples were plotted against average gene expression levels in *Ctrl* sperm derived 4-cell stage embryos samples. From left to right: using all reads regardless parental origin, reads from maternal allele, reads from paternal allele. The numbers of significantly Upregulated (red) or Downregulated (blue) genes are displayed on the left and genes names of the most highly significant differentially expressed genes are shown.

C) Euler diagrams representing the number of differentially expressed genes from (B) for non-allelic, maternal, or paternal allele specific analysis (left), the number of upregulated genes from (B) for non-allelic, maternal or paternal allele specific analysis (middle) and the number of downregulated genes from (B) for non-allelic, maternal or paternal allele specific analysis (right).

D) Euler diagram representing the number of differentially expressed genes from (B) for non-allelic, maternal, or paternal allele specific analysis and the number of hypoDMR regions associated genes.

E) Line plots showing average RNA occupancy at the center of hypoDMR regions (±5 kb) in *DKO* versus *Ctrl* sperm derived 4 cell stage embryos samples. The hypoDMR regions are oriented to display high occupancy on the positive direction and stratified into 4 quantile expression groups (low, midlow, midhigh and high) based in the *Ctrl* sperm derived 4 cell stage embryos. From left to right: using all reads regardless parental origin, reads from maternal allele, reads from paternal allele.

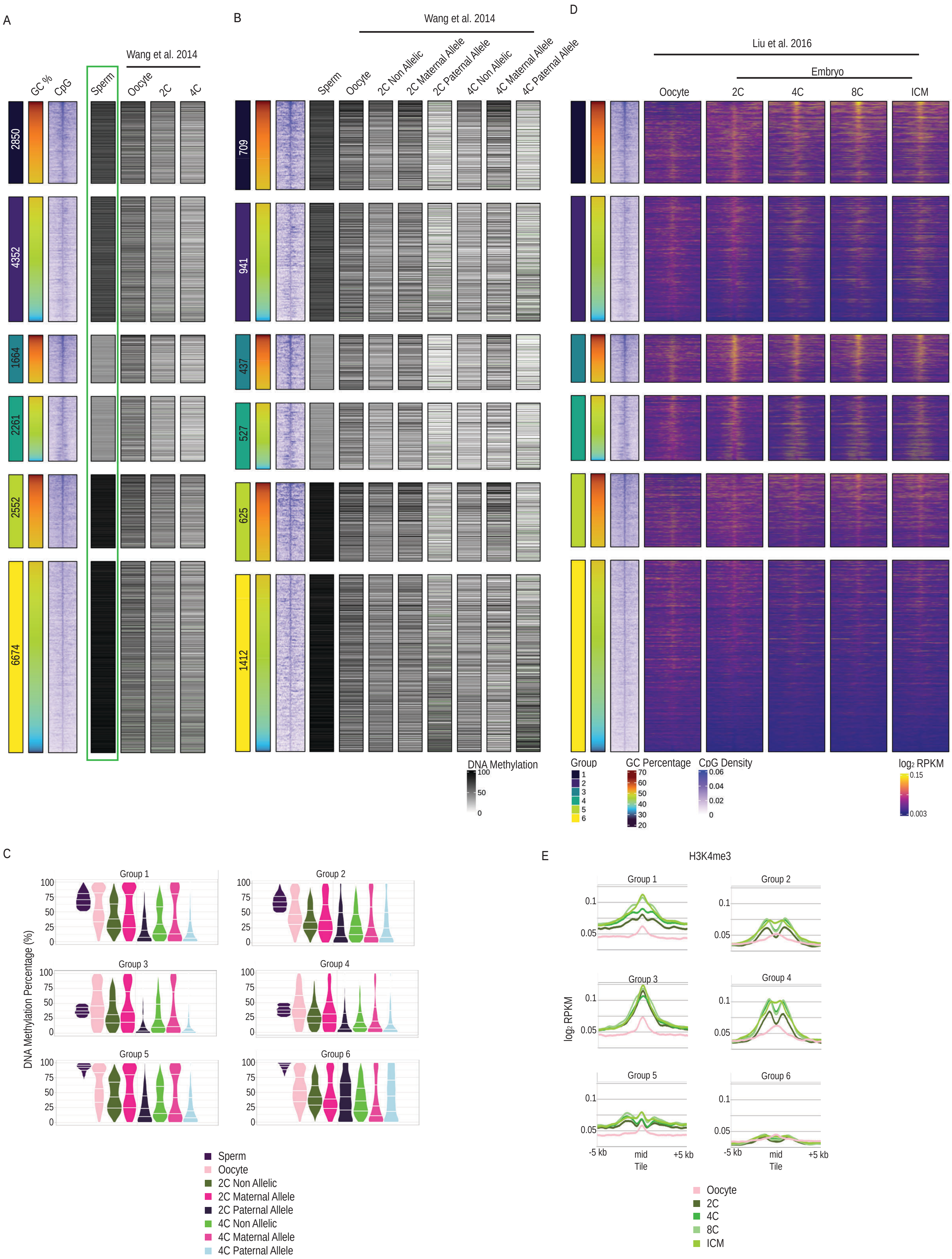
A
GC %
CpG
Sperm
Wang et al. 2014
Oocyte
2C
4C
2850
4352
1664
2261
2552
6674
B
Wang et al. 2014
Sperm
Oocyte
2C Non Allelic
2C Maternal Allele
2C Paternal Allele
4C Non Allelic
4C Maternal Allele
4C Paternal Allele
709
941
437
527
625
1412
DNA Methylation
100
50
0
D
Liu et al. 2016
Embryo
Oocyte
2C
4C
8C
ICM
Group
1
2
3
4
5
6
GC Percentage
70
60
50
40
30
20
CpG Density
0.06
0.04
0.02
0
log2 RPKM
0.15
0.003
C
DNA Methylation Percentage (%)
Group 1
Group 2
Group 3
Group 4
Group 5
Group 6
100
75
50
25
0
Sperm
Oocyte
2C Non Allelic
2C Maternal Allele
2C Paternal Allele
4C Non Allelic
4C Maternal Allele
4C Paternal Allele
E
H3K4me3
log2 RPKM
Group 1
Group 2
Group 3
Group 4
Group 5
Group 6
0.1
0.05
-5 kb
mid
+5 kb
Tile
Oocyte
2C
4C
8C
ICM

**Supplementary Figure 15. DNAme and H3K4me3 dynamics of hypoDMRs in *Ctrl* early embryogenesis.**

A) Heatmap showing the DNAme percentage of hypoDMRs in *Ctrl* sperm (this study), wild type oocytes, 2-cell stage and 4 cell stage embryos (from Wang et al., 2014). The density of CpGs at hypoDMRs (±5 kb from their center) is indicated. Within each group, hypoDMRs are ordered according to decreasing GC percentages. The panel in the green box is presented also in Figure 1E.

B) Heatmap showing the DNAme percentage as in (A), but for the subset of hypoDMRs that contain enough informative SNPs to allow parental allelic assignment. From left to right: *Ctrl* sperm (this study), wild type oocytes, 2-cell stage without genome assignment, 2-cell stage maternal allele, 2-cell stage paternal allele, 4-cell stage without genome assignment, 4-cell stage maternal allele and 4-cell stage paternal allele (from Wang et al., 2014). The density of CpGs at hypoDMRs (±5 kb from their center) is indicated. Within each group, hypoDMRs are ordered according to decreasing GC percentages.

C) Violin plots showing the percentage of DNAme of hypoDMRs in *Ctrl* sperm (this study), wild type oocytes, 2-cell stage without genome assignment, 2-cell stage maternal allele, 2-cell stage paternal allele, 4-cell stage without genome assignment, 4-cell stage maternal allele and 4-cell stage paternal allele (from Wang et al., 2014) for each group. The center lines in the violins represent median values. The upper and lower lines indicate the interquartile range (IQR; from the 25th to 75th percentile).

D) Heatmaps showing H3K4me3 occupancy at center of Group 1 to 6 hypoDMRs (±5kb) in wild type oocytes, 2-cell stage embryos, 4-cell stage embryos, 8-cell stage embryos and ICM blastocyst stage embryos (from Liu et al., 2016). The density of CpGs at hypoDMRs (±5 kb from their center) is indicated. Within each group, hypoDMRs are ordered according to decreasing GC percentages.

E) Line plots showing average H3K4me3 occupancy at Groups 1-6 hypoDMRs (±5kb) in wild type oocytes, 2-cell stage embryos, 4-cell stage embryos, 8-cell stage embryos and ICM blastocyst stage embryos (from Liu et al., 2016).
